# Oxidative stress-induced proteolytic activation of polyphenol oxidase triggers an oxidized flavonoids-mediated stress signaling in *Camellia sinensis*

**DOI:** 10.1101/2025.09.23.677533

**Authors:** Sumanta Mohapatra, Anand Mishra, Shagun Bali, Ritu Godara, Twinkle, Anil Kumar, Ravi Kumar, Neeraj Kumar, Pawan Kumar, Vishal Acharya, Vivek Dogra

**Author notes:** Author for correspondence: Vivek Dogra.

## Abstract

Environmental perturbations often increase reactive oxygen species (ROS) production, inducing oxidative damage to various biomolecules. The cellular system utilizes ROS or ROS-generated metabolites to instigate signaling pathways, resulting in acclimation, growth inhibition, or programmed cell death (PCD) for sustaining stress. Various stress-generated or stress-activated signaling components and downstream pathways are identified in model plants; however, they are largely unexplored in non-model plants. Here, we report a stress-induced proteolytic activation of an evolutionarily conserved chloroplast-localized enzyme, polyphenol oxidase (PPO), that oxidizes catechins into theaflavins (TFs) and initiates a stress signaling under drought in *Camellia sinensis* (Tea). Germplasm-based analysis revealed a heightened proteolytic activation of PPO and consequent TFs accumulation in drought-susceptible genotypes. Transcriptome analysis revealed that PPO activation and TFs accumulation were linked to the activation of UPR(unfolded protein response)-like response, which was reinforced by virus-induced silencing and overexpression of PPO, and direct feeding of TFs in tea plants. Pharmacological treatments revealed that TFs interact with HSP90, activating a canonical IRE-bZIP60-dependent ER stress pathway resulting in PCD. Similar proteolytic activation of PPO and subsequent instigation of stress signaling in other plant species (tomato and wheat) demonstrated that PPO acts as an evolutionarily conserved stress sensor, instigating an inter-organelle communication in plants.

**Significance statement:** PPO, an evolutionarily conserved and chloroplast/plastid localized enzyme, senses stress and undergoes proteolytic activation to generate oxidized flavonoids/quinones that function as signaling molecules in Tea and likely in other plants. The oxidized flavonoids modulate protein folding machinery and augment folding stress, especially in the ER, inducing canonical IRE1-bZI60-dependent PCD. The finding shows that specific secondary metabolites likely have roles in instigating intra- and intercellular communication to sustain the adversities of stress in different plant species.

## Introduction

Plants are sessile and constantly affected by environmental perturbations, which affect their growth and development. Environmental cues such as high light (HL), drought, salinity, cold/heat, and pathogens often increase the production of reactive oxygen species (ROS) in different cell organelles, resulting in oxidative damage to various molecules, including proteins ^1, 2, 3^. Oxidative damage impairs folding and alters protein structure, leading to folding stress marked with accumulated damaged/misfolded proteins in sub-cellular compartments, mainly ER, cytosol, chloroplast, and mitochondria ^4, 5, 6, 7^. The cellular system then activates a folding stress response (FSR), upregulating protein quality control (PQC) machinery to restore proteostasis under mild stress. Upon severe but sub-lethal stress conditions, the FSR involves the activation of a controlled and programmed cell death (PCD) pathway for sustaining stress ^5, 8^. Such PCD does not impair the viability of the affected plant but rather provides resilience ^2, 9, 10, 11, 12^. As a large proportion of proteins are targeted co-translationally through the endomembrane system, unfolded protein accumulation was first detected in the ER, creating ER stress, and a response thus activated to restore proteostasis was referred to as the ER stress response or unfolded protein response (UPR). Lately, unfolded or damaged proteins have been discovered to accumulate in cytosol and organelles, where the generated responses are referred to as cytosol-specific UPR, which sometimes is also referred to as heat stress responses (HSR), and organelle-specific UPRs (Cp-UPR and Mt-UPR)^4, 5, 6, 7, 13^. These UPRs are now collectively called FSR.

In addition to causing oxidative damage, ROS also serves as the potent signaling molecules that regulate plant growth and development and activate responses toward stress stimuli ^14^. Among various subcellular organelles, chloroplasts are the primary sites of ROS generation and oxidative damage; thus, they are eminently sensitive to environmental cues ^15, 16^. The ROS, either by themselves (especially H_2_O_2_), by generating secondary reactive molecules (produced from the oxidation of lipids and carotenoids, oxidative inhibition or activation of metabolic enzymes or proteins, accumulating specific secondary metabolites), activate retrograde signaling cascades to alter nuclear gene expression ^17, 18, 19, 20, 21, 22^. Upon meeting such signals, plants may suspend their growth in varying degrees or activate PCD depending upon the stress severity ^2, 9, 10, 11, 12^. Among various biomolecules, proteins often act as sensors of environmental stimuli and thereby undergo post-translational modifications, including proteolysis, to activate stress signaling through substrate or product molecules. This proteolytic mediated activated response led to initiate numerous signalling in plants. For instance, during ER stress, the membrane-tethered transcription factor bZIP28 is cleaved by Site-1 and Site-2 proteases (S1P and S2P), liberating a soluble fragment that translocates to the nucleus to activate the UPR ^23^. Similarly, cold stress induces the proteolytic processing of the membrane-associated NAC transcription factor NTL6, facilitating its nuclear localization and the upregulation of cold-responsive genes^24^. In systemic defense, mechanical wounding and herbivory trigger the protease-mediated conversion of prosystemin into active systemin, a mobile peptide that orchestrates jasmonate-dependent defense signaling across the plant ^25^. Furthermore, drought stress activates specific subtilisin-like proteases that cleave prophytosulfokine (ProPSK) precursors to generate bioactive peptides that modulate growth and enhance resilience ^26^. The ROS sensor proteins such as EXECUTER 1 (EX1) and EXECUTER 2 (EX2) in chloroplasts, undergo a ROS-induced proteolysis, initiating a signaling pathway leading to growth inhibition and PCD ^2, 27^. These diverse cases illustrate that proteolysis functions as a conserved regulatory mechanism in plant stress signaling, rapidly mobilizing pre-existing proteins into active signaling agents. Various stress-generated signaling molecules and downstream pathways are identified in model plants like *Arabidopsis thaliana*; however, they remain untapped in non-model plants, especially those accumulating a variety of secondary metabolites implicated with stress management.

*Camellia sinensis* (Tea), a non-deciduous evergreen perennial shrub, is the most widely consumed beverage due to its specialized secondary metabolite accumulation ^28, 29^. Tea plants often experience environmental perturbations during their growth and development, including drought, temperature fluctuations, salinity, high pH, and heavy metals in soil. Among these stresses, drought is more prevalent and is reported to affect quality, reduce yield, and increase plant mortality ^30, 31, 32^. To sustain drought, plants exhibit reduced transpirational water loss by stomatal closure and adequate water absorption through deeper roots with elongated root hairs ^33^. However, stomatal closure reduces photosynthetic output, inhibiting growth ^33, 34^. In addition, drought also results in increased ROS generation, mainly in chloroplasts. A range of primary (enzymatic) and secondary (non-enzymatic) antioxidant systems eliminate excess ROS to maintain normal cell metabolism ^3, 35^. The secondary non-enzymatic antioxidants comprise mainly glutathione, ascorbic acid, carotenoids, and polyphenols ^3, 36, 37^. Flavonoids are a class of polyphenols that also act as antioxidants, mainly through hydrogen and electron supply, to scavenge ROS ^36, 37^. Catechins are the primary flavonoid antioxidants, accounting for more than 70% of total polyphenols in Tea plants ^38^. They are synthesized through phenylpropanoid and flavonoid pathways ^39^. Catechins undergo chemical modifications during Tea processing (withering, rolling, fermentation, and drying), affecting taste, liquor color, and aroma quality ^40, 41^. Catechins can be enzymatically oxidized by polyphenol oxidase (PPO) to *o*-quinones, which further undergo oxidation and polymerization, forming four different brown-colored pigments, theaflavins (TFs), which constitute unique quality metabolites in black Tea ^42^. TFs possess ROS scavenging, cytotoxic, and anti-inflammatory properties ^41^. TFs also undergo oxidation, forming thearubigins (TRs), which also possess antioxidation and anticancer properties. Abiotic stresses such as drought have been thought to improve black Tea quality by increasing PPO activity and, thereby, modulating the contents of catechins, TFs, and TRs ^31^. Though these metabolites have been investigated for post-harvest Tea quality, their biological function in plants is largely unexplored.

PPOs are intracellular copper-containing metalloproteins present in bacteria, fungi, and plants that catalyze the *o*-hydroxylation of monophenols to *o*-diphenols (tyrosinase activity) as well as the oxidation of *o*-diphenols to quinones (catecholase activity). In plants, PPO protein contains N-terminal transit peptides, which direct it into the thylakoid lumen, where it remains latent ^43, 44^. Upon activation, it converts its substrates into products. Although the phenolic substrates are generally stored in the vacuole, catechins, notably, also accumulate in chloroplasts after being synthesized in the cytosol ^45^. Due to its localization in thylakoid, PPO has been implicated in photosynthesis ^46^; however, this notion lacks concrete evidence. PPO is present in several plant species, except a few plants including *Arabidpsi thaliana*, as a multigene family, where *PPO* genes express differentially in plant parts and respond to senescence, wounding, and biotic elicitors. Tea genome contains 4 *PPO* genes where *CsPPO1* and *CsPPO2* are induced by wounding and moth infestation ^47^. Methyl jasmonate enhances the expression of *CsPPO2* and *CsPPO4* to improve defense against *Ectropis grisescens* larvae in Tea plants ^48^, through canonical R2R3 transcription factor MYB59 ^49^. Though PPO has been experimentally implicated with biotic stress tolerance, its role in abiotic stress remains inconclusive. In fact, it is debatable whether PPO activity is beneficial or detrimental to the plant ^50^. Under severe stress, plants instigate PCD to sacrifice weaker/damaged cells to sustain energy and nutritional requirements as a survival mechanism ^5^. Interestingly, PPO has been implicated with PCD, where silencing of *PPO* in walnut plants resulted in the accumulation of its substrate tyramine, which is known to elicit cell death in different plant species ^51^. Tea plants also exhibit growth inhibition and PCD under drought stress with an increased TFs content ^30^, suggesting a possible role of PPO and TFs in regulating stress responses. Given the increased levels of phenolic compounds and heightened expression of PPO under drought ^30^, it is likely that the oxidized catechins, namely TFs and TRs, play a role in determining stress response in Tea plants. Unfortunately, despite several studies, the biological relevance of PPO and its catalyzed products in Tea plants remain unelucidated.

Considering this, we hypothesize that PPO might act as a stress sensor, activating TFs-mediated signaling to modulate nuclear gene expression and activate stress responses, including PCD. To prove our hypothesis, we analyzed a set of Tea genotypes exhibiting varying degrees of drought resilience/susceptibility, where susceptible genotypes showed higher ROS accumulation, oxidative damage, and cell death, accompanied by higher activity of PPO and accumulation of TFs. The chloroplast-localized PPO was found to undergo stress-induced proteolytic activation, leading to the conversion of its substrate catechins into TFs in chloroplasts. Contrasting genotypes-based analysis revealed a heightened proteolytic activation of PPO and consequent TFs accumulation in drought-susceptible genotypes. Transcriptome analysis revealed that PPO activation and TFs accumulation were linked to the activation of UPR(unfolded protein response)-like response, which was reinforced by virus-induced silencing and overexpression of PPO, and direct feeding of TFs in tea plants. Pharmacological treatments revealed that TFs interact with HSP90, activating a canonical IRE-bZIP60-dependent ER stress pathway resulting in PCD. Similar proteolytic activation of PPO and subsequent instigation of stress signaling in other plant species (tomato and wheat) demonstrated that PPO acts as an evolutionarily conserved stress sensor, instigating an inter-organelle communication in plants.

## Results

### Drought-induced oxidative stress results in increased flavonoid content, PPO activity and cell death in Tea

To understand the impact of drought on plant performance and flavonoid accumulation, we analyzed a set of 12 Tea genotypes subjected to PEG-induced drought. To ascertain the impact of drought, we analysed chlorophyll fluorescence, relative water content (RWC), oxidative damage through lipid peroxidation as represented by malondialdehyde (MDA) content, and relative electrolyte leakage (REL). The impact of drought on all the genotypes was apparent within 24 h of stress (Fig. 1a-b; Supplementary Fig. 1a). With increasing stress, the leaves showed chlorosis and browning, coinciding with a decline in moisture contents (Fig. 1a and c, and Supplementary Fig. 1a). The chlorosis led to reduced chlorophyll fluorescence and PSII activity (Fig. 1b), marking the impact of photooxidative stress on chloroplast, an early sensor of drought. The photooxidative stress resulted in lipid peroxidation and enhanced membrane damage, leading to cell death (Fig. 1d-e). Interestingly, the selected genotypes under study exhibited varying responses to stress; although all were sensitive to drought, a few were highly susceptible (TV19, TV03, TV01, SA06, TV18, and T383), whereas others were comparatively resilient (TG01, TV17, UPASI09, B157, HV and TEEN ALI). The susceptible genotypes showed rapid chlorosis and a sharp decline in chlorophyll fluorescence and RWC (Fig. 1a-c, Supplementary Fig. 1a). They experienced higher oxidative damage, resulting in cell death (Fig. 1d-e). On the contrary, the resilient genotypes exhibited somewhat delayed stress symptoms revealed by plant phenotype, lesser loss of chlorophyll fluorescence and RWC, and comparatively slighter lipid peroxidation and electrolyte leakage with a gradual increase in stress (Fig. 1a-e). Previously, a germplasm-based analysis was carried out to understand the impact of drought on two contrasting Tea genotypes, namely TV17 and TV03 ^29^. In agreement with earlier findings ^29^, TV17 performed better, and the impact of drought on TV03 was much apparent within 24 h of stress (Fig. 1a-e).

**Fig. 1:**
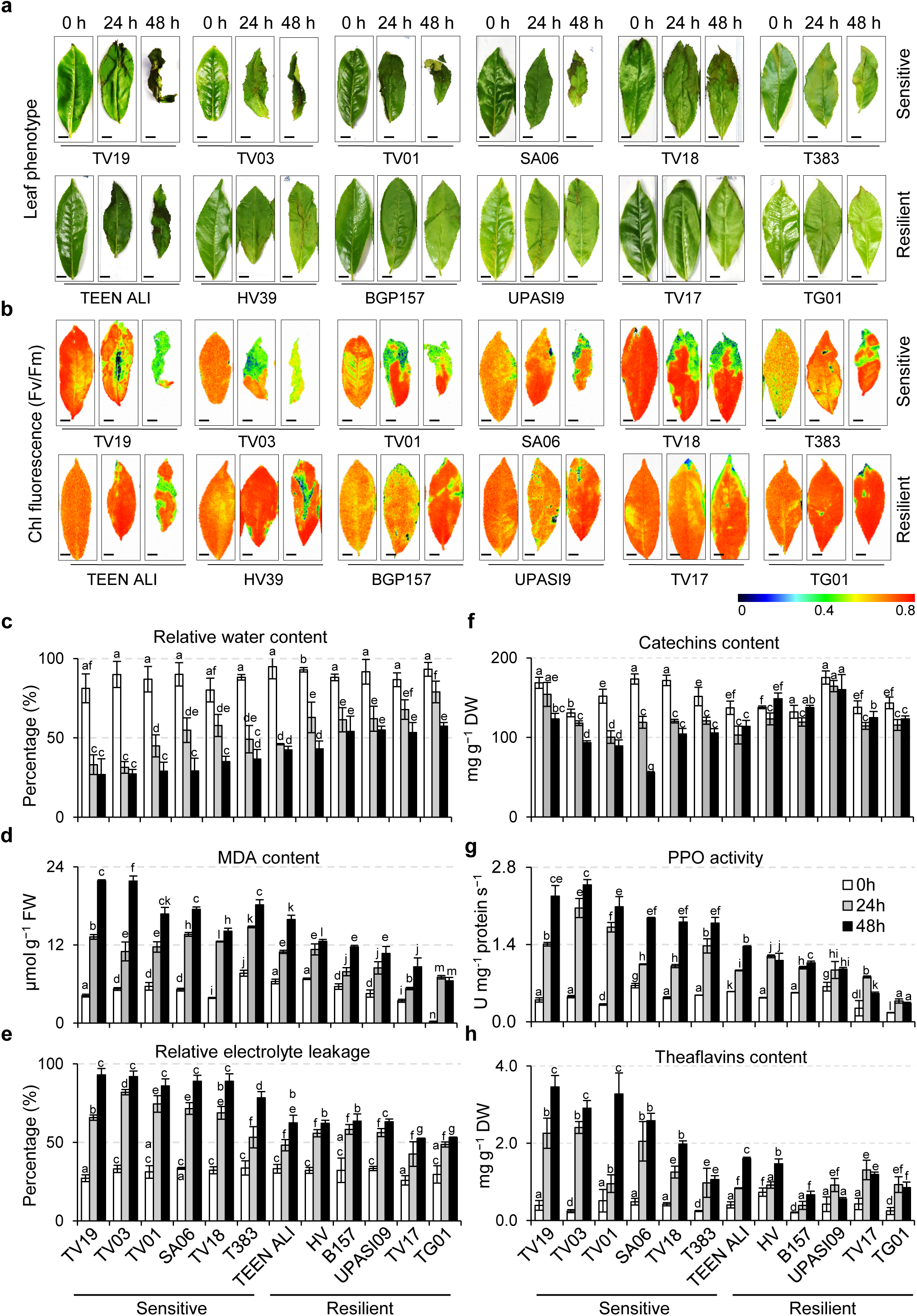
Drought induces oxidative stress, PPO activity, and cell death in Tea. **a** Macroscopic phenotype. **b** Chlorophyll fluorescence representing PSII efficiency. **c** Relative water content (RWC). **d** MDA accumulation as a marker of lipid peroxidation. **e** Relative electrolyte leakage (REL) representing cell death. **f** Polyphenol oxidase (PPO) activity. **g** Catechins content. **h** Theaflavins content. Shoot cuttings from 12 Tea genotypes were incubated in Hoagland’s nutrient medium containing 10% PEG for a period of 48□h. Third leaf was utilized for all experiments (**a-h**). For chlorophyll fluorescence (b),dark-adaptedd leaves were placed under actinic light and visualized in FluorCam 800. Images in (**a** and **b**) are representative of three biological replicates. Metabolites (catechins and theaflavins) were quantified using UPLC-MS. Data in (**c-h**) represents the mean of three independent biological replicates. Error bars indicate standard deviation (SD). Lower case letters indicate statistically significant differences between the mean values (*P*□<□0.05, one-way analysis of variance with post hoc Tukey’s Honest Significant Difference (HSD) test). Scale bar in (a and b) = 2 cm.

Given that catechins are implicated in stress response in Tea ^30^, we analyzed the impact of drought on their levels. Interestingly, as anticipated, drought positively impacted catechin accumulation (Fig. 1f). It was found that though transcriptional expression of catechins biosynthesis pathway genes was upregulated (as depicted in TV03 and TV17 genotypes; Supplementary Fig. 1b), the catechins content was slightly declining with stress in all genotypes (Fig. 1f). However, it was noted that resilient genotypes maintained higher catechins contents, whereas the susceptible genotypes experienced a rapid decline with stress (Fig. 1f, Supplementary Fig. 2a-b, Supplementary Data 1). This observation agrees with earlier reports where higher catechins content is directly related to better stress tolerance ^30^. Reduction in catechins content despite transcriptional upregulation indicated their possible utilization or conversion. Since drought induces oxidative stress in chloroplasts, this likely activates PPO, which converts catechins into theaflavins. As suggested in the literature ^46^, indeed drought-induced oxidative stress in chloroplasts was found to augment the catalytic activity of PPO in all genotypes (Fig. 1g). In line with the higher oxidative stress (Fig. 1b), the PPO activity was also significantly higher in case of susceptible genotypes (Fig. 1g). As PPO activity relates to the oxidation of catechins into TFs, we then estimated their levels. As anticipated, increased PPO activity resulted in an increased accumulation of TFs with stress in given genotypes, where susceptible ones showed a much higher accumulation compared to resilient ones (Fig. 1h, Supplementary Fig. 3a-b, Supplementary Data 1). We also affirmed that the accumulation of catechins and TFs is transcriptionally regulated, where drought positively impacted the expression of genes encoding the flavonoid pathway enzymes, including PPO (Supplementary Fig. 1b).

Collectively, it was found that drought induces oxidative stress in the studied genotypes in varying degrees. In response to drought, plants accumulate protective catechins in a transcriptionally regulated manner. In addition, drought also induces the activity and expression of PPO, resulting in the marked accumulation of TFs.

### PPO localizes in chloroplast and undergoes stress-induced proteolytic activation

Earlier studies on PPO indicated its localization in thylakoids of chloroplasts ^43, 52, 53^. It has also been shown to exist in two forms: latent and activated (formed after C-terminal cleavage) ^52, 54^. However, it is not clear how PPO protein cleavage occurs *in planta*. To strengthen our proposition and to reveal how PPO would be activated and function on its substrates, we investigated its localization and cleavage, if it occurs, in chloroplasts. We first analyzed the sequences of three PPO proteins annotated in *Camellia sinensis* var. Assamica genome with already characterized PPOs from *Prunus armeniaca* and *Vitis vinifera*. Protein sequence alignment revealed more than 50% sequence identity of *Cs*PPOs with *Pa*PPO and *Vv*PPO (Supplementary Fig. 4a). Alignment data and cognate *in silico* analyses affirmed the presence of an N-terminal plastid transit peptide (pTP), followed by a transit peptide for thylakoid (tTP) in all three PPO proteins (Fig. 2a and Supplementary Fig. 4b). Alignment also revealed formation of around 56 kDa mature protein and a probable proteolytic site, generating a cleaved products of around 37 and 19 kDa (Fig. 2a and Supplementary Fig. 4a-b).

**Fig. 2:**
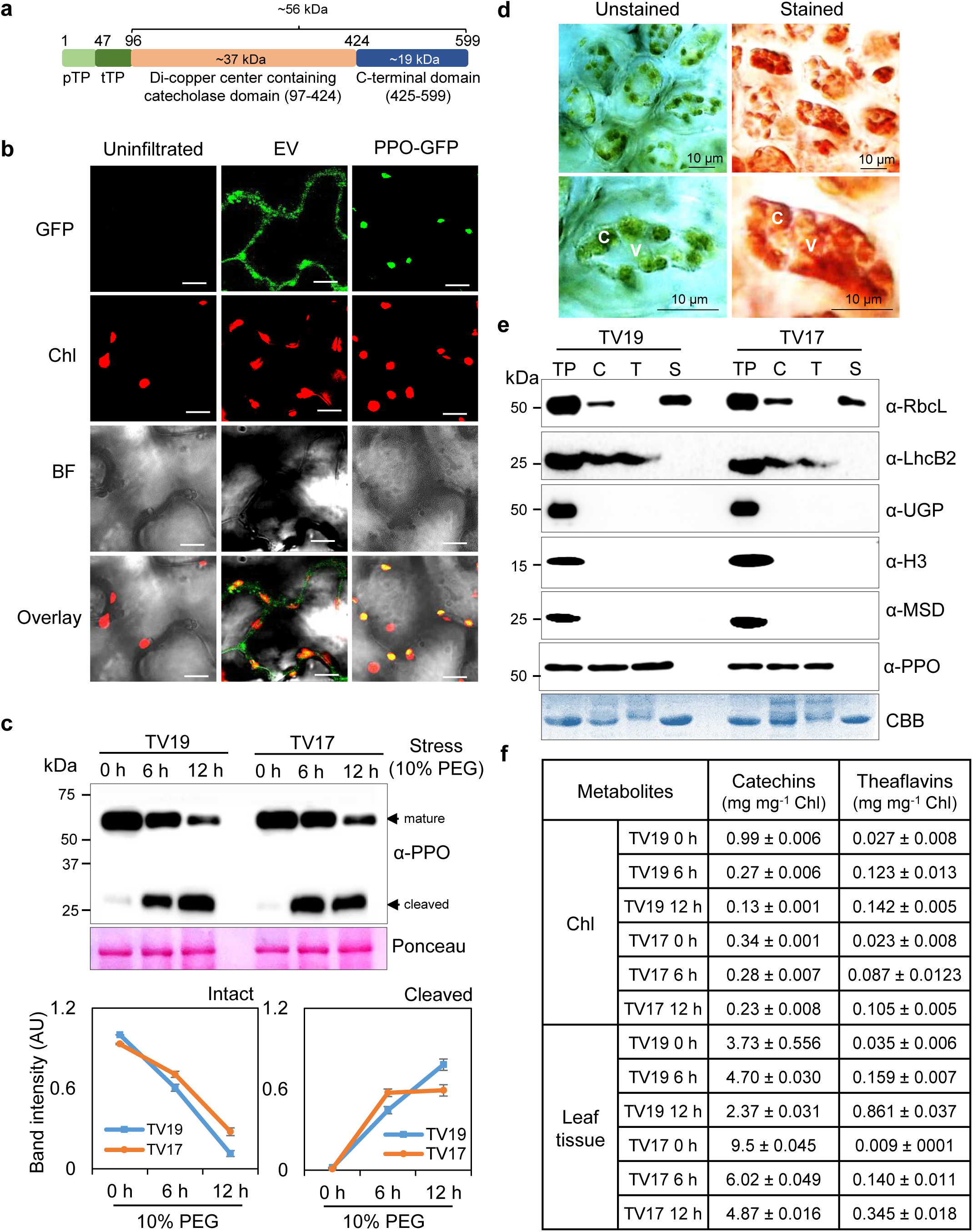
Chloroplast-localized PPO is activated under drought and likely oxidizes catechins in the chloroplast. **a** Domain architecture of *Cs*PPO2 protein. Based on the multiple alignment (Supplementary Fig. 4a), *Cs*PPO2 contains an N-terminal plastid transit peptide (pTP), followed by a transit peptide for thylakoid (tTP), catecholase domain and a C-terminal cleavable domain. **b** Sub-cellular localization of *Cs*PPO-GFP in chloroplasts. The gene construct, *35S::PPO-GFP* was expressed transiently in the leaves of four-week-old *Nicotiana benthamiana* (Benth) plants. After 48 h of infiltration, leaf discs were analyzed under confocal microscope. The green fluorescence shows GFP, and the red fluorescence represents chlorophyll autofluorescence. Scale bar = 10 µm. **c** Immunoblot analysis of steady state levels of *Cs*PPO in Tea. Total proteins extracted from the third leaves of TV19 and TV17 plants subjected to 10% PEG stress were immunoblotted using anti-PPO antibody. Blots were stained with Ponceau S to show equal loading. The band intensities were quantified using ImageJ and represented as arbitrary units. Data represent the mean ± SD of three independent biological replicates. **d** Flavonoid staining in Tea leaves. Hand sections of Tea leaves were stained with Vanillin-HCl. Left panels show unstained sections with green, whereas right panels show red-stained chloroplasts, respectively. C: chloroplast, V: vacuole. Scale bar = 10 µm. **e** Isolation and fractionation of intact chloroplasts. Chloroplasts (C) extracted from TV19 and TV17 were fractionated and analyzed together with total proteins (TP). The fractions were immunoblotted using anti-RbcL,anti-LhcB2, anti-UGPase (UGP), anti-Histone 3 (H3), anti-MnSOD (MSD) as thylakoid membrane (T), stroma (S), mitochondrial, nuclear, and cytosolic marker proteins, and CBB-staining to ensure intactness and purity of chloroplasts. **f** Quantification of Catechins and Theaflavins in total leaf and isolated chloroplasts. The calculations were made in reference to chlorophyll content. Data represent the mean ± SD of three independent biological replicates.

To affirm the localization, we then transiently expressed GFP-tagged *CsPPO2* under the control of 35S promoter in *N. benthamiana* (Fig. 2b). In agreement with *in silico* analysis, the PPO-GFP signals were observed in chloroplasts (Fig. 2b). The GFP signal was intense and appeared to be a membrane signal. Next to affirm the impact of stress on PPO, we isolated total proteins from TV19 and TV17 genotypes subjected to drought and immunoblotted using anti-PPO antibody targeting C-terminal end (Fig. 2c). Immunoblot revealed the presence of ∼60 kDa mature PPO band and a smaller band of ∼27 kDa, indicating the proteolysis of mature PPO protein (Fig. 2c). Interestingly, the mature PPO band showed a gradual stress-induced decline with an apparent increase in the smaller cleaved band in both genotypes (Fig. 2c). Interestingly, the proteolysis of mature PPO was much faster in the susceptible (TV19) genotype compared to the resilient one (TV17), correlating with higher PPO activity in the susceptible genotype. These results show advocates PPO as a stress sensor that undergoes a stress-induced proteolytic activation.

Although catechins, the substrates of PPO in tea, are stored mostly in vacuoles, they also accumulate in cytosol, vessel walls, and chloroplasts. To explore the possibility that activated PPO acts upon catechins in the chloroplast, we next investigated whether catechins are accumulated in chloroplasts, as indicated earlier ^45^. The hand sections of the third leaf of the Tea shoots of TV19 genotype were stained with Vanillin-HCl. As shown earlier by Liu, Gao ^45^, we also observed the red-stained chloroplasts and vacuoles (Fig. 2d), indicating catechin accumulation. To further affirm these results, isolated intact chloroplasts and leaf tissue were analyzed for metabolites using UPLC-MS. Before proceeding with metabolic analysis, the intactness and purity of chloroplasts were analysed through fractionation. Chloroplasts isolated from both genotypes under control conditions (0 h stress) were fractionated and immunoblotted using antibodies for PPO, and thylakoid, stromal, nuclear, mitochondrial, and cytosolic marker proteins (Fig. 2e). Presence of RbcL and LhcB2 in stromal and thylakoid fractions affirmed the intactness of chloroplasts, whereas absence of any band in chloroplast fractions against UGPase, H3, and MnSOD (MSD) confirmed their purity. As anticipated, the presence of a ∼60 kDa band in the total protein, chloroplast, and thylakoid fractions also affirmed the presence of PPO in thylakoids (Fig. 2e).

As shown earlier by Liu, Gao ^45^ and depicted by our Vanillin-HCl staining (Fig. 2d), catechins were detected in intact chloroplast fractions of susceptible (TV19) and resilient (TV17) genotypes. Quantification in reference to chlorophyll content revealed that the susceptible genotype accumulated almost ¼-folds total catechins in chloroplasts compared to leaf tissue, whereas the resilient genotype accumulated almost 27-folds less catechins in chloroplasts than in leaf tissue, under normal conditions. Under normal conditons, almost 3-fold higher catechins were found in chloroplasts of the susceptible genotype whereas in case of leaf tissue, resilient genotype accumulate 3-fold higher catechins (Fig. 2f). In contrast to catechins, negligible amounts of total TFs were detected in the chloroplasts of both genotypes; however, upon stress, the TFs started accumulating with a concomitant gradual decline in catechin levels in the chloroplasts of both genotypes (Fig. 2f). These results showed that though major fractions of catechins are stored in cytosol, a significant proportion translocates into chloroplasts and oxidized into TFs by stress-activated PPO. Though a thorough investigation is needed to define the precise mechanism, given their reactive nature, TFs are likely exported from the chloroplast, perhaps through drought-induced chloroplast leakage or membrane contact sites, and interact with cytosolic machinery.

### Drought- and TFs-induce a similar reprogramming of nuclear gene expression, activating an ER stress-like response

Considering the drought-induced upregulation of PPO activity and accumulation of TFs in chloroplasts, we hypothesized that TFs might play a signaling role and activate cognate stress response. To test this conception, we revisited the publicly available datasets from an earlier study which reported transcriptional changes in two contrasting genotypes, TV03 (susceptible) and TV17 (resilient) subjected to 48 h of PEG-induced drought using ^29^.. Using the differentially expressed genes among the two genotypes (1641 upregulated and 2081 downregulated in TV03 compared to TV17) (Fig. 3a, Supplementary Data 2-3), we performed the gene ontology (GO) enrichment analysis (Fig. 3b, Supplementary Data 4-5). Interestingly, GO analysis revealed that genes involved in canonical drought response, including response to water deprivation, stomatal closure, response to ABA, and response to oxidative stress, exhibited higher expression in TV03. In addition, the genes involved in phenylpropanoid and flavonoid biosynthesis also showed heightened expression in TV03 compared to TV17 (Fig. 3b-d). In addition, a large subset of genes involved in protein quality control and cell death were also induced. Genes involved in folding stress, proteasome assembly, ERAD pathway, autophagy, vesicle-mediated transport, hypersensitive response, response to defense hormones JA and SA, and PCD also showed a higher expression in TV03 (Fig. 3b-c). We then validated the expression pattern of enriched genes using qRT-PCR in plant samples of both TV03 and TV17 subjected to 24 h and 48 h stress. As observed in RNAseq data, several marker genes of folding stress (ER-UPR and HSR), autophagy, and cell death showed upregulation with stress in both genotypes. Interestingly, the expression of ER-stress, UPR and cell death-related genes was higher in drought-sensitive genotype TV03, whereas the drought-resilient genotype TV17 showed higher expression of HSR-related and negative regulators of autophagy (Fig. 3d).

**Fig. 3:**
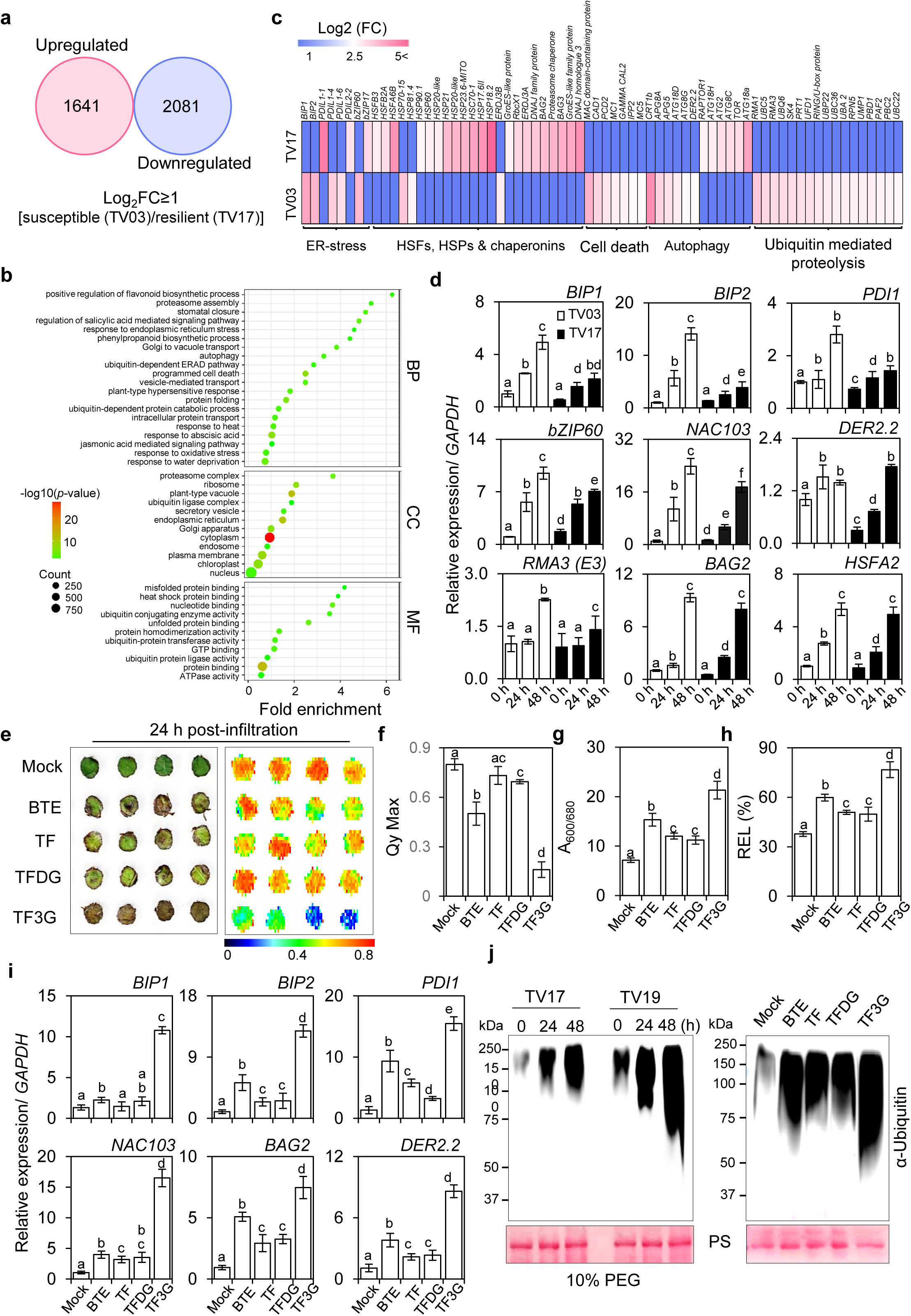
Drought-induced reprogramming of nuclear gene expression activates an ER stress-like response, which is mimicked by exogenous treatment of oxidized flavonoids. **a** Venn diagram showing the differentially expressed genes under drought in susceptible (TV03) and resilient (TV17) Tea genotypes (data adapted from Parmar et al., 2019). **b** Gene Ontology (GO) enrichment analysis of genes upregulated in the susceptible genotype (TV03) presented as an enrichment bubble plot. GO analysis was performed using DAVID, and data were visualized with SR plot. **c** Heatmap showing drought-induced expression of genes involved in general ER stress-like response. **d** Relative expression of ER stress-responsive genes in Tea. qRT-PCR was performed using equal amounts of cDNA prepared from RNA extracted from third leaves of susceptible and resilient genotypes subjected to drought. *CsGAPDH* was used as an internal control. **e** Macroscopic phenotype and chlorophyll fluorescence of Tea leaf discs infiltrated with black tea extract (BTE), theaflavins (TFs), TF3G, TFDG (200 µM each), or DMSO (mock control). After infiltration, discs were incubated in half-strength Hoagland nutrient medium for 24 h, photographed, and analyzed by FluorCam 800 for PSII efficiency following dark adaptation. **f** Quantification of PSII efficiency (Fv/Fm) of treated leaf discs. **g** Cell death quantification using Evans blue staining of treated discs (absorbance at 600/680 nm). **h** Relative electrolyte leakage (REL) in treated discs. Conductivity was measured before and after autoclaving. **i** Relative expression of ER stress- and cell death-associated genes in treated discs, normalized to *CsGAPDH*. **j** Drought and TF-induced polyubiquitination of proteins. Equal amounts of protein extracts from the third leaf of drought-treated susceptible and resilient genotypes were immunoblotted with anti-Ubiquitin antibody. Data in **(d, f-i)** represent mean ± SD of three independent biological replicates. Lowercase letters indicate statistically significant differences (*P*□<□0.05, one-way ANOVA with Tukey’s HSD test).

Next, to assess if the expression of such a response in the susceptible genotype is due to increased PPO activity and TFs accumulation, we exogenously fed the leaf discs of TV19 genotype with TFs-rich black tea extract [BTE; containing more than 80% TFs (TF and TF gallates)], and three commercially available pure theaflavins, namely theaflavin (TF), theflavin-3-gallate (TF3G) and theflavin-di-gallate (TFDG). The BTE contains a mixture of four different TFs, we affirmed the percentage proportion of each theaflavin using UPLC-MS (Supplementary Fig. 5). As commercially only three TFs, namely, TF, TF3G, and TFDG, are available, they were only quantified using pure standards (Supplementary Fig. 5). Results indicated that BTE contains 3.49% TF, 6.35% TF3G and 14.77% TFDG (Supplementary Fig. 5). As anticipated, treatment of BTE and pure TFs resulted in browning of discs with a concomitant loss of chlorophyll fluorescence and PSII activity, and an increase in cell death (Fig. 3e-h). These phenotypes were further manifested with a significant induction of key ER stress marker genes, including *Binding Protein 1 and 2 (BIP1* and *BIP*2) encoding ER-localized HSP70 protein and *Protein disulfide-isomerase 1* (*PDI1*) encoding a key enzyme involved in protein folding in ER, and cell death/autophagy/ERAD genes, *NAC103*, *Bcl-2 associated athanogene 2* (*BAG2*) and *Derlin-2.2* (*DER2.2*) (Fig. 3i). Though all TFs showed such an impact, TF3G induced more prominent browning, ER-stress, and cell death (Fig. 3i), indicating its potent nature.

Since polyubiquitination marks the proteins for degradation, its presence represents a folding stress. To ascertain this, we next examined the stress- and TFs-induced polyubiquitination in total leaf proteins in susceptible and resilient genotypes (Fig. 3j). As revealed by gene expression, immunoblot analysis with anti-Ubiquitin antibody revealed a heightened polyubiquitination of protein in the susceptible genotype (TV19) compared to the resilient one (TV17) (Fig. 3j). Similarly, treatment of BTE or pure TFs also induced protein polyubiquitination, where BTE and TF3G showed a more potent impact as observed in cell death and gene expression patterns (Fig. 3j).

Collectively, re-analysis of publicly available transcriptome dataset, subsequent qRT-PCR-based target gene expression under drought, and a similar response by BTE and pure TFs feeding (Fig. 3), revealed resemblance with general stress response (GSR) ^20^. Drought-induced activation of PPO, and the resemblance of drought- and TFs-induced gene expression and cognate response with GSR, it was plausible to propose that drought(stress)-induced transcriptional changes and cognate response in Tea plants are mediated by PPO-generated TFs.

### Silencing and overexpression of PPO modulate drought-induced phenotypes in susceptible and resilient genotypes of Tea

To further ascertain the plausible role of PPO (and TFs), we next utilized a reverse genetics-based approach by modulating its expression. Developing stable transformants in tea is quite challenging, so we utilized the Tobacco Rattle Virus (TRV)-based system to silence *PPO* expression in tea plants. Fragments designed for VIGS of *PPO* and *Phytoene desaturase (PDS)* gene of Tea were amplified and cloned into the pTRV2 vector. The Agrobacterium containing pTRV1/pTRV2 (empty vector control), pTRV1/pTRV2-*PDS* (*pTRV:PDS*; as a positive control) and pTRV1/pTRV2-*PPO* (*pTRV:PPO*) were inoculated in shoot cuttings of TV03 (drought susceptible genotype) following a recent method developed by Peng, Xue ^55^. The inoculated cuttings showed the appearance of new buds after 3 weeks, which grew into leaves and turned pale to yellowish in varying degrees after 5-6 weeks in the case of *pTRV:PDS* lines compared to lines containing empty vectors (EV), which showed green new leaves (Supplementary Fig. 6a). Based on the yellowish phenotype, the *pTRV:PDS* lines were named L1-L4 lines (Supplementary Fig. 6a). The yellowish leaf phenotype in different *pTRV:PDS* VIGS lines was affirmed by the estimation of their chlorophyll contents (Supplementary Fig. 6b). As anticipated, the VIGS lines showed a reduction in the expression of *PDS* in proportion to the decrease in green color, indicating the efficacy of VIGS in drought-susceptible genotype, TV03 (Supplementary Fig. 6c). In case of *pTRV:PPO*, the lines were named from L1-5 based on the expression of *PPO* compared to EV (Fig. 4a). All five lines showed varying degrees of repression in the *PPO*; L1, L2 and L3 showed a slight decline, whereas L4 and L5 showed a drastic reduction in the expression compared to EV (Fig. 4a). As depicted with the degree of repression of *PPO*, the *pTRV::PPO* lines showed proportional resilience levels to PEG-induced drought with lesser leaf wilting/browning and reduced relative electrolyte leakage (Fig. 4b-c). The L4 and L5 were almost unaffected and showed a similar phenotype to the unstressed EV control (Fig. 4b-c). These results were further affirmed by a reduction in the activity of PPO in different lines, which was in proportion with high catechins and markedly lower TFs accumulations (Fig. 4d). These findings reinforced the notion that PPO suppression effectively limited TFs production. We next investigated the expression of key ER stress-related genes to understand the implications of PPO silencing on stress signaling. In agreement with better phenotype, the decreased PPO expression coincides with repression of key ER stress markers, namely, *BIP1*, *BIP2*, *PDI1*, and *bZIP60*, and cell death regulators/ER-associated degradation components, *NAC103* and *DER2*, revealed a significant downregulation in *PPO*-silenced plants (Fig. 4e). The degree of suppression of these genes was consistent with the extent of *PPO* silencing, further supporting the link between PPO activation, TFs accumulation, and stress-induced transcriptional responses.

**Fig. 4:**
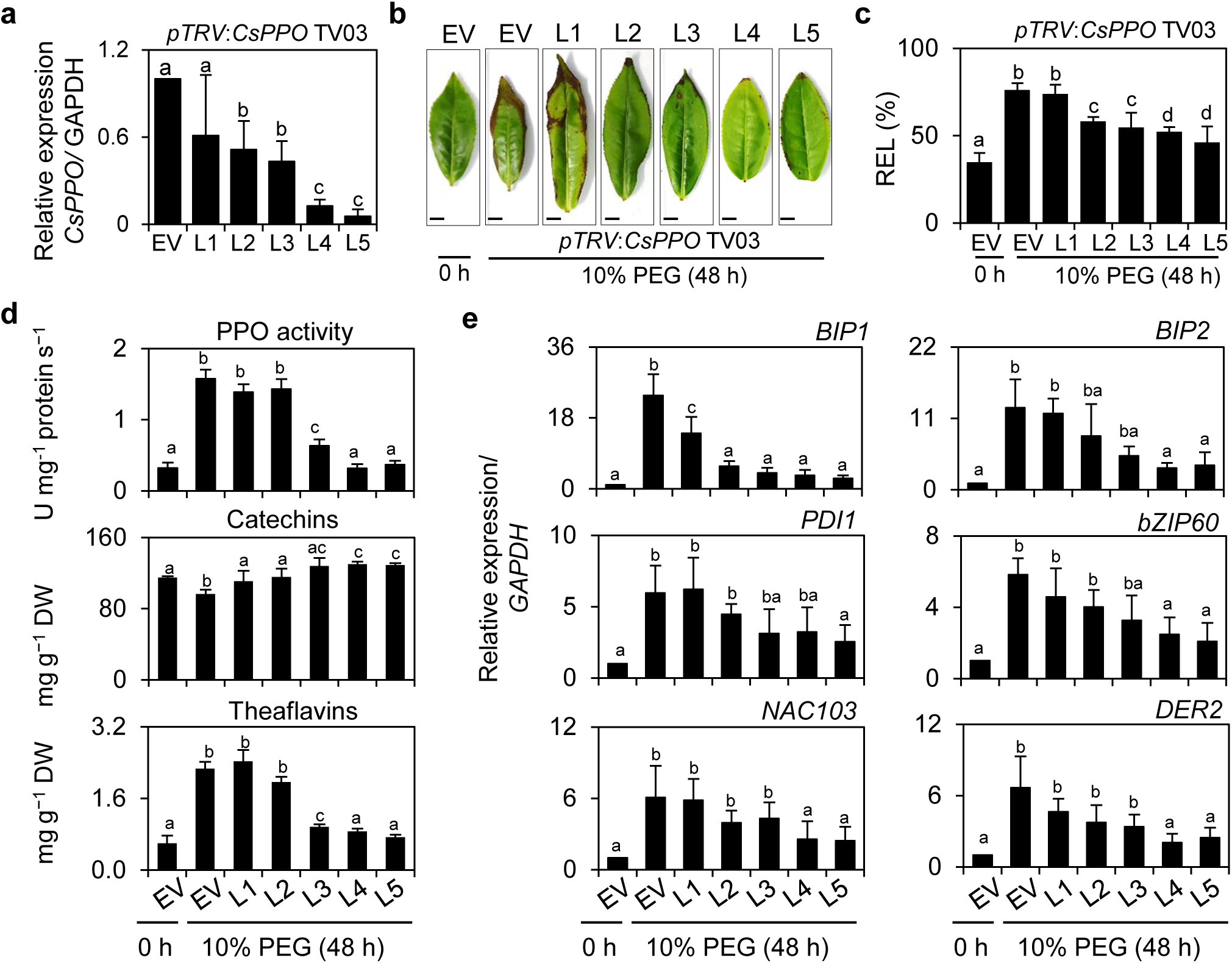
Virus-induced silencing of Cs*PPO* resulted in the suppression of drought-induced cell death. **a** Transcriptional expression of *CsPPO* in empty vector (EV) and *pTRV::CsPPO* VIGS lines measured by qRT-PCR. *CsGAPDH* was used as an internal control. **b-d** Macroscopic leaf phenotype **(b)**, relative water content (RWC) **(c)**, and relative electrolyte leakage (REL) **(d)** of EV and *pTRV::CsPPO* lines subjected to PEG-induced drought. **e** PPO activity and metabolite quantification. PPO activity, catechin levels, and theaflavin levels were measured in EV and VIGS lines. Metabolites were quantified using UPLC-MS. **f** Relative expression of genes involved in ER stress response and programmed cell death in EV and VIGS lines under drought stress, normalized to *CsGAPDH*. For all assays **(b–f)**, VIGS lines were incubated in Hoagland’s nutrient medium containing 10% PEG for 48□h, and the third leaf was used for experiments. Data represent mean ± SD of three independent biological replicates. Lowercase letters indicate statistically significant differences between mean values (*P*□<□0.05, one-way ANOVA with Tukey’s HSD test). Scale bars in **(a)**, 2□cm.

Next we transiently expressed the *35S:CsPPO2-GFP* construct (Fig. 2b), in drought-resilient genotype TV17 (Fig. 5a-e). The PPO overexpression (*OePPO*) lines showed no growth defects under normal conditions; however, they showed hypersensitivity with a rapid browning and higher REL compared to native TV17 plants (Fig. 5a-b). Transcript-level expression revealed ∼1.6-fold upregulation of *CsPPO* in *OePPO* lines compared to native TV17 plants, which was further affirmed by immunoblot analysis (Fig. 5c-d). In agreement, the PPO activity was also found to be elevated by around ∼1.6 fold, which showed a gradual increase with stress (Fig. 5d). Surprisingly, the native PPO protein was not detected in *OePPO* lines; instead, they showed the expression of *Cs*PPO2-GFP with both anti-PPO and anti-GFP antibodies (Fig. 5e). Nevertheless, both native TV17 and *OePPO* lines exhibited a stress-induced proteolysis of PPO, which was slightly higher in case of *Oe* lines (Fig. 5e). The *Oe* lines also exhibited a heightened stress-induced expression of key ER stress markers, ER-associated degradation components and cell death regulators, *BIP1, NAC103*, *DER2*.3 and *BAG2*, compared to native TV17 plants (Fig. 5f).

**Fig. 5:**
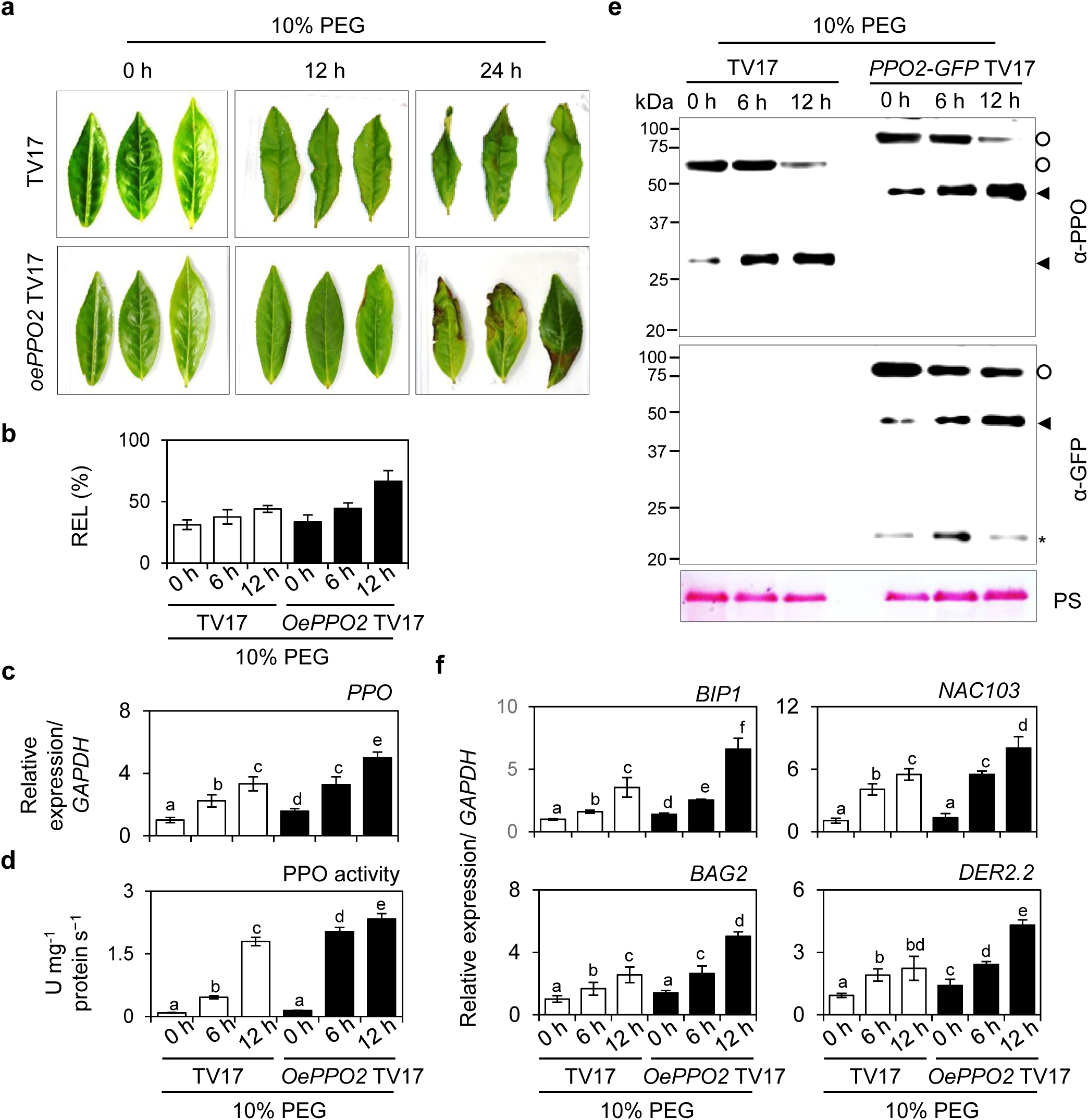
Overexpression of Cs*PPO* resulted in the augmentation of drought- and TFs-induced signaling. **a** Macroscopic leaf phenotype of wild-type TV17 and 35S::CsPPO2-GFP overexpression lines under PEG-induced drought. **b** Relative electrolyte leakage (REL) in TV17 and overexpression lines. **c** Relative transcript levels of *CsPPO2* measured by qRT-PCR. *CsGAPDH* was used as an internal control. **d** PPO enzymatic activity and metabolite levels of catechins and theaflavins. Metabolites were quantified using UPLC-MS. **e** Immunoblot analysis of PPO protein levels and stress-induced proteolysis of *Cs*PPO2 in TV17 and overexpression lines. Equal loading was confirmed by Ponceau S staining. **f** Relative expression of ER stress- and cell death-associated genes in TV17 and overexpression lines under PEG-induced drought, normalized to *CsGAPDH*. For all assays **(a-f)**, plants were incubated in Hoagland’s nutrient medium supplemented with 10% PEG for 12□h, and the third leaf was used for experiments. Data in **(b-d, f)** represent mean ± SD of three independent biological replicates. Lowercase letters indicate statistically significant differences among mean values (*P*□<□0.05, one-way ANOVA with Tukey’s HSD test).

These findings strengthened the proposition that drought-induced activation of PPO leads to the formation of chloroplastidic TFs, which in turn orchestrate inter-organelle signaling events associated with ER stress and cell death. By suppressing or overexpressing *PPO*, we observed a disruption or augmentation in this signaling cascade.

### Drought-induced proteolytic PPO activation and consequent downstream signaling is not limited to Tea

As the PPO is an evolutionarily conserved enzyme present in different plant species, we next examined whether drought-induced activation of PPO also occurs in other PPO-containing plant species. For this, we subjected PPO-containing plants, tomato and wheat, to PEG-induced drought stress (Supplementary Fig.7-8). Both species exhibited leaf wilting with stress duration due to a decrease in RWC and chlorophyll fluorescence (indicating a reduction in PSII efficiency) (Supplementary Fig.7a-c and 8a-c). Additionally, there was an increase in REL, causing cellular damage and death (Supplementary Fig.7d and 8d). In line with drought-induced activation of PPO in tea (Fig. 2c), PPO activity in both tomato and wheat also increased with the duration of drought stress (Supplementary Fig. 7e and 8e), suggesting that drought positively impacts PPO activity. Western blot analysis further revealed PPO proteolysis under stress in both tomato and wheat, similar to that observed in tea plants (Supplementary Fig. 7f and 8f), supporting the idea that PPO functions as a stress sensor. Drought-induced upregulation of key ER-stress markers *PDI, CNX, bZIP60*, and *IRE1*, as well as cell death markers *BI-1*, *MC5*, and *MC8* in tomato, reinforced the notion that stress-induced activation of PPO is linked with the activation of ER stress-like response leading to UPR and cell death (Supplementary Fig. 7g). These results revealed that drought(stress)-induced activation of PPO and its key role in orchestrating stress signaling seems to be an evolutionarily conserved phenomenon in plants.

### Exogenous treatment of theaflavin-rich black Tea extract or pure theaflavins induces ER stress and cell death in Arabidopsis

To further affirm the signaling role of TFs, we utilized the model plant *Arabidopsis thaliana*. The exogenous foliar treatment of TFs-rich BTE to 5-day-old seedlings showed gradual growth inhibition, bleaching, and reduction in PSII efficiency in a dose-dependent manner (Fig. 6a-b). Interestingly, the BTE treatment also induced dose-dependent cell death, as shown by trypan blue staining in the leaves (Fig. 6c). To affirm that such a response was largely because of TFs, the seedlings were also treated with Green Tea Extract (GTE; rich in catechins) (Fig. 6a-c and Supplementary Fig. 5a). The results indicated that GTE has no impact on growth and PSII quantum efficiency and showed no cell death (Fig. 6a-c), suggesting that the impact was due to TFs. To further unveil the impact of TFs, we treated seedlings with pure TFs in a dose-dependent manner (50, 100, and 200 µM) and analyzed after 24 h (Fig. 6d-e). Interestingly, it was revealed that all tested TFs showed slight growth inhibition (not quantified), bleaching, and cell death at higher concentrations (200 µM); however, TF3G showed a significant response and showed cognate phenotypes at lower concentrations (50 µM) also (Fig. 6d-e). We then tested the time-dependent impact of TFs at a fixed concentration (200 µM). These results revealed that TFs-induced cell death is time-dependent and appears after 12 h of treatment in the case of TF3G and 18 h in case of TF and TFDG (Supplementary Fig. 9a-b). The dose- and time-dependent appearance of cell death indicated that the levels of TFs, which are directly proportional to stress intensity (Fig. 1g), determine the stress response in Tea plants as well. Given the apparent potency of TF3G, it was selected for downstream studies.

**Fig. 6:**
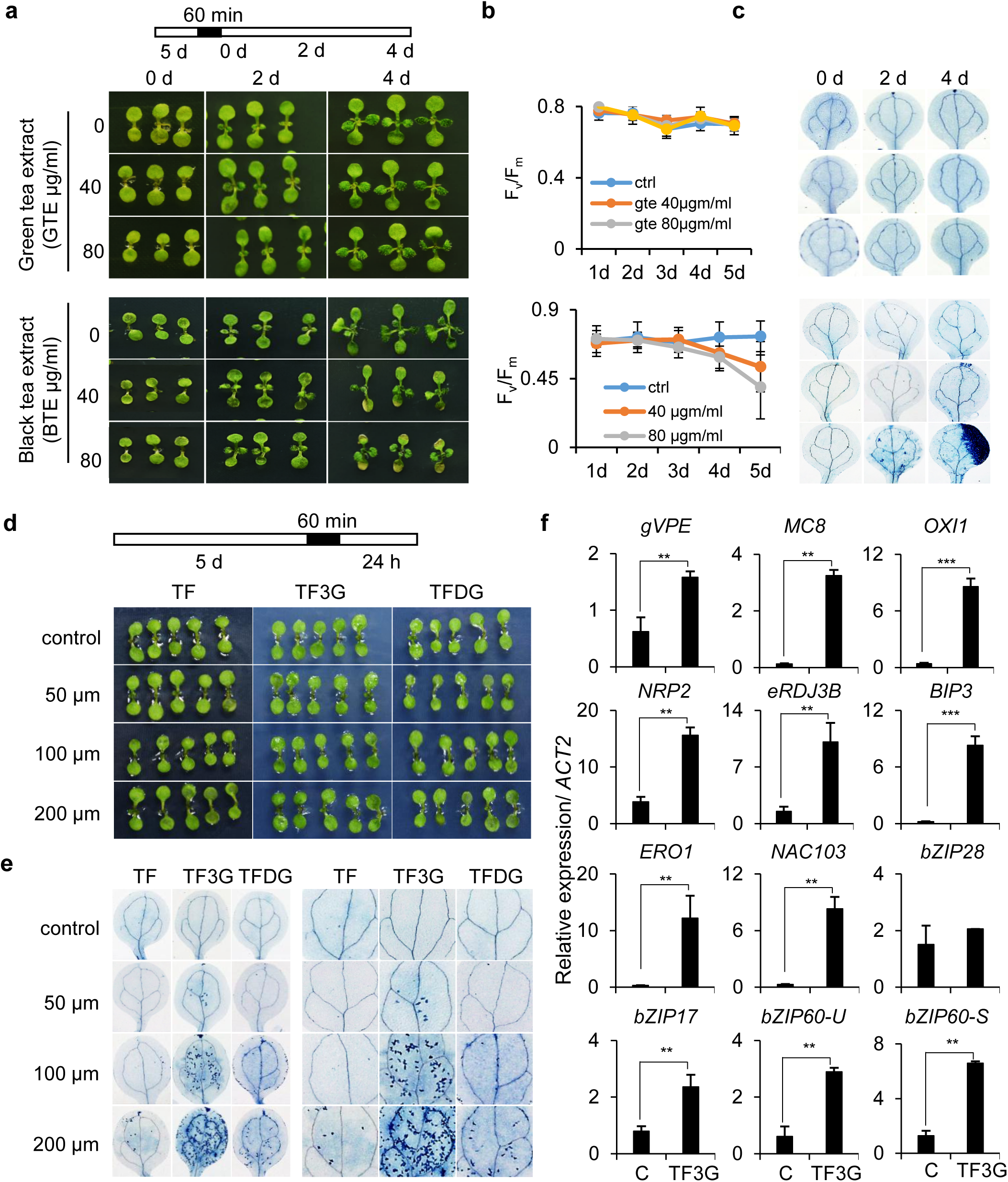
Oxidized flavonoids (theaflavins) repress growth and induce programmed cell death likely via ER stress. **a-c** Effect of black tea extract (BTE) and green tea extract (GTE) on Arabidopsis seedlings. **a** Macroscopic phenotype. **b** PSII efficiency (Fv/Fm) measured using FluorCam 800. **c** Cell death visualized by trypan blue (TB) staining. Five-day-old wild-type (Col-0) seedlings grown on MS agar plates were submerged in BTE or GTE solutions for 60□min, after which the solution was removed and seedlings were grown in a chamber. Five seedlings from each treatment were harvested every 2 days. Images are representative of three independent biological replicates. Data in **(b)** represent mean ± SD of three biological replicates. **d-e** Impact of pure theaflavins on phenotype and cell death. Five-day-old Col-0 seedlings were vacuum infiltrated with TF, TF3G, or TFDG at the indicated concentrations for 5□min and incubated for 55□min in the same solution. Seedlings were then transferred to fresh liquid MS medium and grown for 24□h before imaging **(d)** and TB staining **(e)**. **f** Relative transcript levels of ER stress- and cell death-associated genes in seedlings treated with 200□µM TF3G compared with mock controls. *ACT2* was used as an internal control. Values represent mean ± SD of three independent biological replicates. Asterisks denote statistically significant differences between treatments (\**P*□<□0.05; \*\**P*□J<□0.01, Student’s t-test).

To understand how TFs induce bleaching and cell death phenotypes, we analysed the expression of key cell death and ER stress genes in seedlings treated with TF3G (Fig. 6f) ^56^. Transcript expression data revealed a significant upregulation of cell death marker genes, including cysteine-type endopeptidase, namely, *Gamma*-*Vacuolar Processing Enzyme* (gVPE), which upregulates under stress conditions and mediates toxin-induced cell death ^57^ and *Metacaspase 8* (*MC8*), which is known to promote PCD in response to light-induced oxidative stress ^58^, and *Oxidative inducible 1* (*OXI1*), a well-known cell death mediator in response to oxidative stress initiating in chloroplast ^59^. We also observed a heightened expression of a negative regulator of PCD, *ASPARAGINE-rich protein 2* (*NRP2*), which, along with *NRP1*, restricts or limits cell death. Expression of genes involved in PCD indicates that TF3G-induced cell death is likely genetically controlled. Elevated expression of negative regulator *NRP2* accords with this notion and also underlines the reasons for discrete and localized cell death induced by TF3G (Fig. 6e). Given that ER stress can also culminate in cell death; Mishiba et al., 2013) and that transcriptome analysis in Tea indicated the activation of ER stress-like responses (Fig.s 3a-d), we examined the impact of TF3G on UPR genes. Interestingly, key ER-localized proteins involved in folding, including *ERDJ3B*, an ER-lumen localized J domain protein; *Binding Protein 3* (*BIP3*), an ER-localized HSP70 protein and *Endoplasmic reticulum oxireductin 1* (*ERO1*), an oxidoreductase required for oxidative protein folding in the ER, also showed several-fold upregulation upon TF3G treatment (Fig. 6f). Upregulation of ER stress-responsive transcription factors and ER-localized chaperones showed activation and operation of UPR. The UPR is mediated by two canonical pathways involving either of the ER-localized transcription factor modules, namely bZIP28/bZIP17 or IRE1-bZIP60 ^60^. We then determined the involvement of these modules. Accumulation of unfolded proteins activates ER-localized ER stress sensor transmembrane kinase, namely inositol-requiring ER-to-nucleus signaling protein (IRE1α), an endonuclease that splices mRNA of bZIP60. The involvement of this module can be determined by analyzing the expression of spliced (*bZIP60-S*) and unspliced (*bZIP60-U*) forms of *bZIP60* ^56^. Although both spliced and unspliced forms of *bZIP* showed upregulation, a marked upregulation of the spliced form was observed (Fig. 6f). In case of bZIP28 and bZIP17, which are normally posttranslationally cleaved to be translocated into the nucleus upon ER stress, the latter showed slight upregulation, whereas the former was almost unchanged (Fig. 6f). These results indicate that TF3G-induced UPR is likely mediated through IRE1-bZIP-dependent module. Upregulation of *NAC103* (Fig. 6f), a bZIP60-driven UPR marker transcription factor ^61^, further reinforced the possibility.

### Theaflavins interfere with protein folding machinery to activate ER stress response

To determine the mechanism of how TFs induce slight bleaching, localized cell death, and ER stress response, we surveyed the literature. A recent study in animal systems indicated TFs as potential anticancer molecules that inhibit protein folding and quality control machinery, potentially targeting the HSP90 protein ^62^. A plethora of literature shows that HSP90 is an evolutionarily conserved ATP-dependent chaperone that functions in protein folding downstream of HSP70 ^63^. In addition, it also regulates cellular signaling networks, especially retrograde signaling, operating under normal growth and stress conditions ^64^. Arabidopsis genome comprises seven HSP90 proteins: four localized in cytosol (four; HSP90.1 to HSP90.4) and one each in chloroplast (HSP90.5), mitochondria (HSP90.6), and ER (HSP90.7) ^65^. Given the significant similarity of these HSP90 proteins, their biochemical functions are largely similar. Inhibition of HSP90 in both cytosol and ER has been shown to activate folding stress response, where activation of folding stress response in ER (UPR) is more prominent ^66^. To determine if TFs can bind to HSP90, we performed an *in silico* protein-ligand docking analysis. As sequences of all seven HSP90 were broadly similar, we chose HSP90.1, comprising three domains: the N-terminal ATPase domain, the middle domain important for client binding, and the C-terminal dimerization domain ^67^ and modeled its structure. To further elaborate on the TFs-HSP90 binding, we included geldanamycin (GDA), a known inhibitor of HSP90, as a control (Fig. 7a). The GDA binds to HSP90 at the ATP-binding pocket, competing with ATP to bind to the site ^68^. The resulting analysis revealed that all four TFs have a higher binding affinity to HSP90 compared to GDA (Supplementary Fig. 10, Fig. 7a). Interestingly, as exhibited by the intensity of cell death phenotype, among four TFs, TF3G showed the highest affinity for the HSP90 protein (Supplementary Fig. 10), and hence was utilized in further studies. We then analyzed whether GDA also shows phenotypes similar to TFs. Interestingly, Arabidopsis seedlings treated with GDA and TF3G in a dose-dependent manner showed almost identical dose-dependent phenotypes of slight bleaching of cotyledons and localized cell death (Fig. 7b-c). The TB staining results indicated that the intensity of cell death was slightly higher in TF3G (Fig. 7c). The quantification of dead cells asserted that TF3G was comparatively more potent (Fig. 7d). The cell death was correlated with higher binding affinity of TF3G with HSP90, as observed in docking studies (Fig. 7a). Like TF3G, GDA-induced cell death was also found to be time-dependent (Fig. 7e). These results indicated that TF3G and GDA show similar phenotypes, which are likely HSP90-inhibition dependent. It is well established that GDA activates UPR by inhibiting ER-localized HSP90 (also called Glucose-regulated protein 94; GRP94), which stabilizes IRE1α ^66^. To affirm this conception, we analyzed the time-dependent (12 and 24 h after treatment) impact of TF3G and GDA on the expression of key cell death and UPR genes. As anticipated, both TF3G and GDA induced the expression of *gVPE*, *MC8*, *OXI1* and *NRP2* in a time-dependent manner and to almost similar levels (Fig. 7f). Similar expression was observed for chaperones/co-chaperones, namely, *ERDJ3B*, *BIP3*, *ERO1* (Fig. 7f). As observed in Fig. 5f, the expression of bZIP28 was almost unaffected, whereas bZIP17 showed a slight upregulation upon GDA treatment. Interestingly, both spliced and unspliced forms of *bZIP*60 were upregulated (Fig. 7f), indicating activation of the IRE1-bZIP60 module. The upregulation of *NAC103* and *BIP3* (Fig. 7f) further reinforced the likelihood that TF3G and GDA induce UPR in a bZIP60-dependent manner. The operation of UPR was further reinforced by TF3G- and GDA-induced polyubiquitination in total proteins as observed in anti-Ubiquitin antibody immunoblot (Fig. 7g).

**Fig. 7:**
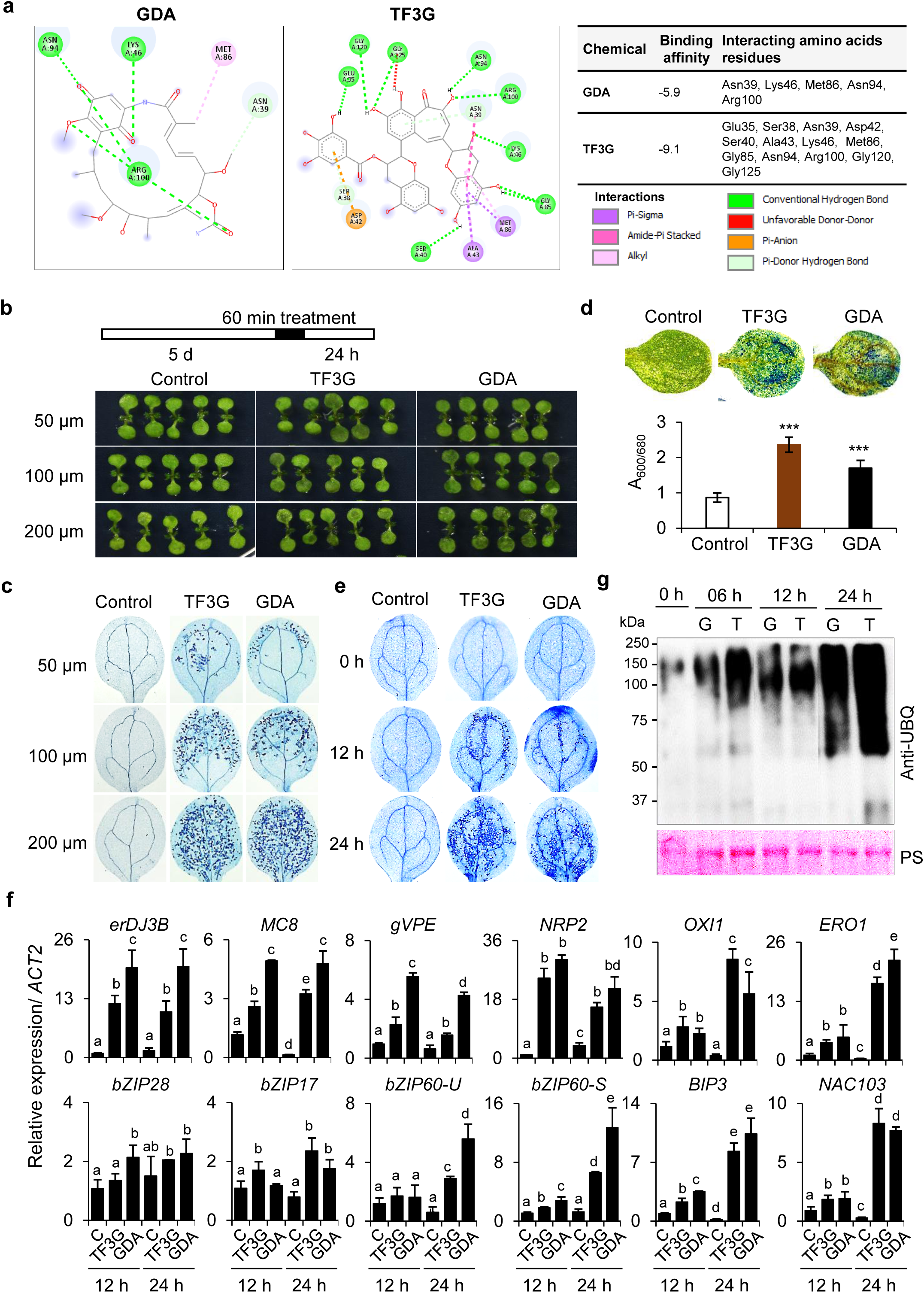
Theaflavins, especially TF3G, inhibit HSP90 (folding machinery) to activate ER stress. **a** *In-silico* docking analysis showing stronger affinity of TF3G to HSP90 compared with the HSP90 inhibitor geldanamycin (GDA). b**-d** Impact of TF3G and GDA on Arabidopsis seedlings. **b** Macroscopic phenotype. **c** Cell death visualized by trypan blue (TB) staining. **d** Cell death quantified by Evans blue (EB) staining, expressed as relative absorbance (A600/A680). Five-day-old Col-0 seedlings were vacuum infiltrated with TF3G or GDA for 5□min, incubated for 55□min in the same solution, and then transferred to liquid MS medium for 24□h before imaging and staining. For EB quantification, five cotyledons per treatment were used. Data represent mean ± SD of three independent biological replicates. **e** Time-dependent cell death progression. Seedlings treated with TF3G or GDA were harvested after 12□h and 24□h, stained with TB, and imaged. **f** Impact of TF3G and GDA on ER stress and cell death marker genes. Relative transcript levels were quantified by qRT-PCR. *ACT2* was used as an internal control. Data represent mean ± SD of three biological replicates. Lowercase letters indicate statistically significant differences among treatments (*P*□<□0.05, one-way ANOVA with Tukey’s HSD test). **g** Theaflavins-induced protein ubiquitination. Total proteins from TF3G- and GDA-treated seedlings were immunoblotted with anti-ubiquitination antibody. Ponceau S staining was used as a loading control.

To further test that TF3G-induced ER stress underlines the onset of PCD, we utilized tauroursodeoxycholic acid (TUDCA), a hydrophilic bile salt and an FDA-approved potent inhibitor of apoptosis in animal cells. It modulates the mitochondrial cell death pathway, inhibits ROS production, and reduces ER stress ^69^. TUDCA represses ER stress-induced cell death by improving protein folding capacity and mitigating protein aggregation. In plants, TUDCA was found to alleviate tunicamycin- and dithiothreitol-induced ER stress and cell death ^69^. Recently, TUDCA has been shown to protect WT Arabidopsis plants from high light stress ^56^ and *sal1* mutant from cadmium toxicity ^70^. As anticipated, the exogenous treatment of TUDCA reduced the impact of GDA and TF3G, showing an alleviated cell death in a dose-dependent manner (Supplementary Fig. 11a-b). This result reinforced that both TF3G and GDA induce a folding stress response leading to cell death, which can be attenuated by promoting protein folding machinery or adding artificial chaperones.

While surveying the literature, we also found an *in silico* study stating that one of the catechins, epigallocatechin gallate (EGCG), having a somewhat similar structure to TF3G, can also potentially inhibit HSP90. Although GTE didn’t show any cell death in our case, since EGCG is a major constituent in GTE, we tested the pure chemical in a dose-dependent manner (50, 100 and 200 µM). In agreement with our GTE data, pure EGCG also didn’t show cell death under used concentrations (Supplementary Fig. 12).

### TF3G induces reprogramming of nuclear gene expression, activating pronounced ER-UPR and a general stress response

We performed a temporal transcriptome analysis to further ascertain the folding stress-induced responses. Cell death analysis revealed that both TF3G and GDA showed an onset of cell death after 12 h of treatment (Fig. 8a). So, we performed RNA sequencing at 6 h (the stage before cell death) and 12 h (the onset of cell death) after treatment. Interestingly, a large number of genes were differentially expressed even before the onset of the cell death in both TF3G and GDA treatments, which coincided and further augmented during the onset of the cell death (Fig. 8b, Supplementary Fig. 13a-c, Supplementary Data 6, 7 and 10). As anticipated, gene expression was significantly overlapped between TF3G and GDA treatments (Fig. 8b, Supplementary Fig. 13a-c, Supplementary Data 6, 7 and 10). Since the transcriptomic changes were almost similar, we emphasized TF3G data only. A total of 1984 genes were induced with at least two-fold upregulation after 6 h, which were augmented to 2420 genes after 12 h; with a significant portion of these genes (1588 genes) overlapping (Fig. 8c, Supplementary Data7-S8). For further analysis, we considered 2816 genes collectively showing upregulation at 6 and 12 h (Fig. 8c, Supplementary Data 8). The GO enrichment analysis of the set of 2816 genes (Supplementary Data9), revealed an overrepresentation of genes involved in biological processes (BP) like response to stimulus, response to stress, defense response, response to wounding, response to hormones, immunity, encapsulating structure organization, secondary metabolism, especially phenylpropanoid metabolic process and root development. In molecular function (MF), genes involved in protein folding, protein binding and dimerization, antioxidant and detoxification response, transcriptional regulation, and transportation were significantly enriched. Similarly, in the case of cellular compartment (CC), genes encoding proteins localized in secretory vesicles (indicative of autophagy), extracellular region and cell-cell junction (indicative of intercellular communication), ER (indicative of ER stress) and vacuole (indicative of cell death) were enriched significantly (Fig. 8d). The GO enrichment analysis depicted activation of a general stress response (Supplementary Data 12), with a notable overrepresentation of typical ER-UPR signatures (Fig. 8d, Supplementary Data 13). The ER-UPR signature genes include those directly related to ER-stress, heat shock factors, proteins and chaperones, genes involved in ubiquitination-mediated proteolysis and autophagy, which exhibited upregulation after 6 h, and genes related to cell death that showed higher expression after 12 h in both TF3G and GDA (Fig. 8e). To ascertain whether the ER-UPR and cell death genes showed a temporal expression, selected genes were analysed from 3 h onwards through qRT-PCR. As indicated in RNA seq data, expression of ER stress markers, *BIP3* and *PDI6* showed a gradual upregulation from 3 h onwards in both TF3G and GDA (Fig. 8f). As HSP90 protein is present in cytosol as well, GDA is known to activate cytosolic folding stress response (also referred to as heat stress response; HSR), we ascertained the activation of master heat shock factor *HSFA2* and key HSR-induced gene *HSP101* (*HOT1*). Though both genes showed a gradual upregulation with time, the expression was much higher in GDA compared to TF3G, indicative of a slight specificity of TF3G towards ER and generalization of GDA towards both ER-UPR and HSR. This observation was reinforced by a higher expression of ER-specific *HSP90.7* in the case of TF3G, and a comparatively augmented upregulation of cytosol-specific *HSP90.1* in the case of GDA (Fig. 8f). Higher expression of *HSP90.7* could be attributed to a relatively higher protein folding and targeting load in ER (endomembrane system), where HSP90.7 is crucial for refolding of misfolded proteins, preventing their aggregation and maintaining ER homeostasis (Marzec et al., 2012). The expression of ER-localized cell death suppressor *BI1-1*, and ER stress-induced PCD and autophagy regulator *Metacaspase 5* (*MC5*) (Fig. 8f), further affirmed this notion. Interestingly, the expression of *BI1-1* was increased until 6 h and then declined; on the contrary, *MC5* showed a gradual upregulation with a drastically heightened expression after 12 h (Fig. 8f), indicating a tight regulation of ER stress-induced cell death onset. These observations indicated a temporal expression of UPR and PCD genes and stated an apparent specificity of TF3G towards ER-UPR.

**Fig. 8:**
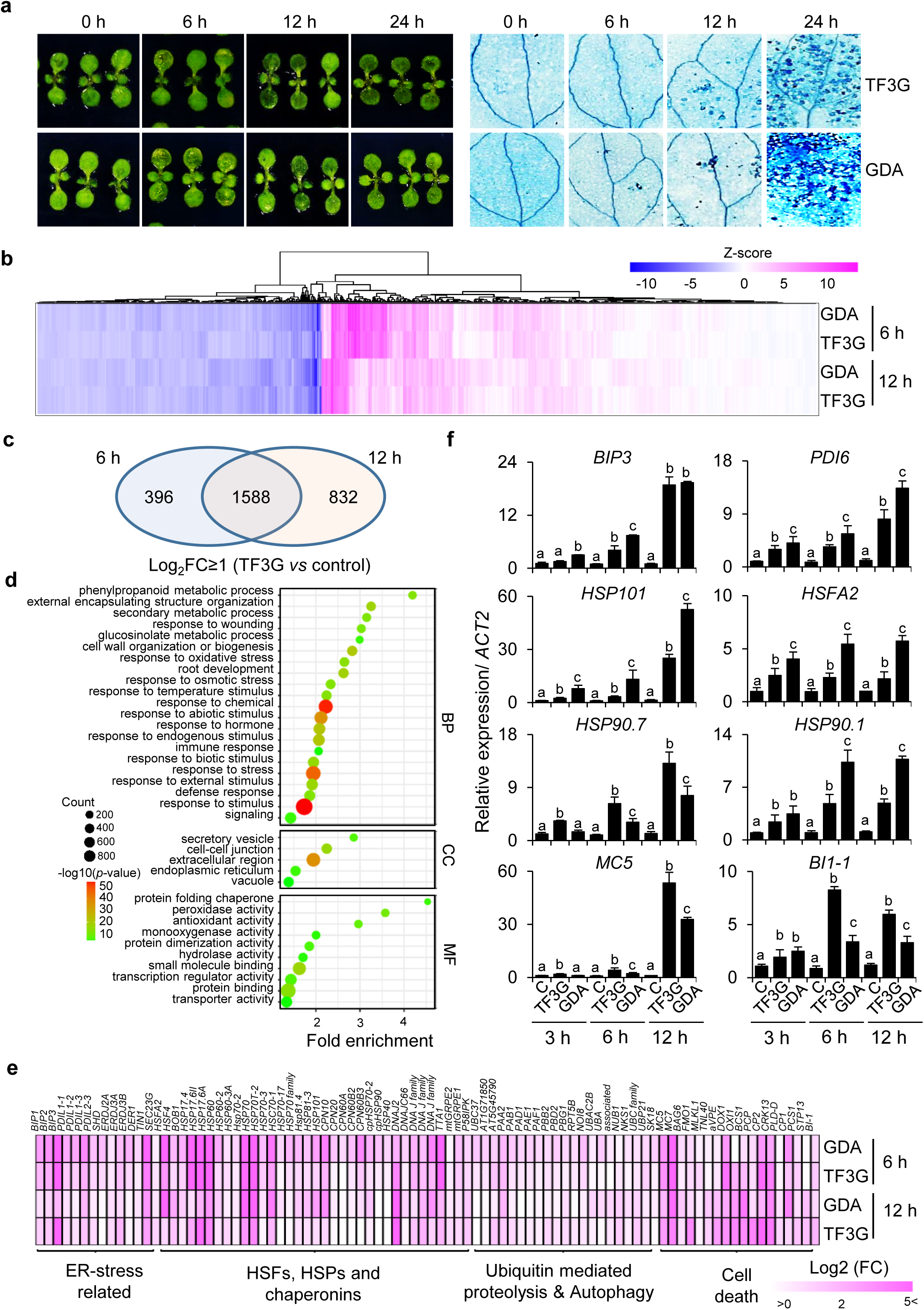
TF3G reprograms the gene expression, mimicking the one induced by drought in Tea. **a** Time-dependent cell death. Five-day-old Col-0 seedlings treated with 200 µM TF3G or GDA were imaged and stained with trypan blue (TB) after 24 h. **b** Heatmap showing global transcriptional changes upon GDA and TF3G treatments at 6 h and 12 h, corresponding to early stress signaling events. Z-scores indicate relative expression, with upregulated genes in pink and downregulated genes in blue. **c** Venn diagram showing overlap and distribution of genes upregulated under TF3G treatment. Differentially expressed genes were selected based on log2(fold change) thresholds. **d** Enrichment bubble plot displaying gene ontology categories significantly enriched among genes upregulated at both 6 h and 12 h TF3G treatments. **e** Heatmap showing the expression of ER stress-related genes upon TF3G and GDA treatments, represented as log2(fold change). **f** Relative transcript levels of key ER stress-related genes. Total RNA was extracted from five-day-old seedlings subjected to TF3G or GDA for 3, 6, and 12 h. Gene expression was quantified using qRT-PCR, with *ACT2* as an internal control. Values represent mean ± SD of three independent biological replicates. Lowercase letters denote statistically significant differences among treatments (*P* < 0.05, one-way ANOVA with Tukey’s HSD test).

In the case of downregulated genes, 1282 genes were repressed after 6 h, whereas expression of 2061 wes declined after 12 h, with an overlap of 934 genes in both time points (Supplementary Fig. 13d, Supplementary Data 10). The GO enrichment analysis of collective 2409 genes revealed an overrepresentation of genes responding to stress, abiotic and biotic stimulus, starvation, and involved in energy metabolism, photosynthesis, and light reactions among BP; genes involved in ion binding, metallopeptidase activity, oxidoreductase activity, tetrapyrrole binding etc, were overrepresented in MF; and similarly, genes residing in chloroplasts, thylakoids, peroxisomes and apoplast were enriched in CC (Supplementary Fig. 13e, Supplementary Data 11). The GO analysis showed that TF3G induces growth inhibition mainly by affecting the energy metabolism, photosynthetic process and cell cycle regulation. Temporal expression analysis of selected genes involved in photosynthesis and photoprotection, namely, *photosystem II subunit S* (*PSBS*), *Violaxanthin De-epoxidase 1* (*VDE1*) and protease *FtsH11*, and cell cycle inhibitor *SIAMESE-RELATED Cyclin-Dependent Kinase* (*SMR7*), reinforced these observations (Supplementary Fig. 13f). These results supported the notion that stress-activated PPO and resulting TFs mediate stress-induced growth inhibition and PCD in Tea plants.

### TF3G-induced response is dependent upon IRE1-bZIP60-driven canonical ER-stress pathway

Upon severe or chronic ER stress, UPR promotes PCD, which kills unwanted cells to protect other cells ^71^. Such a PCD mimics a hypersensitive response observed in immune and abiotic stress responses and is regulated mainly by IRE1-bZIP60 module ^72^. Given the induction of cell death and upregulation of specific bZIP60-dependent genes upon TF3G and GDA treatments, we postulated the involvement of IRE-bZIP60 module. To test this notion, we utilized knock-out (T-DNA) mutants of bZIP17, bZIP28, and bZIP60, which were confirmed through PCR-based genotyping (Supplementary Fig. 14). Interestingly, in line with gene expression data and upregulation of spliced form of *bZIP60* upon TF3G and GDA treatments, the cell death was abolished in the absence of functional bZIP60 protein. The TB staining showed that the cell death was almost comparable in *bzip17* and *bzip28* mutants, whereas it was largely rescued in *bzip60* (Fig. 9a-b). Transcript-based expression of unspliced and spliced forms of *bZIP60* gene (Fig. 9c) confirmed that *bzip60* mutant was indeed knock-out. Besides cell death, expression of bZIP60-dependent genes, *NAC103,* and *BIP3*, remained unaffected in *bzip60* (Fig. 9c). The involvement of bZIP60 was then tested with BTE treatment. As observed with pure metabolites, a similar reduction of bleaching and cell death was also observed upon BTE treatment (Supplementary Fig. 15). Transcription factor binding site analysis on the promoters of 2816 TF3G-induced genes using Find Individual Motif Occurrences (FIMO) v5.5.5 against motif database “ArabidopsisPBM_20140210” with default parameters supported the notion further (Supplementary Data14). Among 2816, 1062 genes contained motifs of either bZIP60_2 (gcCACGTCAy), or bZIP60 (rTGACGTCAy) or both in their promoters (Fig. 8d and Supplementary Data 14). These results specified that, indeed, TF3G-induced response is dependent upon the bZIP60-driven pathway.

**Fig. 9:**
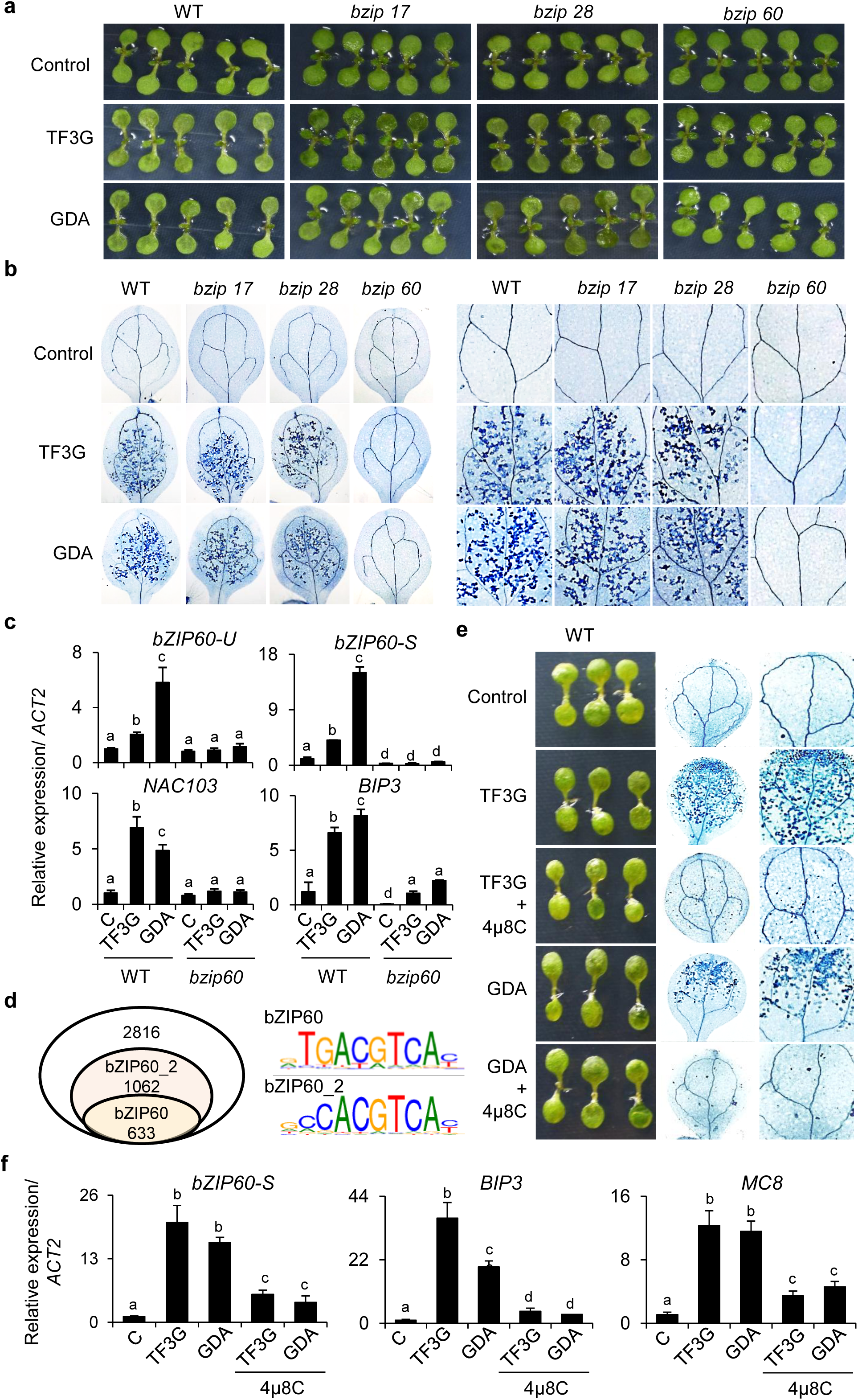
TF3G-induced phenotypes are driven by IRE1-bZIP60-dependent ER stress pathway. **a-b** Macroscopic phenotypes **(a)** and trypan blue (TB) staining **(b)** of five-day-old Col-0, *bzip17*, *bzip28*, and *bzip60* seedlings treated with TF3G or GDA for 24 h. **c** Relative transcript levels of ER stress marker genes determined by qRT-PCR. *ACT2* was used as an internal standard. **d** Overlap of TF3G-induced genes with bZIP60-driven genes. Promoter sequences (3 kb upstream) of 2,816 TF3G-induced genes were scanned using Find Individual Motif Occurrences (FIMO v5.5.5) against the “ArabidopsisPBM_20140210” motif database. A total of 1,062 genes contained the bZIP60_2 motif (‘gcCACGTCAy’) and 633 genes harbored the bZIP60 motif (‘rTGACGTCAy’). **e** Rescue of TF3G-induced cell death by IRE1 inhibition. Five-day-old Col-0 seedlings were treated with TF3G or GDA for 60 min, transferred to liquid MS medium with or without 200 µM 4µ8C, and incubated in growth chambers for 24 h prior to phenotyping and TB staining. **f** Relative transcript levels of ER stress- and cell death-related marker genes in seedlings treated as in **(e)**. Expression was quantified by qRT-PCR with *ACT2* as internal control. Values in **(c** and **f)** represent mean ± SD of three independent biological replicates. Lowercase letters denote statistically significant differences between treatments (P < 0.05, one-way ANOVA with Tukey’s HSD test).

To further affirm the involvement of IRE1-bZIP60 module, we then treated TF3G and GDA together with IRE1 inhibitor 4µ8C, preventing the activation of bZIP60. In agreement with *bzip60* mutant data, inhibition of IRE1-driven activation of bZIP60 resulted in repression of ER stress-induced cell death and gene expression (Fig. 9e-f). These results agree with earlier reports where GDA-mediated selective inhibition of ER-localized HSP90 was shown to activate IRE1 kinase-dependent bZIP60 splicing and downstream upregulation of BIP genes in animals ^66^.

TF3G activates basal immunity and elicits hypersensitive response-like cell death in plants

Transcriptome analysis revealed activation of stress-responsive genes upon TF3G and GDA treatments, where a significant proportion (171 genes) represented “defense response” (Fig. 8d, Supplementary Data 9 and 14). This subset includes key salicylic acid (SA) and jasmonic acid (JA)-related genes involved in basal defense (Fig. 10a). The qRT-PCR-based expression analysis of SA biosynthetic gene *Isochorismate synthase 1* (*ICS1*), its targeted transcription factor *WRKY40* and cognate downstream gene *Pathogenesis related 1* (*PR1*), validated the transcriptome results (Supplementary Fig. 16b). Upregulation of defense genes prior to the onset of cell death indicated their possible involvement in triggering immune response and consequent hypersensitive response (HR)-like cell death. Since HR-like localized cell death is observed upon biotrophic pathogens infection and also both TF3G and GDA activate genes involved in immunity (Supplementary Fig. 16a-b), we then tested whether BTE, TF3G, and GDA play some role in activating defense during plant-pathogen interactions in Arabidopsis. Four-week-old plants of WT Arabidopsis infiltrated with BTE, TF3G and GDA for 24 h, were inoculated with *Pseudomonas syringae* pv. tomato (*Pst*) DC3000. The growth of *Pst* DC3000 was checked 0- and 3-dpi. Interestingly, the bacterial growth was markedly decreased in both BTE and pure chemicals pre-infiltrated samples compared to mock controls (Supplementary Fig. 16c-d). These results clearly indicated that both Tea metabolite TF3G and bacterial metabolite GDA, interfere with folding machinery and induce HR-like localized cell death. To check whether these metabolites are general elicitors of cell death, we treated seedlings of tomato and tobacco (*N. benthamiana*) with TF3G and GDA. The TB staining results demonstrated that both HSP90 inhibitors activate cell death in tomato and tobacco leaves (Supplementary Fig. 16e-f). These results demonstrated that theaflavins-rich BTE, especially pure TF3G, elicits HR-like cell death, which can prime immune response. This notion is well supported by the fact that many NLR and immunity-related proteins are clients of HSP90 in ER ^73^, and TF3G-HSP90 interaction promoted expression of ER-specific HSP90.7, likely stimulates immune response. Not only BTE and TF3G, but it seems general HSP90 inhibitors, as depicted in case of GDA, can also prime immunity in plants and perhaps in animals as well if applied in a specific optimized dose.

**Fig. 10.**
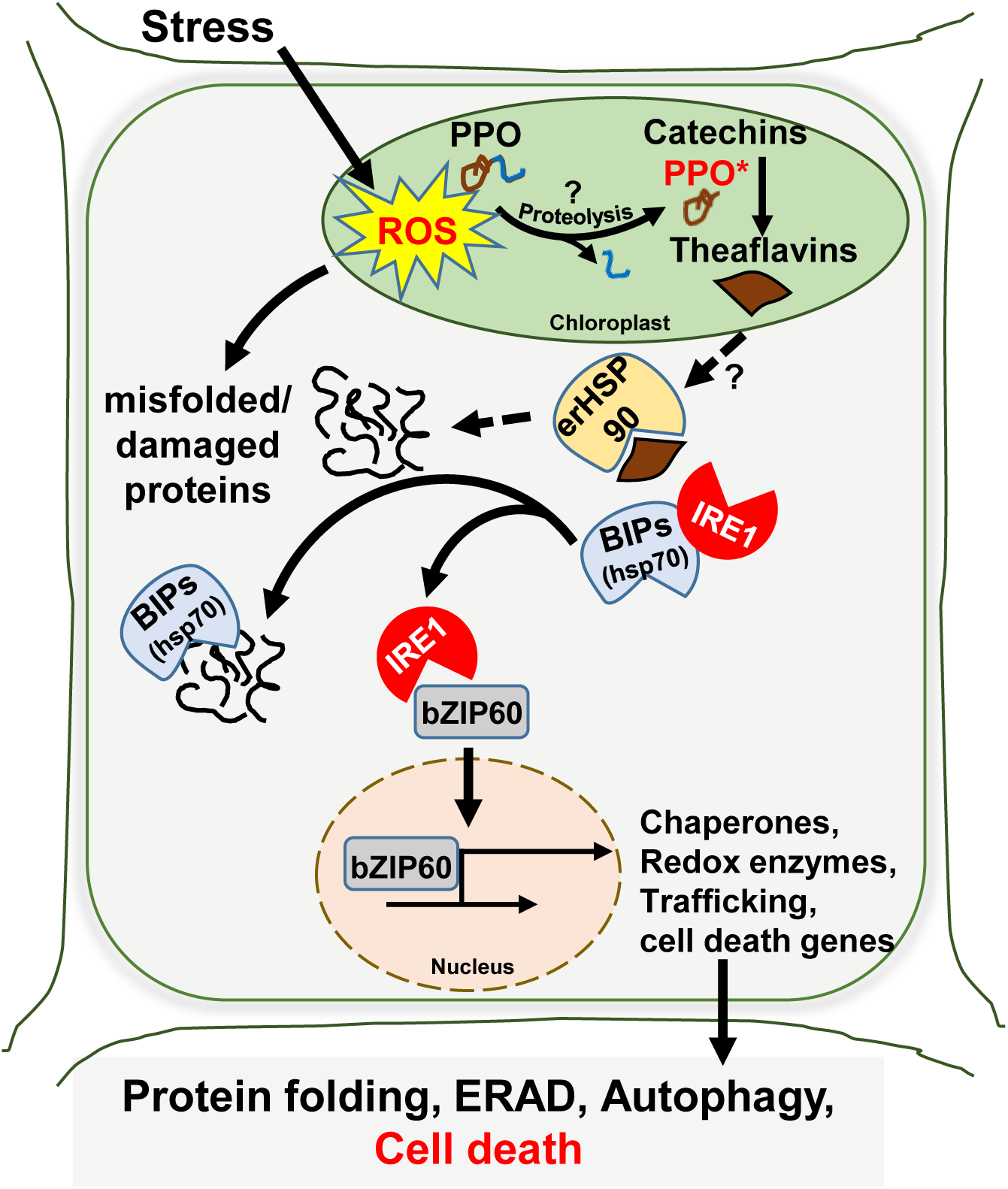
Proposed model depicting how PPO senses stress and triggers TFs-mediated stress signaling. Drought-induced oxidative stress results in proteolytic activation of PPO (through an unknown enzyme) in Tea plants, leading to heightened production of oxidized flavonoid metabolites, particularly theaflavins. The theaflavins, especially TF3G, likely move into the cytosol and ER, where it interacts with HSP90 proteins, augmenting the accumulation of unfolded or misfolded proteins in these sub-cellular localizations. Accumulation of unfolded client proteins disintegrates the BIP(HSP70)-IRE1 complex in ER to cater to client proteins and activate UPR through the IRE1-bZIP pathway. The cognate signaling pathway activates chaperones, redox enzymes, and cellular trafficking genes, which improve protein folding or activate ERAD and autophagy to maintain proteostasis. In case, the folding stress is severe, a controlled and programmed cell death is activated to sustain the adversity of stress.

## Discussion

Due to their sessile nature, plants must continuously adapt to biotic and abiotic stresses, relying on complex physiological and biochemical responses. Stress perception in plants triggers metabolic and signaling cascades, activating specific enzymes and accumulating metabolites ^74^. Chloroplast, an indispensable organelle housing photosynthesis and various metabolic pathways, is susceptible to biotic and abiotic stresses. In response, it activates distinct retrograde signaling pathways involving enzyme-metabolites interplay where chloroplast-derived metabolites such as methylerythritol cyclodiphosphate (MEcPP), 3’-phosphoadenosine 5’-phosphate (PAP), Mg-protphyrin IX, etc. influence nuclear gene expression, ensuring the plant’s adaptation and survival ^19, 20^. In many plants, chloroplast contains PPO enzymes, which catalyze the oxidation of phenolic compounds into *o*-quinones ^46, 50^. In various species such as Tea, tomatoes, potatoes, apples, and walnuts, PPO has been shown to induce browning, deter herbivory, fortify cell walls against pathogens, and may induce cell death ^49, 51, 75^.

Our results revealed that drought induces a proteolytic activation of PPO in Tea plants, leading to enhanced production of oxidized flavonoid metabolites, particularly theaflavins (Figs 1-2, Supplementary Fig. 1). The activation of PPO coupled induction of an ER stress-like phenotype, in which genes involved in typical ER-UPR and PCD were markedly upregulated (Fig. 3). This signature was stress-intensity dependent and augmented in susceptible genotypes, which showed higher PPO activity and TFs accumulation (Fig. 3). Silencing and overexpression of PPO resulted in repression and augmentation, respectivey, of downstream signalling (Figs. 4 and 5). The exogensous feeding of TFs-rich BTE or pure theaflavins showed onset of an ER stress response with augmented activation of ER stress and cell death genes (Fig. 5). These findings reinforced that stress-induced proteolytic PPO activation triggers the stress signaling via its catalytic producs (TFs). Similar stress induced proteolytic activation of PPO and activation oif cognate genes in other plants, including tomato and wheat (Supplementary Figs 7-8), supports the notion that this pathway is not limited to Tea and operates in many other plant species. These results also reinforced earlier findings where activation of PPO was implicated with defense and resistance to stress factors in both plants and fungi ^50, 76^. The results unveiled a long standing question odf PPO activation and suggests the importance of this evolutionarily conserved enzyme as a an important stress sensor.

Pharmacological treatments-based approach in PPO-deficient plant Arabidopsis, further validated the intertwining of PPO activation and revelead mechanism underlining the augmented UPR. Treatment of TFs-rich BTE and pure TFs (markedly TF3G) induced typical ER stress-like signature phenotypes, including growth inhibition, activation of chaperones, and cell death genes ^77, 78, 79^, in a dose- and time-dependent manner in Arabidopsis (Fig.s 6-7, Supplementary Fig. 9). These results resembled the general stress response with a heightened ER stress response observed in the case of MEcPP-driven retrograde signaling ^20^. The TF3G likely moves into the cytosol and ER, where it interacts with HSP90 proteins (Fig. 7), augmenting the accumulation of unfolded or misfolded proteins in these sub-cellular localizations. Accumulation of unfolded client proteins disintegrates BIP(HSP70)-IRE1 complex to cater to client proteins and activate UPR through the IRE1-bZIP pathway (Fig. 9 and 10, Supplementary Fig. 11). These results were reinforced by a known HSP90 inhibitor GDA, which showed almost similar phenotypes of growth inhibition, ER stress, and cell death (Fig. 7-10, Supplementary Fig. 11-S13). Interestingly, although GDA is routinely used for inhibiting HSP90 proteins in plant studies, the cell death phenotype has not been reported before. Our results also demonstrated that, like GDA ^66^, TF3G inhibits HSP90 in both cytosol and ER; however, its action seems more in ER, possibly due to its higher availability in this organelle. The presence of membrane contact sites between chloroplasts and ER ^80^ could be the possible route for TF3G transport from the chloroplast to the ER. In addition, a significant overrepresentation of genes localized in plasmodesmata and cellular junctions (Fig. 8) indicated the possible routes for inter-cellular transport of TFs. Further investigations in the future on transport, mode, and site of TFs’ actions would certainly enhance our understanding of this inter-organellar and inter-cellular communication linked to cellular adaptation. The stress-dependent activation of PPO and activation of ER stress-like response in Tea (Fig. 1-3), which is apparently demonstrated in a dose and time-dependent manner in Arabidopsis (Fig. 6-8, Supplementary Fig. 9, S11-12), suggests that under mild stress, PPO-triggered and TF-mediated signaling activates UPR, which activates PQC machinery involving HSPs, HSFs, chaperones, UPS- and autophagy-related genes, to maintain cellular proteostasis. However, with increased stress intensity, which is proportional to PPO activity and TFs generation (Fig. 1), the ER stress then activates cell death signaling pathways ^5, 8^, sacrificing weaker cells to nourish other cells and sustain stress conditions.

Since the ER stress-associated cell death pathway overlaps with defense pathway, TF3G/GDA treatments can prime immune responses and HR-like cell death in plants (Supplementary Fig. 16). These results showed the potential applications of TF3G and GDA as defense/stress priming agents, which can elicit synthesis of protective secondary metabolites *in vivo*, and *in vitro* cell culture systems. Given that TFs accumulation is higher in drought-susceptible plants and that Tea is prone to blister blight, a fungal disease ^81^, it would be intriguing to see any link between the responses to drought and blight in the Tea germplasm, which could help develop strategies against these serious issues in global Tea cultivation. Besides, heat stress, intensified by global warming and climate change, is another inclusive threat to Tea cultivation ^82^. Since HSR significantly overlaps with UPR, and as heat also directly impacts chloroplast redox state, heat stress might also activate the PPO-TFs module to augment HSR/UPR. This notion is supported by a recent finding where HSP90 was found to be the key regulator of HSR, and its inhibition was found to promote thermotolerance in Tea ^83^. Germplasm-based analysis of such interconnection between the multitude of stresses and the PPO-TFs module could be explored to develop sustainable management of Tea cultivation around the globe.

As PPO is present in many plant species and is also found in bacteria and fungi ^50, 76^, the signaling mediated by its catalyzed products, *o*-quionones, seems to be evolutionarily conserved (Supplementary Fig. 7-8) and should be looked upon for exploring its implications and utility in different plants and microbes. In summary, the present study revealed the role of chloroplast-localized PPO as a stress sensor and an activator of stress signaling pathways through its catalyzed product TFs under drought stress. Further research to understand its activation mechanism and TFs’ downstream functioning would pave the way for genetic manipulations of this module to manage various stress factors in Tea and other plants.

## Methods

### Plant materials and growth conditions

Twelve genotypes of Tea [*C. sinensis* (L) O. Kuntze] were used in the study. These genotypes are maintained at CSIR-Institute of Himalayan Bioresource Technology Palampur, Himachal Pradesh (1,300□m Altitude, 32°06′N, 76°33′E). Shoot cuttings of the genotypes were collected during the actively growing period (April, 2023; forenoon) in de-ionised water and acclimatized in a growth chamber for 24 h under long-day conditions [(16 h light (200 µmol□m^-2^□s^-1^)/8 h dark; at 25±1□°C)] with a relative humidity of 70±5 %. After that, the cuttings were transferred to half-strength Hoagland’s No. 2 basal salt mixture medium (Himedia, India) containing 10% polyethylene glycol 8000 (PEG 8000; Promega, USA) and incubated for 48□h. For downstream analyses, three leaves and a bud were harvested for each genotype after 0, 24, and 48□h.

All *Arabidopsis thaliana* seeds used in this study were derived from Columbia-0 (Col-0) ecotype and were harvested from plants grown under long-day conditions (16 h light/8 h dark; 100 µmol□m^−2^□s^−1^ with a relative humidity (70±5 %) at 22□±□2□°C). Mutants used in this study, namely, *bzip17*-1 (SALK_104326), *bzip28*-2 (SALK_132285C), and *bzip60*-1 (SALK_050203C), were obtained from the Nottingham Arabidopsis Stock Centre (NASC) and the genotypes were confirmed through PCR-based analyses. The primers used in this study are listed in Supplemental Supplementary Dataet 16. Seeds were sterilized in a 20% hypochlorite solution followed by five washes with sterile water and sown on half-strength Murashige and Skoog (MS) medium (Duchefa Biochemie) with 0.5% (w/v) Suc and 0.7% (w/v) agar. The seedlings were grown in a growth chamber under continuous light conditions (µmol□m^-2^□s^-1^) and relative humidity (70±5 %).

Four-week-old *Nicotiana benthamiana* plants grown in a controlled growth chamber under long-day conditions (16 h light/8 h dark) at 25 °C were used for all transient assays (visualization, immunoblot analysis, and chloroplast fractionation assay).

### Chemical treatments

Black Tea extract (BTE; Cat. no. T5550), tauroursodeoxycholic acid (TUDCA; Product no. 580549) and Epigallocatechin galate (EGCG; Cat. no. E4143) were purchased from Sigma Aldrich, USA. Green Tea extract (GTE) was prepared in-house at CSIR-IHBT, Palampur. Geldanamycin (Item no. 13355), Theaflavin (Item No. 25129), Theaflavin-3-gallate (Item No. 31510), and Theaflavin 3,3’-digallate (Item No. 25215), were purchased from Cayman Chemicals, USA.

For pharmacological treatments, 5-day-old seedlings grown on MS Agar plates were used. In the case of BTE and GTE treatments, seedlings were submerged in the solution and subjected to 5 min vacuum infiltration followed by 55 min incubation at room temperature (RT) in a laminar airflow. Subsequently, chemicals were discarded, and plates were transferred to a growth room, maintaining controlled conditions. On the indicated day of post-treatment, seedlings were harvested for scoring phenotypes, cell death staining, and RNA and/or protein isolation. In case of theaflavins and EGCG treatments, seedlings grown on MS Agar plates were transferred to 6-well cell culture plate containing chemical solutions. The seedlings were subjected to 5 min vacuum infiltration followed by 55 min incubation at RT. After incubations, chemicals were replaced with liquid MS media and transferred to the growth room on a platform with gentle shaking for uniform nutrient absorption. Samples were collected for phenotypes, cell death staining, and RNA isolation after desired time periods.

### Photosystem II (PSII) quantum efficiency (Fv/Fm) analysis

At least five seedlings of Arabidopsis and three third leaves of Tea were used for Fv/Fm analysis using FluorCam 800. Samples were subjected for 10 min dark, followed by an exposure to a flash of actinic light to measure photosystem II efficiency (Fv/Fm) using FluorCam 800 (Photon System Instruments). The maximal quantum yield of PSII (Fv/Fm) was determined from equation; Fv/Fm = (Fm - Fo)/Fm, where Fo is the initial minimal fluorescence on dark-adapted leaves for 10 min and Fm is maximal dark-adapted fluorescence.

### Relative water content (RWC), Relative electrolyte leakage (REL)

To measure the relative water content, the third leaf of Tea shoot was taken after 24 and 48 h of PEG-treatment. After weighing for FW, leaves were completely submerged in 100 ml de-ionized water and incubated at 4 ^°^C overnight. Next morning, excess water on the leaf’s surface was removed using paper towels, followed by weighing to record turgid weight (TW). Leaves were then dried in a hot air oven at 70 ^°^C and weighed to determine dry weight (DW). RWC was calculated using following equation: RWC (%) = (FW-DW)/(TW-DW) × 100 Relative electrolyte leakage (REL) was determined to assess the relative quantity of dead cells in response to stress conditions. Three leaf discs of 5 mm diameter were excised from third leaf and incubated in de-ionized water with gentle shaking for 2 h. Electric conductivity was estimated before (BA) and after autoclaving (AA), using a conductivity meter (Oakton PC700). The REL was calculated using the following equation: REL (%) = (BA/AA) × 100

### Lipid peroxidation assay

Malondialdehyde (MDA) content was determined to quantify lipid peroxidation. Briefly, lyophilized powder (0.05 g) of third leaf was homogenized with trichloroacetic acid solution (0.1% w/v), followed by centrifugation at 10,000 × *g* for 10 min. The supernatant was mixed with an equal volume of 20% TCA containing 0.5% (w/v) thiobarbituric acid and incubated at 90 °C for 1 h followed by cooling. The resultant mixtures were centrifuged at 10,000 × *g* for 5 min, and the supernatant was saved. MDA contents were determined spectrophotometrically by recording absorbance at 450, 532, and 600 nm. MDA contents were determined on fresh weight (FW) basis using following formula:

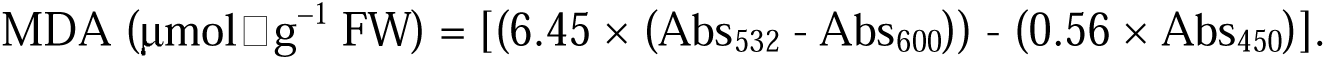

### Polyphenol oxidase (PPO) activity assay

The PPO assay was performed according to Singh and Ravindranath ^84^, with some modifications. Briefly, 0.5 gm of third leaf tissue was grounded in liquid nitrogen and treated with chilled acetone until the powder became white and free from phenolics. The white powder was dried under a vacuum to remove acetone. The dried powder was homogenized in 400 µl water and centrifuged at 400 × *g* for 10 min. The pellet was homogenized with 300 µl 0.2 M Na_2_SO_4_, followed by centrifugation at the same speed. The resulting supernatant was used as enzyme extract for PPO activity assay. The protein content in the enzyme extract was estimated using BCA protein assay kit (Purgene). The PPO activity was measured spectrophotometrically in a UV-treated 96-well plate. The reaction mixture contained 12.5 µl of 5 mM catechin (substrate), 175 µl of 0.2 M Na_2_SO_4_ and 12.5 µl of enzyme extract. Absorption was recorded every 3 min for 15 min. One unit of enzyme activity is defined as the rate of change of 0.001 absorption unit per min at 380 nm. The specific activity of PPO activity was expressed in unit mg^-1^ protein min^-1^.

In tomato and wheat, leaf tissues were homogenized in 500 µL of 0.1 M sodium citrate buffer (pH 5.0) at 4°C. The homogenate was centrifuged at 10,000 rpm for 5 min at 4°C, and the protein content in the enzyme extract was quantified using the BCA protein assay kit (Purgene). PPO activity was measured in a 96-well plate using a reaction mixture consisting of 150 µl of 0.1 M sodium phosphate buffer (pH 6.5), 20 µl of enzyme extract, and 30 µl of 0.01 M catechol as the substrate. The reaction was monitored by measuring absorbance at 505 nm, and enzyme activity was expressed as absorbance (units) min□¹ mg□¹ of protein.

### Metabolite analysis

The third leaves from the drought-subjected cuttings were flash-frozen in liquid nitrogen to preserve metabolite integrity. The frozen samples were then pulverized into a fine powder using a pre-chilled mortar and pestle. Subsequently, the powdered samples were lyophilized overnight. Dried samples weighing 100 mg were transferred into fresh 2 ml microcentrifuge tubes and resuspended in 70% HPLC grade methanol. The mixtures were subjected to sonication for 20 min in a bath sonicator at ambient temperature, followed by centrifugation at 13,000 rpm for 10 min. The resulting supernatants were filtered through a 0.22 µm polyvinylidene fluoride filter (Millipore, USA), achieving a final concentration of 20 mg/ml. Finally, 200 µl aliquots of the filtered extracts were transferred to high-recovery vials (Agilent, USA) for subsequent analytical assessment.

The selected Tea flavonoids were quantified using a UPLC H-class system equipped with a photodiode array detector (PDA) (Waters Acquity, USA). Standard stock solutions were prepared at a concentration of 1 mg/mL in LC-MS grade methanol, syringe filtered, and stored at 4 °C. The calibration curve was then prepared using working solutions with concentrations ranging from 2 to 200 ppm. Two microliters of standards and samples were loaded onto the Zorbax Eclipse Plus RRHD C18 column (2.1 × 150 mm, 1.8 µm particle size) (Agilent, USA). The mobile phase consisted of 0.05% trifluoroacetic acid in water (solvent A) and 0.05% trifluoroacetic acid in HPLC-grade acetonitrile (solvent B). The gradient started with 6% of solvent B at 0 min, held constant until 0.5 min, followed by an increase to 28% of solvent B from 0.5 to 8 min. The proportion of solvent B was further increased to 75% from 8 to 10 min. Subsequently, the mobile phase composition was held at 45% of solvent B from 10 to 11.5 min. At 11.51 min, the conditions were reverted to the initial 6% of solvent B, which was maintained until the end of the 16-min run. The flow rate was set to 0.28 mL/min, and detection was carried out at a wavelength of 270 nm. Phenolic acids and flavonoids were identified by assessing the absorption at 270 nm and retention time and quantified by comparing the area of the peaks against the calibration curve based on the retention time and UV spectra at 270 nm.

For quantifying theaflavins, 2 µl of standards and samples and standard mix were loaded onto a BEH-HSS-T3 C19 column (2.1 x 100 mm, 1.8 µm particle size) in an LC-MS system equipped with PDA and SQ MS detector (Waters Acquity, USA). Mobile phase used solvent A: 0.1%HCOOH in water and solvent B: 0.1% HCOOH in Acetonitrile. The gradient was run for 33 min with 95% of A and 5% B, at 0 min, held constant until 0.5 min, followed by an increase to 26% of solvent B from 0.5 to 26 min. The proportion of solvent B was then decreased to 15% from 26 to 29 min, and then to 5% from 30 to 33 min. The flow rate was set to 0.3 −0.35 mL/min, and detection was carried out at a wavelength of 275 nm. Stock solutions (5 mM) of TF, TF3G and TFDG were prepared in methanol. Appropriate dilutions were made in assay buffer to test the molecules at various concentrations keeping methanol concentration less than 1% in the final reaction volume. Theaflavins were confirmed with peak mass and quantified using UV spectra. MS scan were run from 100-1800 Da in a positive ion mode.

### Transient expression and localization of Tea PPO in *N. benthamiana*

The stop codon-less cDNA of Tea PPO gene (*CsPPO2,* Gene Id: ACN38060.1) was cloned into the pDONR221 Gateway vector (Invitrogen, ThermoFisher Scientific, USA) through the BP clonase reaction (Invitrogen, ThermoFisher Scientific, USA) and subsequently recombined into the Gateway-compatible plant binary vector pGWB505 for C-terminal fusion with sGFP through the LR clonase reaction (Invitrogen, ThermoFisher Scientific, USA) ^85^. The generated vectors were transformed into Agrobacterium tumefaciens strain GV3101. The suspensions of agrobacteria carrying different constructs were infiltrated into healthy leaves of 4-week-old *N. benthamiana* plants and analyzed after 48 h. The PPO-GFP signals and chlorophyll autofluorescence were visualized using a confocal laser scanning microscope (TCS SP8; Leica Microsystems). Images were acquired and processed using LAS AF Lite software version 2.6.3 (Leica Microsystems).

### Subcellular accumulation of Catechins

Subcellular accumulation of catechins was affirmed using Vanillin-HCl staining, according to Liu, Gao ^45^. Briefly, the transverse hand sections of the third leaf from Tea cuttings were incubated in 0.1% (w/v) Vanillin-HCl solution for 5 min, followed by destaining in milli-Q water. Sections were observed under the bright field in Axioscope 5 microscope (Zeiss, Germany).

### Chloroplast Isolation, Protein extraction and immunoblotting

Intact chloroplasts were isolated from healthy leaves of Tea plants of the UPASI09 genotype subjected to PEG-induced drought stress, as described previously in Dogra, Duan ^4^. To extract proteins, intact chloroplasts corresponding to equal amounts of chlorophyll (200 mg) were lysed in HM buffer (10 mM HEPES-KOH, pH 8.0 and 5 mM MgCl_2_) to a final concentration of 1 mg/ml of chlorophyll. The chloroplast suspensions were then processed to remove chlorophyll as previously described by Wang et al. (2016), and resuspended in 1× Laemmli SDS sample buffer. All proteins were denatured for 10 min at 95 °C. Equal amounts of proteins were separated on 10% SDS-PAGE and blotted onto Immun-Blot polyvinylidene fluoride membrane (Bio-Rad, USA). The PPO proteins were detected using rabbit anti-PPO (1:1000, catalog no. MBS9458498; MyBioSource, Inc.) and ECL prime western blotting detection reagent (Cytiva Amersham, Fisher scientific).

### RNA Extraction and QRT-PCR

For Tea samples, total RNA was extracted using the *iRIS* method ^86^. In Arabidopsis, RNeasy mini kit (Qiagen, Germany) was used to extract total RNA. The isolated RNA was quantified on Nanodrop 2000 (ThermoFisher Scientific, USA). One microgram of total RNA was reverse transcribed using Verso cDNA synthesis kit (ThermoFisher Scientific, USA). The qRT□PCR was performed using Maxima SYBR Green qPCR Kit (ThermoFisher Scientific, USA) on a CFX Opus 96 Real□Time PCR System (Bio-Rad, USA). The relative expression of target genes was calculated by the comparative 2^−ΔΔCt^ method and normalized to the transcript levels of *GLYCERALDEHYDE-3-PHOSPHATE DEHYDROGENASE* (*GAPDH*) for Tea, *ACTIN5* (NM_001308447.1) for Tomato, and *ACTIN2* (*ACT2; AT3G18780*) for Arabidopsis. The primers used in this study are listed in Supplemental Supplementary DataetS1.

### Virus-induced gene sliencing in Tea

Full-length coding sequences of *PDS* and *PPO* genes were retrieved from the Tea Plant Information Archive (TPIA) database. These sequences were then amplified using primers designed according to ‘pssRNAit’ (https://www.zhaolab.org/pssRNAit/Analysis.gy) tool from cDNA of TV03. The amplified fragments were ligated into the *pTRV2* vector using *EcoRI* and *BamHI* restriction sites. The ligated products were transformed into *E. coli* DH5α, and positive clones were identified by PCR and subsequent sequencing. Plasmids isolated from positive clones were subsequently transformed into *Agrobacterium tumefaciens* GV3101. The infection solution was prepared according to Peng, Xue ^55^. Briefly, the *Agrobacterium* cell were harvested when the culture reached an OD□□□ of 1.0 and resuspended in the infiltration solution (full strength MS media containing 2M 6-BA, 2M Acetosyringone, 100 μM naphthalene acetic acid, and 16 g/L MgCl_2_ at pH 5.8). The *pTRV1* was mixed with *pTRV2* (as a control) and *pTRV2-PDS or pTRV2-PPO* plasmid in 1:1 ratio, followed by incubation at 28 °C for 1 h. Healthy shoot cuttings of TV03 were placed in a beaker containing Agrobacterium cells and subjected to vacuum infiltration at 0.8 kPa for 5 min, followed by injection at petiole sites. The cuttings were then incubated in a half-strength Hoagland’s No. 2 basal salt mixture medium. The infiltrated shoot cuttings were kept dark in a plant growth chamber at 25 °C and transferred to 16 h light/8 h dark cycle at 25 °C and a relative humidity of 60-70% until noticeable phenotypic changes appeared. The leaves were then flash-frozen in liquid nitrogen for further analysis.

### Visualization of Cell Death

Cell death was visualized by trypan blue (TB) and Evans blue (EB) staining. For TB staining, plant cotyledons were submerged in TB staining solution (0.02 g of trypan blue and 10 g of phenol dissolved in 30 mL of glycerol, lactic acid, and water [1:1:1]; this solution was further diluted with ethanol in 1:2 [v/v]) and boiled for 2 min. After a 16-h incubation at room temperature, nonspecific staining was removed with the destaining solution (250 g of chloral hydrate dissolved in 100 mL of water, pH 1.2). Plant tissues were then kept in 10% (v/v) glycerol for imaging. In case of EB staining, cotyledons were submerged in 1 mL of a 0.1% (w/v) EB solution and incubated for 16 h at room temperature. The cotyledons were then thoroughly washed three times with water to remove unbound dye. Images were acquired and processed using LAS software version 4.2.0 in S9i microscope (Leica Microsystems, Germany). For Quantification of EB stained cells, the dye bound to dead cells was solubilized in 50% (v/v) methanol containing 1% (w/v) SDS for 8 h. Cell death was quantified spectrophotometrically by measuring the ratio of absorbance at 600 and 680 nm (*A*_600_/*A*_680_) of the resulting dye solution.

### Protein structure modeling and Molecular docking simulations

The protein sequences of HSP90 were obtained from The Arabidopsis Information Resource (TAIR). Domains and motifs in the protein sequences were deduced using CDD tool at NCBI and 3D structures were retrieved from Protein Data Bank (PDB) (Berman et al., 2000), which were used as templates for the homology modeling approach. Modeler v9.20 (Webb and Sali, 2016) was used to model the 3D structures for the retrieved sequences. The DOPE and Z-scores (Zhang and Skolnick, 1998) were used by default to evaluate the models. The modeled structures were then validated using various software tools, including ProSAweb server (https://prosa.services.came.sbg.ac.at/prosa.php) and PROCHECK (Laskowski et al., 1993), which are stand-alone packages. Additionally, GROMACS 2018 (Van Der Spoel et al., 2005) was used to minimize the free energy and enhance the stereochemistry of the model. Finally, RaptorX (https://prosa.services.came.sbg.ac.at/prosa.php) and COACH (Zhang, 2008) were utilized to identify binding sites and assess conserved domains.

Molecular docking simulations were employed to predict the interactions of HSP90 with theaflavins and GDA, using Autodock tools (Morris et al., 2009) and Autodock Vina (Trott and Olson, 2010) software. The docking process was performed in two steps: (a) evaluation of conformation pattern of the ligand within the active site of the protein and (b) forecasting the binding affinity between the binding site and the appropriate conformation of the ligand. The results of the docking simulation were visualized and assessed using Pymol, while DS Visualizer (https://www.3ds.com/products/biovia/discovery-studio/visualization) was used to plot 2-dimensional interactions within the binding sites.

### RNA-Seq Library Construction and Data Analysis

Total RNA was extracted from three independent biological replicates of the 5-day-old wild-type seedlings treated with TF3G and GDA for 6 and 12 h. The RNA extracted using the RNeasy Plant Mini Kit (Qiagen) was subjected to on-column DNase digestion with RNase-free DNase Set (Qiagen) according to the manufacturer’s instructions. The purity of RNA was verified by a Nano Photometer spectrophotometer (IMPLEN). The Qubit RNA Assay Kit in Qubit 2.0 Fluorometer (Life Technologies) was used to measure RNA concentration. The RNA Nano 6000 Assay Kit of the Bioanalyzer 2100 system (Agilent Technologies) was used to evaluate RNA integrity. Samples with RNA integrity (RIN) ≥8 were used for constructing libraries using TruSeq RNA stranded sample prep kit (Illumina, USA) as per the manufacturer’s instructions. The libraries were pooled at 400□pmol concentration, denatured, and sequenced on an Illumina NovaSeq 6000 system (Illumina) to generate 101□bp paired□end reads. The raw reads of each sample generated from RNA-seq libraries were examined for quality performance using the bioinformatics tool fastp ^87^. Clean reads were further screened for rRNA contamination using a bioinformatic tool RiboDetector (https://github.com/hzi-bifo/RiboDetector). The reads aligned with the RiboRNA sequences were eliminated ^88^. The clean reads obtained were mapped to the Arabidopsis genome (TAIR10) using hierarchical indexing for spliced alignment of transcripts (HISAT2) ^89^. StringTie2 ^90^ is used to assemble the align reads in a reference-guided manner for each sample and extract the potential transcript and their expression level using general feature format (gff3) file (https://ftp.ensemblgenomes.ebi.ac.uk/pub/plants/release56/gff3/arabidopsis_thaliana/Arabid <u>opsis_thaliana.TAIR10.56.gff3.gz</u>). It employs efficient algorithms for the abundance estimation from the aligned read to the reference genome. The reference and assembled transcripts for all samples were merged by StringTie ^90^. Each gene’s raw integer read counts across all samples generated and used for the downstream analysis. Principal component analysis (PCA) was widely adopted method in the RNAseq datasets. The PCA plot was generated by the ggfortify ^91^ package in R using gene expression values of each sample. Differentially expressed genes were identified using R package edgeR, which uses counts per gene in different samples and performs data normalization using the trimmed mean of M-values method. The genes with low expression across the samples might not have enough evidence for differential gene expression. So, genes with CPM values greater than one in at least two samples are candidates for downstream study. The Trimmed Mean of M-values (TMM) normalization method was used for between-sample normalization to normalize the expected counts for relative expression and adequate library size. The genes with at least a twofold change in expression and a false discovery rate of less than 0.05 were considered to be differentially expressed. The transcripts per million values were used to build the expression matrix, and the subsequent clustering and visualization were done using Multi-Experiment Viewer 4.9.0. GO enrichment analysis of differentially expressed genes was performed using the Generic GO Term Finder tool (http://go.princeton.edu/cgi-bin/GOTermFinder), with a significance of *P* < 0.05.

### Accession Numbers

Sequences of the Aranoidopsis genes studied in this article can be found in TAIR database (https://www.arabidopsis.org) under the following accession numbers: *ACT2* (At3g18780), *NAC103* (At5g64060), *OXI1* (At3g25250), *ERDJ3B* (At3g62600), b*ZIP17* (At2g40950), b*ZIP28* (At3g10800), *bZIP60* (At1g42990), g*VPE* (At4g32940), *MC8* (At1g16420), *ERO1* (At1g72280), *NRP1* (At5g42050), *NRP2* (At3g27090), *BIP3* (At1g09080), *HSP70* (AT4G16660), *HSFA2* (AT2G26150), *HSP101* (AT1G74310), *PSBS* (AT1G44575), *VDE* (AT1G08550), *WRKY40* (AT1G80840), *PDI6* (AT1G77510), *SMR7* (AT3G27630), *PR1* (AT2G14610), *BI1* (AT5G47120), *FTSH11* (AT5G53170), *ICS1* (AT1G74710), *HSP90.7* (AT4G24190), *HSP90* (AT5G52640), *MC5* (AT1G79330). Sequences of the Tea genes studied in this article can be found in NCB database (https://www.ncbi.nlm.nih.gov) under the following accession numbers: *PAL1* (KY615669), *PAL2* (KY615671), *CHS* (KC357707), *ANS* (KY615704), *ANR* (KY615701), *PPO2* (MK977643), *BAG2* (XM_028244308.1), *BIP2* (XM_028210952.1), *bZIP60* (XM_028262224.1), *CRT* (XM_028216501.1), *DER2*.2 (XM_028239771.1), *HSFA2* (XM_028252153.1), *NAC103* (XM_028268369.1), *PDI2*.3 (XM_028251828.1), *PDI1* (XM_028226933.1), *RMA3* (XM_028200165.1), *TOR* (XM_028205854.1), *GAPDH* (XM_028237220.1). Sequnences information of Tomato genes studied in this article can be retrived from NCBI database (https://www.ncbi.nlm.nih.gov) having following accession numbers: *ACT7* (NM_001308447.1), *BIP* (XM_004234937.5), *PDI* (XM_004241984.5), *CNX* (NM_001247200.2), *IRE1* (XM_004238397.5), *BI-1* (XM_004238421.4), *bZIP60* (XM_004238421.4), *MC5* (XM_004230556.5), *MC8* (OR255958.1).

## Acknowledgments

The authors would like to thank the Director, CSIR-Institute of Himalayan Bioresource Technology, Palampur, for providing infrastructural facilities. Furthermore, the authors are thankful to Dr. Mohit Swarnkar, Core Facility of Genomics, and Dr. Rimpy Dhiman, Core Facility of Cell Biology, CSIR-HBT, for conducting RNA Sequencing and Microscopy analyses. We also acknowledge Dr. Saikat, RCB, for providing the *Pst* DC3000 strain. This research was supported by the financial assistance from Ramalingaswami Re-entry Fellowship Grant, Department of Biotechnology, Government of India (grant number BT/RLF/Re-entry/02/2019), Emerging Frontiers in Biotechnology Research grant, Department of Biotechnology, Government of India (grant number BT/PR51519/PBN/18/13/2023), and CSIR-Fundamental & Innovative Research in Science of Tomorrow (CSIR-FIRST) research grant, CSIR, Government of India (grant number 6/1/FIRST/2020-RPPBDD-TMD-SeMI) and to V.D.. S.M. and R.G. gratefully acknowledge UGC, New Delhi, for providing the UGC-SRF and JRF fellowships (1003/(CSIR-UGC NET June 2019) & (JUN21C02082), respectively). T. gratefully acknowledges CSIR, New Delhi, for providing the CSIR-SRF fellowship (No. 31/054[0167]/2020-EMR-I). S.B. gratefully acknowledges the Department of Biotechnology, Government of India, for providing the Research Associateship (DBT-RA/2022/July/N/2552). The article represents the CSIR-IHBT communication number 3941.

## Author Contributions

S.M. and V.D. conceived and designed the project. S.M., A.M., R.G., S.B. and T. performed experiments. A.K and P.K. assisted in metabolite analysis. R.K., N.K. and V.A. assisted in RNAseq data annotation and protein-ligand interactions. S.M., A.M. and V.D. analyzed the data. S.M. and V.D. wrote the manuscript with significant input from all authors.

## Supporting Information

### Supporting Data

**Supplementary Data 1.** Targeted metabolite content in Tea genotypes subjected to drought.

**Supplementary Data 2.** List of genes upregulated in TV-03 genotype as filtered from Parmar et al., 2019.

**Supplementary Data 3.** List of genes downregulated in TV-03 genotype as filtered from Parmar et al., 2019.

**Supplementary Data 4.** Gene ontology enrichment analysis of genes upregulated in TV-03 genotype.

**Supplementary Data 5.** Gene ontology enrichment analysis of genes downregulated in TV- 03 genotype.

**Supplementary Data 6.** List of genes upregulated upon 6 h or 12 h treatment of GDA.

**Supplementary Data 7.** List of genes upregulated upon 6 h or 12 h treatment of TF3G.

**Supplementary Data 8.** List of genes upregulated in either 6 h or 12 h treatment of TF3G.

**Supplementary Data 9.** Gene ontology enrichment analysis of genes upegulated in either 6 h or 12 h treatment of TF3G.

**Supplementary Data 10.** List of genes downregulated in either 6 h or 12 h treatment of TF3G.

**Supplementary Data 11.** Gene ontology enrichment analysis of genes downregulated in either 6 h or 12 h treatment of TF3G.

**Supplementary Data 12.** List of genes stress-responsive genes upregulated in either 6 h or 12 h treatment of TF3G.

**Supplementary Data 13.** List of genes ER stress response-related genes in either 6 h or 12 h treatment of GDA and TF3G.

**Supplementary Data 14.** Promoter analysis of genes upregulated in either 6 h or 12 h treatment of TF3G.

**Supplementary Data 15.** List of genes defense response-related genes in either 6 h or 12 h treatment of GDA and TF3G.

**Supplementary Data 16.** List of primers and oligos used in this study.

## References

1. Apel K, Hirt H. Reactive oxygen species: metabolism, oxidative stress, and signal transduction. Annu Rev Plant Biol 55, 373–399 (2004).

2. Dogra V, Li M, Singh S, Li M, Kim C. Oxidative post-translational modification of EXECUTER1 is required for singlet oxygen sensing in plastids. Nature communications 10, 2834 (2019).

3. Kumar A, Guleria S, Ghosh D, Dogra V, Kumar S. Managing reactive oxygen species - Some learnings from high altitude extremophytes. Environmental and Experimental Botany 189, 104525 (2021).

4. Dogra V, Duan J, Lee KP, Kim C. Impaired PSII proteostasis triggers an UPR-like response in the var2 mutant of Arabidopsis thaliana. J Exp Bot 70, 3075–3088 (2019).

5. Srivastava R, et al. Response to Persistent ER Stress in Plants: A Multiphasic Process That Transitions Cells from Prosurvival Activities to Cell Death. Plant Cell 30, 1220–1242 (2018).

6. Llamas E, Pulido P, Rodriguez-Concepcion M. Interference with plastome gene expression and Clp protease activity in Arabidopsis triggers a chloroplast unfolded protein response to restore protein homeostasis. PLoS Genet 13, e1007022 (2017).

7. Wu GZ, et al. Control of retrograde signalling by protein import and cytosolic folding stress. Nature Plants 5, 525–538 (2019).

8. Yang ZT, et al. The Membrane-Associated Transcription Factor NAC089 Controls ER-Stress-Induced Programmed Cell Death in Plants. Plos Genetics 10, (2014).

9. Meskauskiene R, Nater M, Goslings D, Kessler F, op den Camp R, Apel K. FLU: a negative regulator of chlorophyll biosynthesis in Arabidopsis thaliana. Proc Natl Acad Sci U S A 98, 12826–12831 (2001).

10. Dogra V, Kim C. Singlet Oxygen Metabolism: From Genesis to Signaling. Front Plant Sci 10, 1640 (2020).

11. Kim C, et al. Chloroplasts of Arabidopsis Are the Source and a Primary Target of a Plant-Specific Programmed Cell Death Signaling Pathway. Plant Cell 24, 3026–3039 (2012).

12. Li B, et al. FATTY ACID DESATURASE5 Is Required to Induce Autoimmune Responses in Gigantic Chloroplast Mutants of Arabidopsis. Plant Cell 32, 3240–3255 (2020).

13. Löchli K, et al. Crosstalk between endoplasmic reticulum and cytosolic unfolded protein response in tomato. Cell Stress Chaperon 28, 511–528 (2023).

14. Foyer CH, Ruban AV, Noctor G. Viewing oxidative stress through the lens of oxidative signalling rather than damage. Biochem J 474, 877–883 (2017).

15. Dong J, Chen W. The role of autophagy in chloroplast degradation and chlorophagy in immune defenses during Pst DC3000 (AvrRps4) infection. PloS one 8, e73091 (2013).

16. Pinto-Marijuan M, Munne-Bosch S. Photo-oxidative stress markers as a measure of abiotic stress-induced leaf senescence: advantages and limitations. J Exp Bot 65, 3845–3857 (2014).

17. Bali S, Mohapatra S, Michael R, Arora R, Dogra V. Plastidial metabolites and retrograde signaling: A case study of MEP pathway intermediate MEcPP that orchestrates plant growth and stress responses. Plant Physiol Biochem 222, 109747 (2025).

18. Chan KX, et al. Sensing and signaling of oxidative stress in chloroplasts by inactivation of the SAL1 phosphoadenosine phosphatase. Proc Natl Acad Sci U S A 113, E4567–4576 (2016).

19. Estavillo GM, et al. Evidence for a SAL1-PAP chloroplast retrograde pathway that functions in drought and high light signaling in Arabidopsis. Plant Cell 23, 3992–4012 (2011).

20. Xiao Y, et al. Retrograde signaling by the plastidial metabolite MEcPP regulates expression of nuclear stress-response genes. Cell 149, 1525–1535 (2012).

21. Ramel F, Birtic S, Ginies C, Soubigou-Taconnat L, Triantaphylides C, Havaux M. Carotenoid oxidation products are stress signals that mediate gene responses to singlet oxygen in plants. Proc Natl Acad Sci U S A 109, 5535–5540 (2012).

22. Veal EA, Day AM, Morgan BA. Hydrogen peroxide sensing and signaling. Mol Cell 26, 1–14 (2007).

23. Liu JX, Srivastava R, Che P, Howell SH. Salt stress responses in Arabidopsis utilize a signal transduction pathway related to endoplasmic reticulum stress signaling. The Plant Journal 51, 897–909 (2007).

24. Seo PJ, et al. Cold activation of a plasma membrane□tethered NAC transcription factor induces a pathogen resistance response in Arabidopsis. The Plant Journal 61, 661–671 (2010).

25. Ryan CA, Pearce G. Systemins: a functionally defined family of peptide signals that regulate defensive genes in Solanaceae species. Proceedings of the National Academy of Sciences 100, 14577–14580 (2003).

26. Stührwohldt N, Bühler E, Sauter M, Schaller A. Phytosulfokine (PSK) precursor processing by subtilase SBT3. 8 and PSK signaling improve drought stress tolerance in Arabidopsis. Journal of Experimental Botany 72, 3427–3440 (2021).

27. Dogra V, Singh RM, Li M, Li M, Singh S, Kim C. EXECUTER2 modulates the EXECUTER1 signalosome through its singlet oxygen-dependent oxidation. Mol Plant 15, 438–453 (2022).

28. Paul A, Jha A, Bhardwaj S, Singh S, Shankar R, Kumar S. RNA-seq-mediated transcriptome analysis of actively growing and winter dormant shoots identifies non-deciduous habit of evergreen tree tea during winters. Sci Rep-Uk 4, 5932 (2014).

29. Parmar R, Seth R, Singh P, Singh G, Kumar S, Sharma RK. Transcriptional profiling of contrasting genotypes revealed key candidates and nucleotide variations for drought dissection in Camellia sinensis (L.) O. Kuntze. Sci Rep-Uk 9, 7487 (2019).

30. Lv Z, et al. Research progress on the response of tea catechins to drought stress. J Sci Food Agric 101, 5305–5313 (2021).

31. Cheruiyot EK, Mumera LM, Ng’etich WK, Hassanali A, Wachira F, Wanyoko JK. Shoot epicatechin and epigallocatechin contents respond to water stress in tea [Camellia sinensis (L.) O. Kuntze]. Bioscience, biotechnology, and biochemistry 72, 1219–1226 (2008).

32. Chaeikar SS, Marzvan S, Khiavi SJ, Rahimi M. Changes in growth, biochemical, and chemical characteristics and alteration of the antioxidant defense system in the leaves of tea clones (Camellia sinensis L.) under drought stress. Scientia Horticulturae 265, 109257 (2020).

33. Basu S, Ramegowda V, Kumar A, Pereira A. Plant adaptation to drought stress. F1000Res **5**, (2016).

34. Ling Q, Jarvis P. Regulation of Chloroplast Protein Import by the Ubiquitin E3 Ligase SP1 Is Important for Stress Tolerance in Plants. Curr Biol 25, 2527–2534 (2015).

35. Halliwell B. Reactive species and antioxidants. Redox biology is a fundamental theme of aerobic life. Plant physiology 141, 312–322 (2006).

36. Nimse SB, Pal D. Free radicals, natural antioxidants, and their reaction mechanisms. RSC Advances 5, 27986–28006 (2015).

37. Agati G, Azzarello E, Pollastri S, Tattini M. Flavonoids as antioxidants in plants: location and functional significance. Plant Sci 196, 67–76 (2012).

38. van der Hooft JJ, et al. Structural annotation and elucidation of conjugated phenolic compounds in black, green, and white tea extracts. J Agric Food Chem 60, 8841–8850 (2012).

39. Wang WZ, et al. Insight into catechins metabolic pathways of Camellia sinensis based on genome and transcriptome analysis. J Agr Food Chem 66, 4281–4293 (2018).

40. Leung LK, Su YL, Chen RY, Zhang ZH, Huang Y, Chen ZY. Theaflavins in black tea and catechins in green tea are equally effective antioxidants. J Nutr 131, 2248–2251 (2001).

41. Shan ZG, Nisar MF, Li MX, Zhang CH, Wan CP. Theaflavin Chemistry and Its Health Benefits. Oxid Med Cell Longev 2021, (2021).

42. Subramanian N, Venkatesh P, Ganguli S, Sinkar VP. Role of polyphenol oxidase and peroxidase in the generation of black tea theaflavins. J Agric Food Chem 47, 2571–2578 (1999).

43. Boeckx T, Winters AL, Webb KJ, Kingston-Smith AH. Polyphenol oxidase in leaves: is there any significance to the chloroplastic localization? J Exp Bot 66, 3571–3579 (2015).

44. Tran LT, Taylor JS, Constabel CP. The polyphenol oxidase gene family in land plants: Lineage-specific duplication and expansion. BMC Genomics 13, 395 (2012).

45. Liu Y, Gao L, Xia T, Zhao L. Investigation of the site-specific accumulation of catechins in the tea plant (Camellia sinensis (L.) O. Kuntze) via vanillin-HCl staining. J Agric Food Chem 57, 10371–10376 (2009).

46. Boeckx T, Webster R, Winters AL, Webb KJ, Gay A, Kingston-Smith AH. Polyphenol oxidase-mediated protection against oxidative stress is not associated with enhanced photosynthetic efficiency. Ann Bot 116, 529–540 (2015).

47. Huang C, et al. Two New Polyphenol Oxidase Genes of Tea Plant (Camellia sinensis) Respond Differentially to the Regurgitant of Tea Geometrid, Ectropis obliqua. Int J Mol Sci 19, (2018).

48. Zhang J, Zhang X, Ye M, Li XW, Lin SB, Sun XL. The Jasmonic Acid Pathway Positively Regulates the Polyphenol Oxidase-Based Defense against Tea Geometrid Caterpillars in the Tea Plant (Camellia sinensis). J Chem Ecol 46, 308–316 (2020).

49. Huang X, et al. The R2R3 Transcription Factor CsMYB59 Regulates Polyphenol Oxidase Gene CsPPO1 in Tea Plants (Camellia sinensis). Front Plant Sci 12, 739951 (2021).

50. Mayer AM. Polyphenol oxidases in plants and fungi: going places? A review. Phytochemistry 67, 2318–2331 (2006).

51. Araji S, et al. Novel roles for the polyphenol oxidase enzyme in secondary metabolism and the regulation of cell death in walnut. Plant physiology 164, 1191–1203 (2014).

52. Chevalier T, de Rigal D, Mbeguie AMD, Gauillard F, Richard-Forget F, Fils-Lycaon BR. Molecular cloning and characterization of apricot fruit polyphenol oxidase. Plant physiology 119, 1261–1270 (1999).

53. Sommer A, Ne’eman E, Steffens JC, Mayer AM, Harel E. Import, targeting, and processing of a plant polyphenol oxidase. Plant physiology 105, 1301–1311 (1994).

54. Derardja AE, Pretzler M, Kampatsikas I, Barkat M, Rompel A. Purification and Characterization of Latent Polyphenol Oxidase from Apricot (Prunus armeniaca L.). J Agric Food Chem 65, 8203–8212 (2017).

55. Peng KL, Xue CJ, Huang XZ. Enhancing virus-induced gene silencing efficiency in tea plants (Camellia sinensis L.) and the functional analysis of CsPDS. Scientia Horticulturae 337, (2024).

56. Beaugelin I, Chevalier A, D’Alessandro S, Ksas B, Havaux M. Endoplasmic reticulum-mediated unfolded protein response is an integral part of singlet oxygen signalling in plants. Plant Journal 102, 1266–1280 (2020).

57. Kuroyanagi M, Yamada K, Hatsugai N, Kondo M, Nishimura M, Hara-Nishimura I. Vacuolar Processing Enzyme Is Essential for Mycotoxin-induced Cell Death in Arabidopsis thaliana. Journal of Biological Chemistry 280, 32914–32920 (2005).

58. He R, et al. Metacaspase-8 modulates programmed cell death induced by ultraviolet light and H2O2 in Arabidopsis. Journal of Biological Chemistry 283, 774–783 (2008).

59. Shumbe L, Chevalier A, Legeret B, Taconnat L, Monnet F, Havaux M. Singlet Oxygen-Induced Cell Death in Arabidopsis under High-Light Stress Is Controlled by OXI1 Kinase. Plant physiology 170, 1757–1771 (2016).

60. Angelos E, Ruberti C, Kim SJ, Brandizzi F. Maintaining the factory: the roles of the unfolded protein response in cellular homeostasis in plants. Plant Journal 90, 671–682 (2017).

61. Sun L, et al. The plant-specific transcription factor gene NAC103 is induced by bZIP60 through a new cis-regulatory element to modulate the unfolded protein response in Arabidopsis. Plant Journal 76, 274–286 (2013).

62. Bhadresha K, et al. Theaflavin-3-gallate, a natural antagonist for Hsp90: *In-silico* and in-vitro approach. Chem-Biol Interact 353, (2022).

63. Schopf FH, Biebl MM, Buchner J. The HSP90 chaperone machinery. Nat Rev Mol Cell Bio 18, 345–360 (2017).

64. Luengo TM, Mayer MP, Rüdiger SGD. The Hsp70-Hsp90 Chaperone Cascade in Protein Folding. Trends in cell biology 29, 164–177 (2019).

65. Kadota Y, Shirasu K. The HSP90 complex of plants. Bba-Mol Cell Res 1823, 689–697 (2012).

66. Marcu MG, Doyle M, Bertolotti A, Ron D, Hendershot L, Neckers L. Heat shock protein 90 modulates the unfolded protein response by stabilizing IRE1α. Mol Cell Biol 22, 8506–8513 (2002).

67. Pearl LH, Prodromou C. Structure and mechanism of the Hsp90 molecular chaperone machinery. Annual review of biochemistry 75, 271–294 (2006).

68. Li L, Wang L, You QD, Xu XL. Heat Shock Protein 90 Inhibitors: An Update on Achievements, Challenges, and Future Directions. J Med Chem 63, 1798–1822 (2020).

69. Watanabe N, Lam E. BAX inhibitor-1 modulates endoplasmic reticulum stress-mediated programmed cell death in Arabidopsis. Journal of Biological Chemistry 283, 3200–3210 (2008).

70. Xi HM, Xu H, Xu WX, He ZY, Xu WZ, Ma M. A SAL1 Loss-of-Function Arabidopsis Mutant Exhibits Enhanced Cadmium Tolerance in Association with Alleviation of Endoplasmic Reticulum Stress. Plant and Cell Physiology 57, 1210–1219 (2016).

71. Sitia R, Braakman I. Quality control in the endoplasmic reticulum protein factory. Nature 426, 891–894 (2003).

72. Moreno AA, et al. IRE1/bZIP60-Mediated Unfolded Protein Response Plays Distinct Roles in Plant Immunity and Abiotic Stress Responses. PloS one 7, (2012).

73. Kadota Y, et al. Structural and functional analysis of SGT1-HSP90 core complex required for innate immunity in plants. Embo Rep 9, 1209–1215 (2008).

74. Marone D, et al. Specialized metabolites: Physiological and biochemical role in stress resistance, strategies to improve their accumulation, and new applications in crop breeding and management. Plant Physiol Biochem 172, 48–55 (2022).

75. Christian D, Stahl N, Niri V, Geetha-Loganathan P. Biochemical Analysis of Browning Activities in Apples. Biol Bull+ 51, 619–624 (2024).

76. Flurkey WH, Inlow JK. Proteolytic processing of polyphenol oxidase from plants and fungi. J Inorg Biochem 102, 2160–2170 (2008).

77. Bertolotti A, Zhang YH, Hendershot LM, Harding HP, Ron D. Dynamic interaction of BiP and ER stress transducers in the unfolded-protein response. Nature Cell Biology 2, 326–332 (2000).

78. Iwata Y, Fedoroff NV, Koizumi N. bZIP60 Is a Proteolysis-Activated Transcription Factor Involved in the Endoplasmic Reticulum Stress Response. Plant Cell 20, 3107–3121 (2008).

79. Ko DK, Brandizzi F. Advanced genomics identifies growth effectors for proteotoxic ER stress recovery in. Communications Biology 5, (2022).

80. Sandelius AS, Andersson MX, Goksör M, Tjellström H, Wellander R. Membrane contact sites:: physical attachment between chloroplasts and endoplasmic reticulum revealed by optical manipulation. Chem Phys Lipids 149, S42–S43 (2007).

81. Jayaswall K, et al. Transcriptome Analysis Reveals Candidate Genes involved in Blister Blight defense in Tea ( (L) Kuntze). Scientific Reports 6, (2016).

82. Babu A, et al. Impact of Climate Change on Tea Cultivation and Adaptation Strategies. In: Climate Change and Agriculture) (2022).

83. Seth R, Maritim TK, Parmar R, Sharma RK. Underpinning the molecular programming attributing heat stress associated thermotolerance in tea ( (L.) O. Kuntze). Hortic Res-England 8, (2021).

84. Singh HP, Ravindranath SD. Occurrence and Distribution of Ppo Activity in Floral Organs of Some Standard and Local Cultivars of Tea. J Sci Food Agr 64, 117–120 (1994).

85. Nakagawa T, et al. Improved Gateway binary vectors: high-performance vectors for creation of fusion constructs in transgenic analysis of plants. Bioscience, biotechnology, and biochemistry 71, 2095–2100 (2007).

86. Ghawana S, et al. An RNA isolation system for plant tissues rich in secondary metabolites. BMC Res Notes 4, 85 (2011).

87. Chen SF, Zhou YQ, Chen YR, Gu J. fastp: an ultra-fast all-in-one FASTQ preprocessor. Bioinformatics 34, 884–890 (2018).

88. Deng ZL, Münch PC, Mreches R, McHardy AC. Rapid and accurate identification of ribosomal RNA sequences via deep learning. Nucleic Acids Res 50, (2022).

89. Kim D, Paggi JM, Park C, Bennett C, Salzberg SL. Graph-based genome alignment and genotyping with HISAT2 and HISAT-genotype. Nature Biotechnology 37, 907-+ (2019).

90. Pertea M, Kim D, Pertea GM, Leek JT, Salzberg SL. Transcript-level expression analysis of RNA-seq experiments with HISAT, StringTie and Ballgown. Nature Protocols 11, 1650–1667 (2016).

91. Tang Y, Horikoshi M, Li WX. ggfortify: Unified Interface to Visualize Statistical Results of Popular R Packages. R J 8, 474–485 (2016).

