## Supplementary Figures 1-16 for "Oxidative stress-induced proteolytic activation of polyphenol oxidase triggers an oxidized flavonoids-mediated stress signaling in *Camellia sinensis*"

[*]Author for correspondence: Vivek Dogra

- Supporting Information
  - Supplementary Figures 1-16
  - Supplementary Data 1-16

a

b

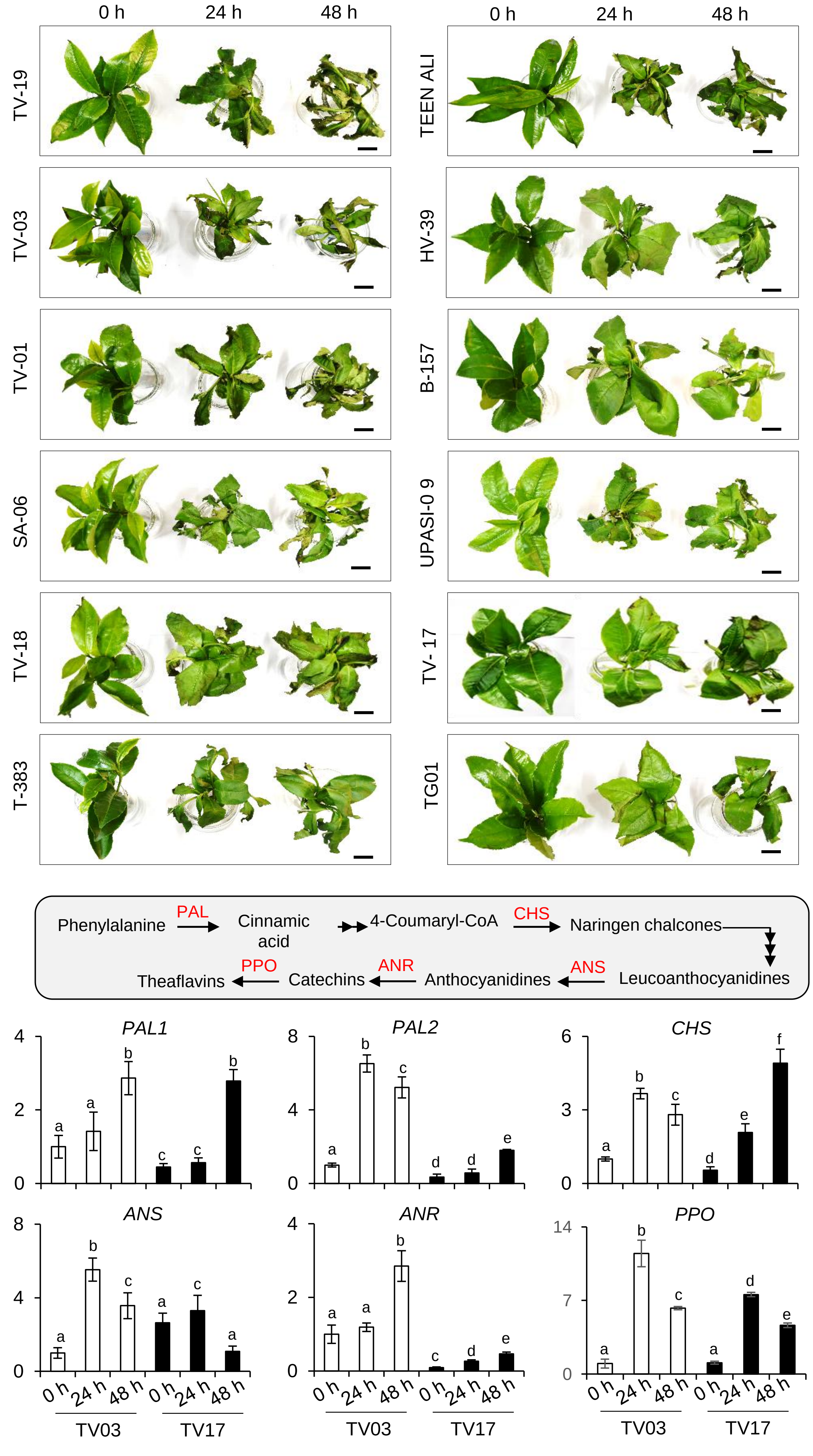

0 h
24 h
48 h
TV-19
TV-03
TV-01
SA-06
TV-18
T-383
TEEN ALI
HV-39
B-157
UPASI-0 9
TV- 17
TG01
Phenylalanine
PAL
Cinnamic acid
4-Coumaryl-CoA
CHS
Naringen chalcones
Leucoanthocyanidines
ANS
Anthocyanidines
ANR
Catechins
PPO
Theaflavins
Relatve expression/ GAPDH
PAL1
PAL2
CHS
ANS
ANR
PPO
TV03
TV17

**Supplementary Fig. 1:** Drought-induced plant phenotype and expression of theaflavins biosynthesis.

**a** Macroscopic phenotype. Shoot cuttings of 12 genotypes collected were incubated in Hoagland's nutrient medium containing 10% PEG for 48 h. Images are representative of three biological replicates. Scale bar = 2 cm. **b** Theaflavins biosynthetic pathway and relative expression of key genes in susceptible (TV03) and resilient (TV17) genotypes. The relative expression levels of *PAL1*, *PAL2*, *CHS*, *ANS*, *ANR*, and *PPO* were analyzed using qRT-PCR. *GAPDH* was used as an internal control. Data represent the mean of three independent biological replicates. Error bars indicate standard deviation (SD). Lowercase letters indicate statistically significant differences between the mean values ($P < 0.05$, one-way analysis of variance with post hoc Tukey's Honest Significant Difference (HSD) test).

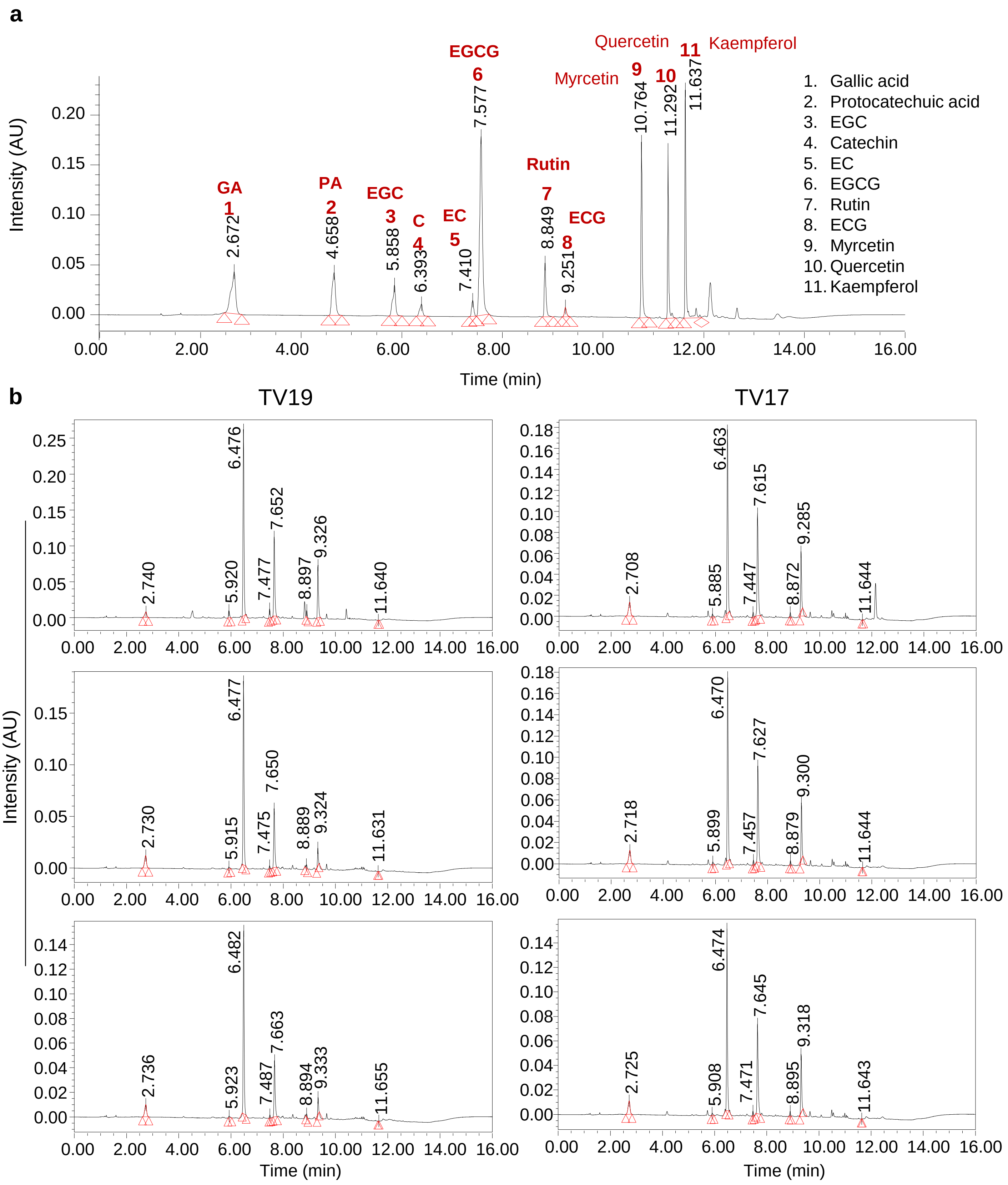


**Supplementary Fig. 2:** UPLC chromatograms of catechins.

**a** Standard-mix of phenolic acids and flavonoids. **b** Drought-treated TV19 (susceptible) and TV17 (resilient) genotypes of Tea.

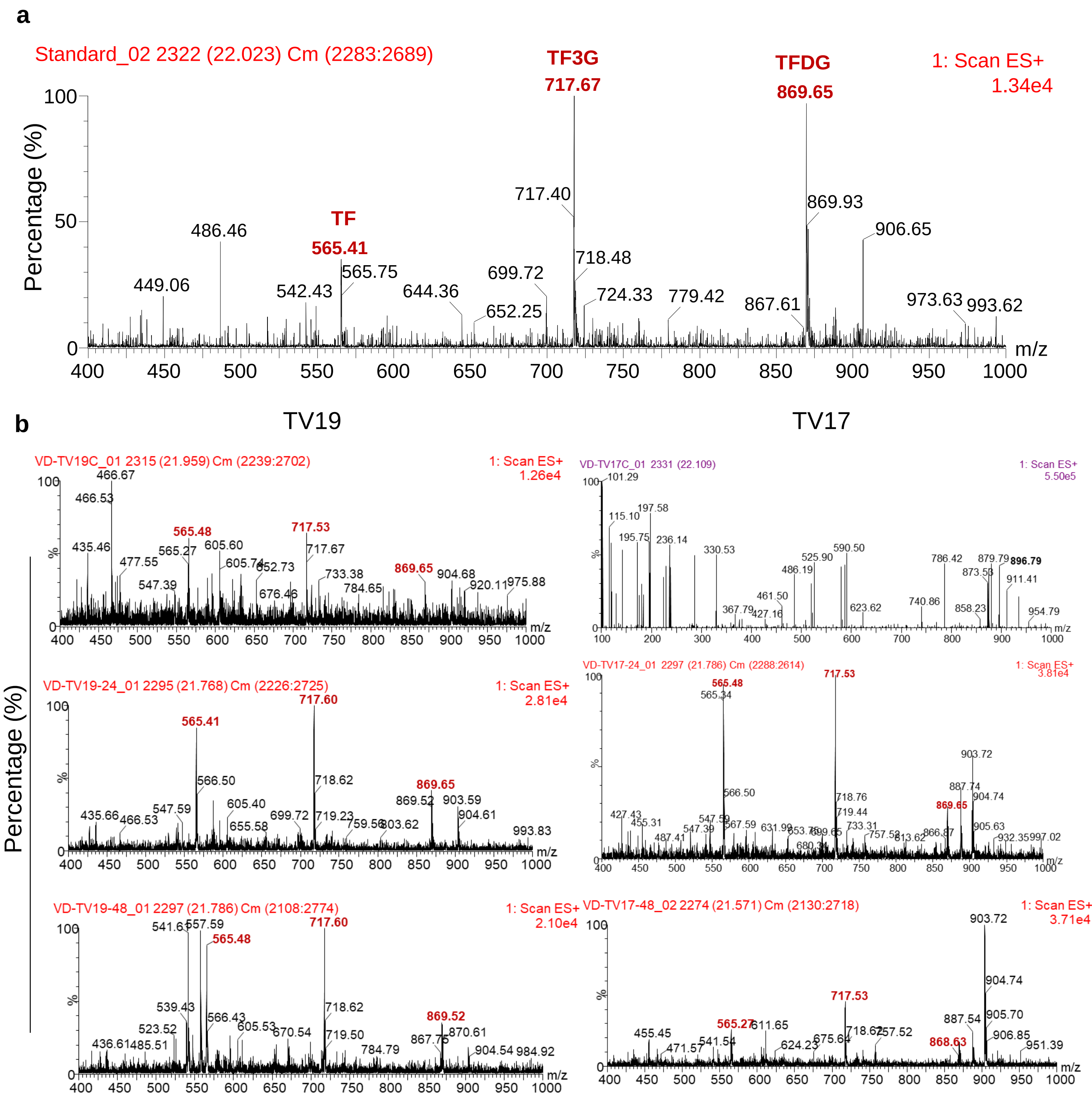


**Supplementary Fig. 3:** UPLC-Mass spectra of theaflavins.

**a** Standard-mix of TF, TF3G and TFDFG. **b** Drought-treated TV19 (susceptible) and TV17 (resilient) genotypes of Tea. Theaflavin (TF) MW: 564.499; Theaflavin-3-gallate (TF3G) MW: 716.604, Theaflavin-di-gallate (TFDG) MW: 868.700

**a**

```
PaPPO   MATAPSPTTMGT---YSSLISTNSFSTFLPNKSQLSLSGKSKHYVARRSSISCKATNNNNSNNQNEQQEESSRLLGKLDRRNILIGLGGLYGAT--TLDRKPFAFADPIA 105
VvPPO   MASLPWSLTTSTAIANTTNISAFPPSPLFQRASHVPVARNRSRRFA-PSKVSCNSANGD-PNS-DSTSDVRETSSGKLDRRNVLLGIGGLYGAAGGLGATKPLAFGAPIQ 107
CsPPO2  MASILPPTTTK----TTTTSSTLYSYPIFKNTSKIPTIRKHNHSF----NVSCSKAKDSDPNL-TPPSQNTQTSLGKFDRRNMVIGLGGLYGAA-GLTDTDPPALAAPVT 100
CsPPO1  MNSLPPSCTTI---------PTHSSCAFFKNTFQVPTIGKSKHCF----KVSCKAIDGDTNEK------TPNPNPPKLDRRNMLIGLGGYGAAT---LAGNPPATASPIS 88
CsPPO3  MASFPPSTTTT---AASTATITPTSSPFFKKTLQIPTLRNQRHSF----KISCKSTDNDNQNP-NTTSKNEETFLGKLDRRNVLLGLGGYSAAT---ILANPLAFAAPVT 99
        * :     *              :     :: .  ::    :  :  .    .:**.  ... :          .    *:****:::*:**  .*:      .* * . *:

PaPPO   PPDLTTCKPAEITPGGSETVPCCPPVTTKIKTFKP-DLSIPLRTSPAAHQVTDEYLAKFKKAQAAMRALPDDDPRSMVQQAKVHCAYCNGAYPQVGFTDNDIQVHFSWLF 214
VvPPO   APDISKCGTATVP-DGVTPTNCCPPVTTKIIDFQLPSSGSPMRTRPAAHLVSKEYLAKYKKAIELQKALPDDDPRSFKQQANVHCTYCQGAYDQVGYTDLELQVHASWLF 216
CsPPO2  APDLSKCGAADLP-ADAKPTNCCPPKTNKIIEFKLPPPSNILRVRPAAHLADEKYIAKFSKALQLMKSLPDDDPRSFKQQSNIHCAYCEGAYHQVGFPSTELQVHNSWLF 209
CsPPO1  PPDVTKCGPAELP-SGAKPTNCCPPKSTKIIDFKLPPPPTIVRVRPAVHLADDEYIAKFSKALALMKALPEDDPRSFTQQANVHCAYCDGAYHQVGFPDLELQVHNSWLF 197
CsPPO3  APDLAKCGAADLP-AGAKPTNCCPPKATKIIDFKFPQS-NSMRVRPAAHLADAEYVAKFSKALALMKALPEDDPRSFKQQANVHCAYCDGAYHQVGFPELDLQVHNSWLF 207
         **::.*  * :    .    . **** :.**  *:         :*. **.* .  :*:**:.**    ::**:*****: **:::**:**:*** ***: . ::*** ****

PaPPO   FPFHRMYLYFYERILGKLIDDPTFALPYWNWDSPVGFPIPDIYTDTSSPLYDQYRNADHQPPVLVDLSYGGKDDDVDE-QTRIDENLAIMYRQMVSGAKTPDLFFGHAYR 323
VvPPO   LPFHRYYLYFNERILAKLIDDPTFALPYWAWDNPDGMYMPTIYASSPSSLYDEKRNAKHLPPTVIDLDYDGTEPTIPD-DELKTDNLAIMYKQIVSGATTPKLFLGYPYR 325
CsPPO2  FPFHRFYLYFFEKILGMLLDDPAFAIPFWNWDSPAGMKIPAMYADINSPLYNRLRDAKHQPPTLIDLDYNLTDPKNVDEEKQKLRNLTIMYRQVVSGGKTPRLFLGSSYR 319
CsPPO1  FPYHRYYLYFFEKILGKLIGDPSFAIPFWNWDSPIGMSMPPIYADPKSPMYDRLRDAKHQPPITIDLDYNLTDSKSPS-EKLISNNLTIMYRQVVSSGKTAKLFLGSAYR 306
CsPPO3  FPFHRYYLHFFERILGSLIGDPSFAIPFWNWDSPNGMPMPAMYADPNSPLYDVLRDAKHQPPTLVDLDYNGSDPTTTD-AQQKSSNLNVMYRQMVSNSKSARLFLGSPYR 316
        :*:** **:* *:**. *:.**:**:*:* **.* *: :* :*:.  * :*:  *:*.* **  :**.*. .:     .        ** :**:*:**...:  **:*  **

PaPPO   AGNLNTGKYPGTIENMPHNNIHIWVGDPSQTHQEDMGNFYSAGRDPLFYAHHANVDRMWNIWKTLGGK-RKDITDTDWLDAEFLFYDENAELVRVKVRDSLEPEKQLRYN 432
VvPPO   AGDAI-DPGAGTLEHAPHNIVHKWTGLADKP-SEDMGNFYTAGRDPIFFGHHANVDRMWNIWKTIGGKNRKDFTDTDWLDATFVFYDENKQLVKVKVSDCVDT-SKLRYQ 432
CsPPO2  AGDDP-DPGAGSLENIPHGPVHIWCGDRTQPNLEDMGNFYSAGRDPIFYGHHANVDRIWTVWKTLGGK-RNDFKDSDCLNSEFTFYDENAQLVTVKVKESLDH-RKLGYV 426
CsPPO1  AGDDP-DPGAGALENIPHGPVHIWCGDRTQPNIENMGNLYSSGRDPMFFAHHGNVDRMWTIWKSLG---RRDFTDPDWLDSGFFFYDENAQPVKVKVKDCLDM-TKLGYV 411
CsPPO3  AGDES-DPGAGSVENIPHGPVHIWCGDRTQPNLEDMGNFYSAGRDPIFFGHHANIDRMWSIWKTLGGK-RQDFTDPDWLNAGFVFYDENAELVRVKVKDCLDT-RKLGYV 423
        **:    .    *::*: **. :* * *   :    *:***:*::****:*:.**.*:**:*.:**::*    *.*:.* * *:: * ***** : * *** :.::   :* *

PaPPO   YEPVSLPWLFTKPTARKTKNK------TKAKVAAT------QLTSKFPATLVEVTTVEVARPKPRKRSKKEKVDEEELLIIKDIEFEGTEAVKFDVFINDDAESLSRRDK 530
VvPPO   YQDIPIPWLPKNTKAKAKTTT--KSSKSGVAKAAELPKTTISSIGDFPKALNSVIRVEVPRPKK-SRSKKEKEDEEEVLLIKGIELDRENFVKFDVYINDEDYSVSRPKN 539
CsPPO2  YQDVEIPWLNARPSPRISIFFRKIKNKAGIAMATETLD---SAAIVFPRKLDEVVKVVVKRPTK-SRSEREKEEEEEVVVVEGIEMERDVSVKFDVFINDEDEAASGPEK 532
CsPPO1  YQDVDIPWLKSRPTPRVSKLSRKT-NKAGVAKAAETQ-------PTFPTKLDKVVRATVQRPKK-SRSKKEKEEEEEILVIDGIEVERDQFVKFDVFINDGDEPVIGPEN 512
CsPPO3  YQDVDIPWLNTRPTPGVSKLSRKI-NKAGVAMAADTPS---TATEVFPKKLDKAVRVMVPRPKK-SRSKKEKDDEEEILVIEGIEVDRDVFVKFDVFINDEHEVVIGPNN 528
        *: : :***  . .   .          :    *:            **  *  ..  . * **.  .**::** :***::::..**.:    *****:***         .:

PaPPO   SEFAGSFVHVPQGKTTKAKTKTNLKLGITDLLEDLGAEDDSSVLVTLVPRVSNSPITIGGFKIEYSS- 597
VvPPO   SEFAGSFVNVPHKHMKEMKTKTNLRFAINELLEDLGAEDDESVIVTIVPRAGGDDVTIGGIEIEFVSD 607
CsPPO2  TEFAGSFVNVPRKHKHDKKIRTSLRLGITELLEDLEAEDDESVLVTLVPRYGSDAVTIGGVKIEFDS- 599
CsPPO1  SEFAGSFVNVPHRHKHGKKMKTCLRLGISELLEDLEAEDDESVLVTLVPRKGSDALIIGGIKIEFDS- 579
CsPPO3  TEFAGSFVNVPHKHKHGKKIKTILRLGITELLEELGAEDDDGVLVTLVPKNGAGAVTIGGVKIEFDS- 595
        :*******:**: :    * :* *::.*.:***:* ****.*..*:**:**: . . : ***.:**: *
```

**b**

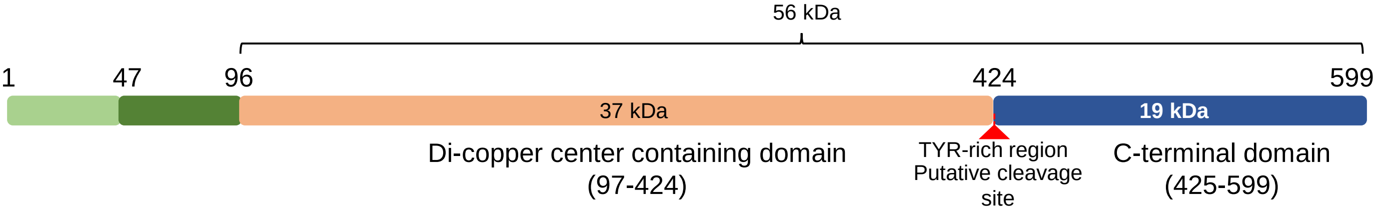


**Supplementary Fig. 4:** Alignment of *Cs*PPO proteins with characterised PPOs from other plant species.

**a** Protein sequences of three PPO proteins of *Camellia sinensis* var. assamica, were retrieved from NCBI, *Cs*PPO1 (QDX19551), *Cs*PPO2 (QDX19552 which is same as ACN38060.1, characterized earlier Sing et al., 2017), and *Cs*PPO3 (QDX19553) and were aligned with PPOs of *Prunus armeniaca* (*Pa*PPO; Uniprot Id: O81103) and *Vitis vinifera* (*Vv*PPO; Uniprot Id: P43311). Alignment suggests the presence of a plastid transit peptide (light green), followed by a transit peptide for thylakoids (dark green). The latent PPO is shown in dark red, which is expected to be cleaved from full mature protein. **b** Domains and putative cleavage site in *Cs*PPO2 protein.

**a**

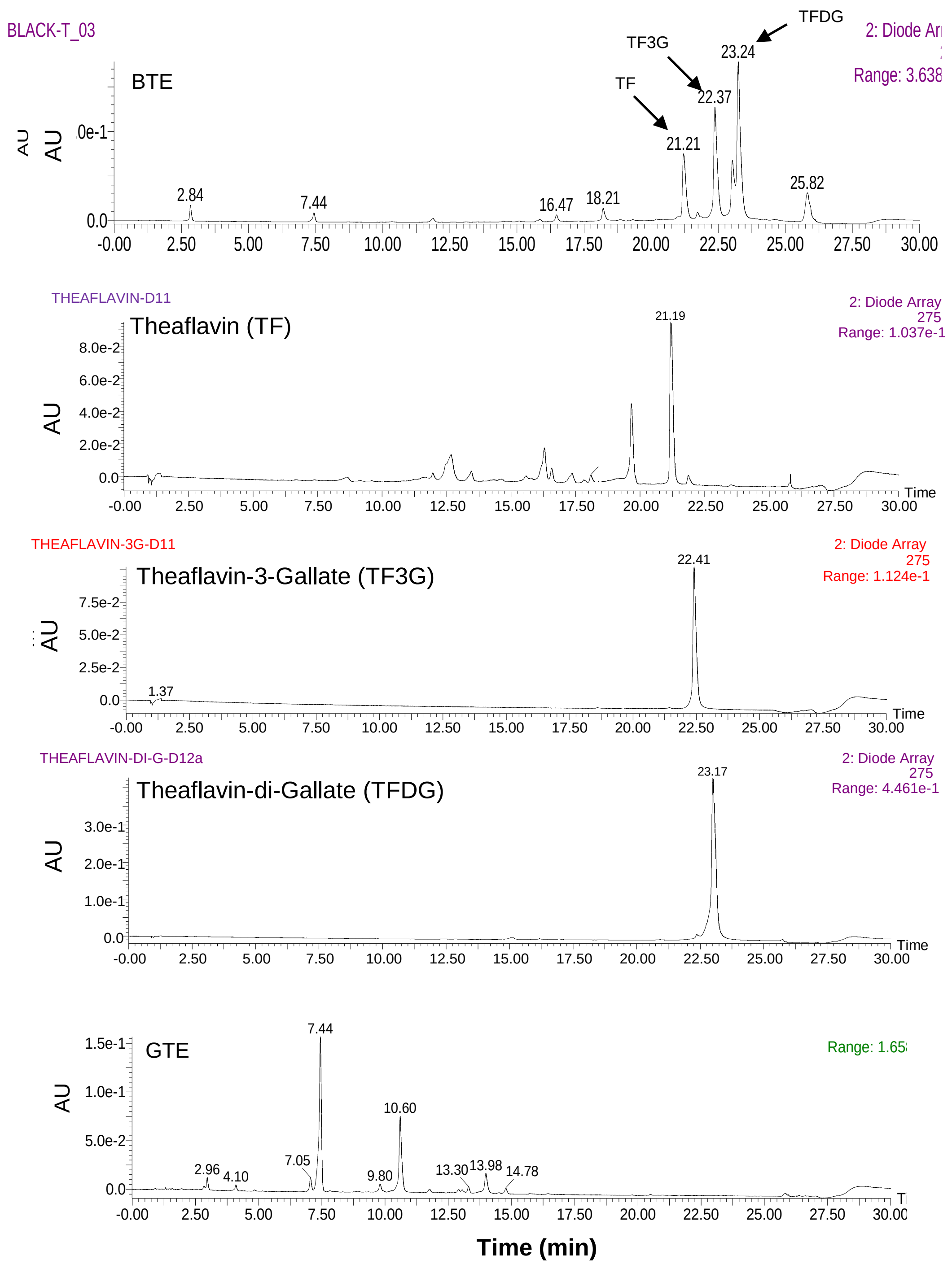


**b**

| TF | Concentration (%) | |
|---|---|---|
| | GTE | BTE |
| TF | 0.03 | 3.49 |
| TF3G | 0.30 | 6.35 |
| TFDG | 0.65 | 14.77 |

**Supplementary Fig. 5** UPLC chromatograms of Tea extracts and pure theaflavins.

**a** Black tea extract (BTE), Theaflavin (TF), Theaflavins-3-gallate (TF3G), and Theaflavin-3-digallate (TFDG) and Green Tea extract (GTE). **b** Proportion of TFs in BTE..

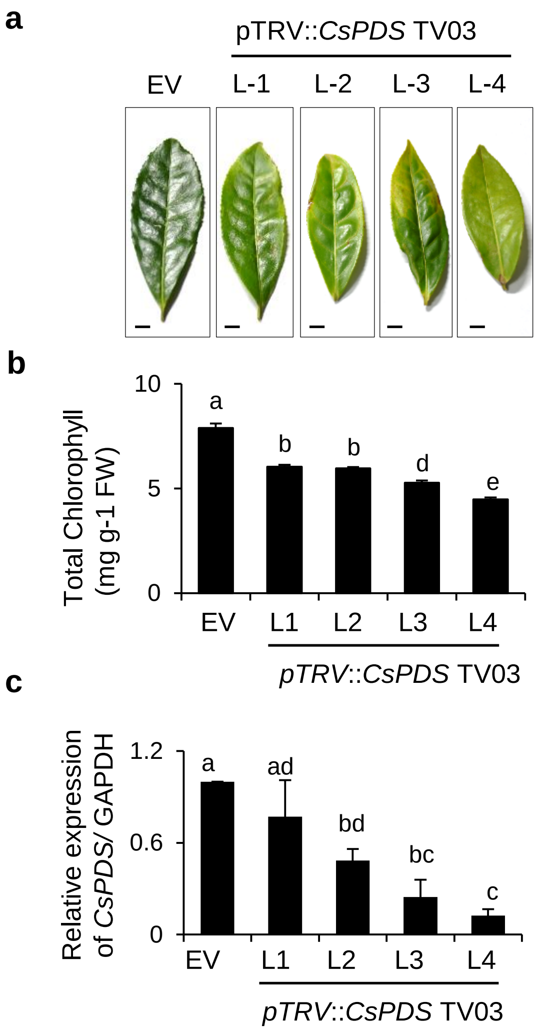


**Supplementary Fig. 6:** Virus-induced gene silencing of *Phytoene desaturase* (*CsPDS*) in Tea.

**a** Macroscopic phenotype. **b** Chlorophyll content. **c** Relative expression levels of *CsPDS*. Shoot cuttings of TV03 genotype were inoculated with either pTRV1-pTRV2 (empty vector; EV) or pTRV1-pTRV2-CsPDS (*pTRV::CsPDS*) were grown for 5 weeks and newly developed leaves were observed and harvested. For qRT-PCR, equal amounts of cDNA prepared from RNA extracted from the leaf tissue, were used. *CsGAPDH* was used as an internal control. Data in (**b-c**) represent the mean of three independent biological replicates ± SD. Lowercase letters indicate statistically significant differences between the mean values ($P < 0.05$, one-way analysis of variance with post hoc Tukey's HSD test). Scale bar in **(b)** = 2 cm.

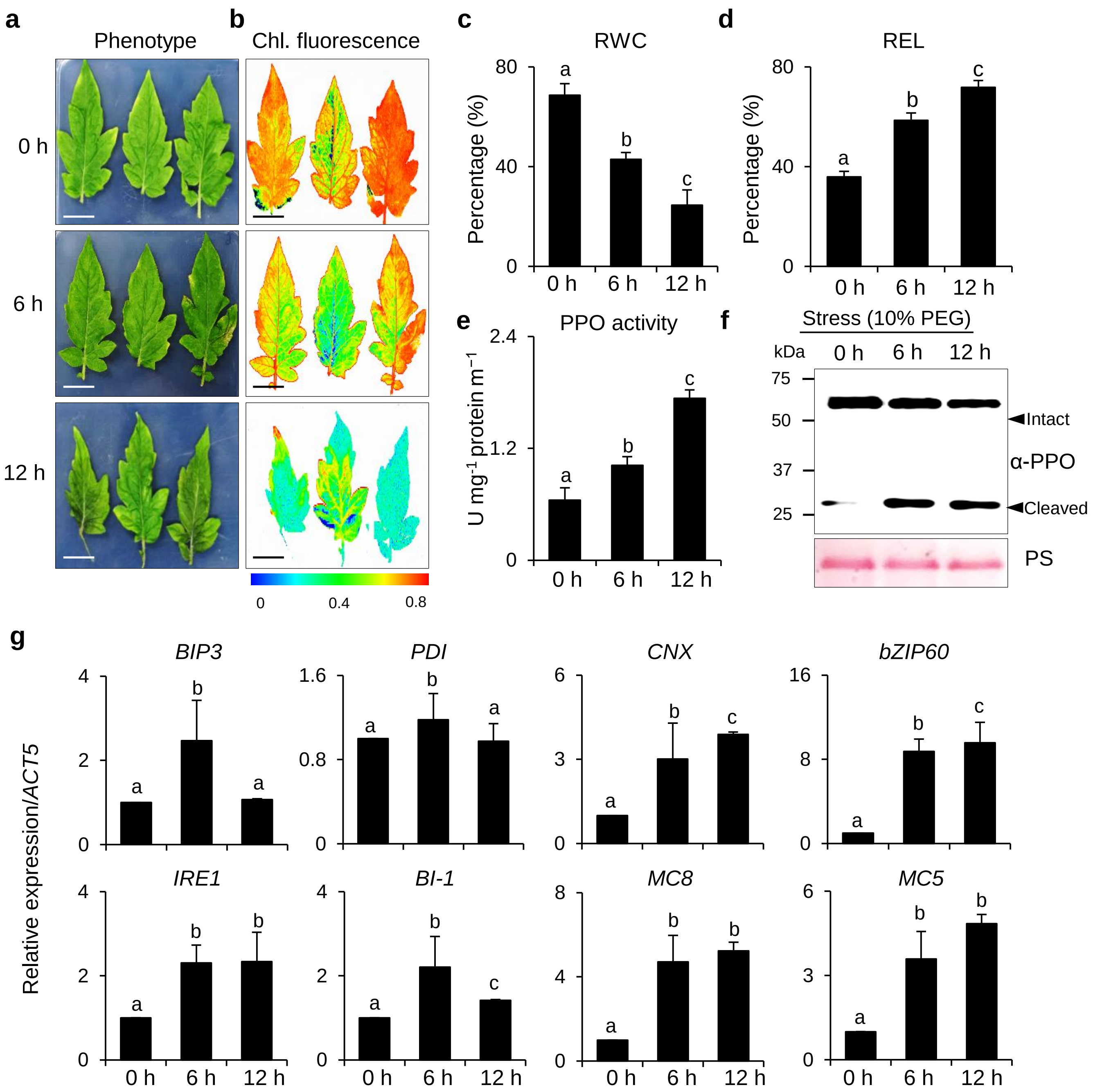


**Supplementary Fig. 7:** Drought-induced activation of PPO and downstream ER-stress-like response in Tomato.

**a-d** Macroscopic phenotype (**a**), chlorophyll fluorescence representing PSII efficiency (**b**), Relative water content (RWC) (**c**) and Relative electrolyte leakage (REL) (**d**). Three-week-old Tomato seedlings were treated with Hoagland's nutrient medium containing 10% PEG for a period of 12 h. **e** Polyphenol oxidase (PPO) activity. **f** Immunoblot analysis of PPO. Total proteins were extracted from leaf tissue and immunoblotted using anti-PPO antibody. **g** Relative expression levels of genes involved in general ER-stress response. qRT-PCR was performed using equal amounts of cDNA prepared from RNA extracted from the leaf tissue. *Tomato Actin 5 (ACT5)* was used as an internal control. Data in (**c-e** and **g**) represent the mean of three independent biological replicates. Error bars indicate standard deviation (SD). Lower case letters indicate statistically significant differences between the mean values ($P < 0.05$, one-way analysis of variance with post hoc Tukey's HSD test). Scale bar in **(a)** = 2 cm.

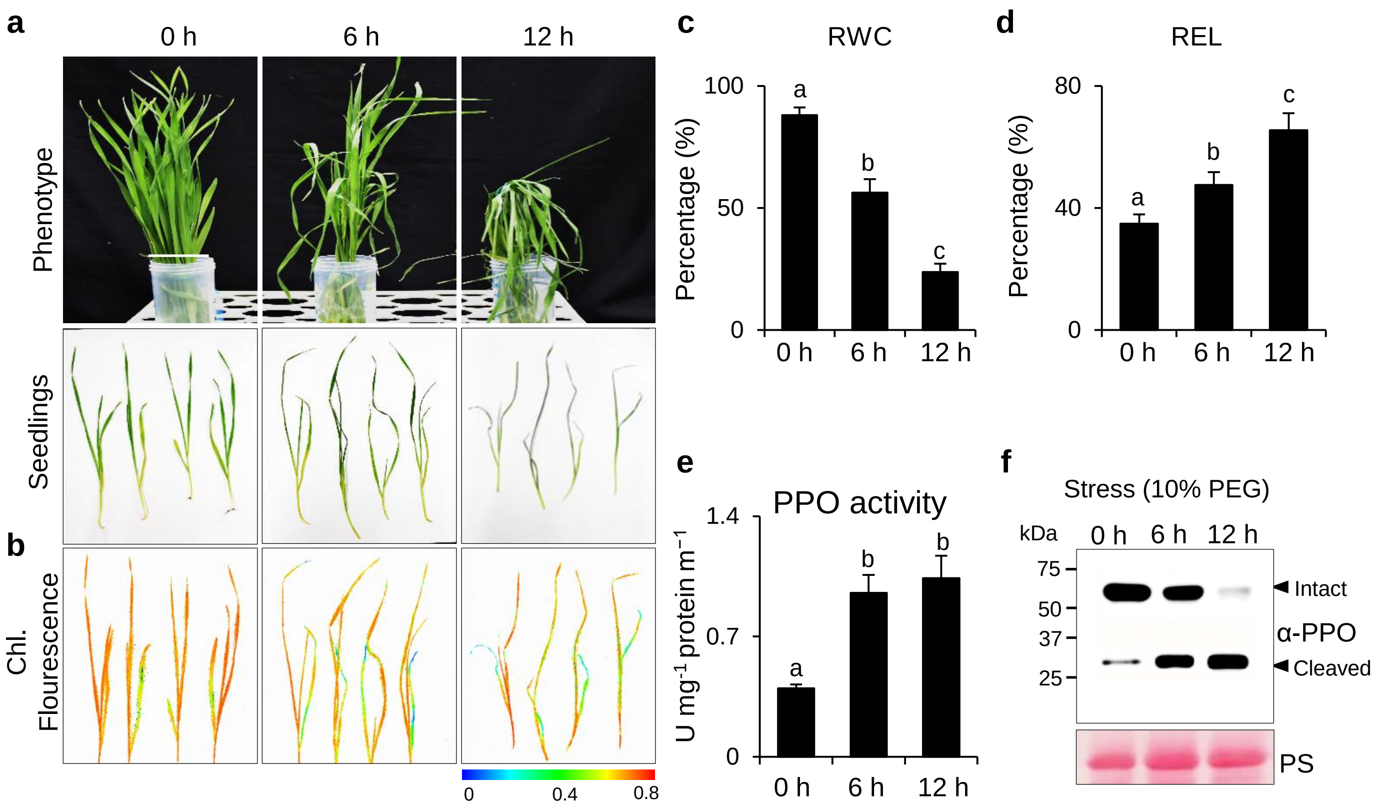


**Supplementary Fig. 8:** Drought-induced activation of PPO and downstream cell death in Wheat.

**a-d** Macroscopic phenotype (**a**), chlorophyll fluorescence representing PSII efficiency (**b**), Relative water content (RWC) (**c**) and Relative electrolyte leakage (REL) (**d**). Three-week-old Wheat seedlings were treated with Hoagland's nutrient medium containing 10% PEG for a period of 12 h. **e** Polyphenol oxidase (PPO) activity. **f** Immunoblot analysis of PPO. Total proteins were extracted from leaf tissue and immunoblotted using anti-PPO antibody. Data in (**c-e**) represent the mean of three independent biological replicates. Error bars indicate standard deviation (SD). Lowercase letters indicate statistically significant differences between the mean values ($P < 0.05$, one-way analysis of variance with post hoc Tukey's HSD test).

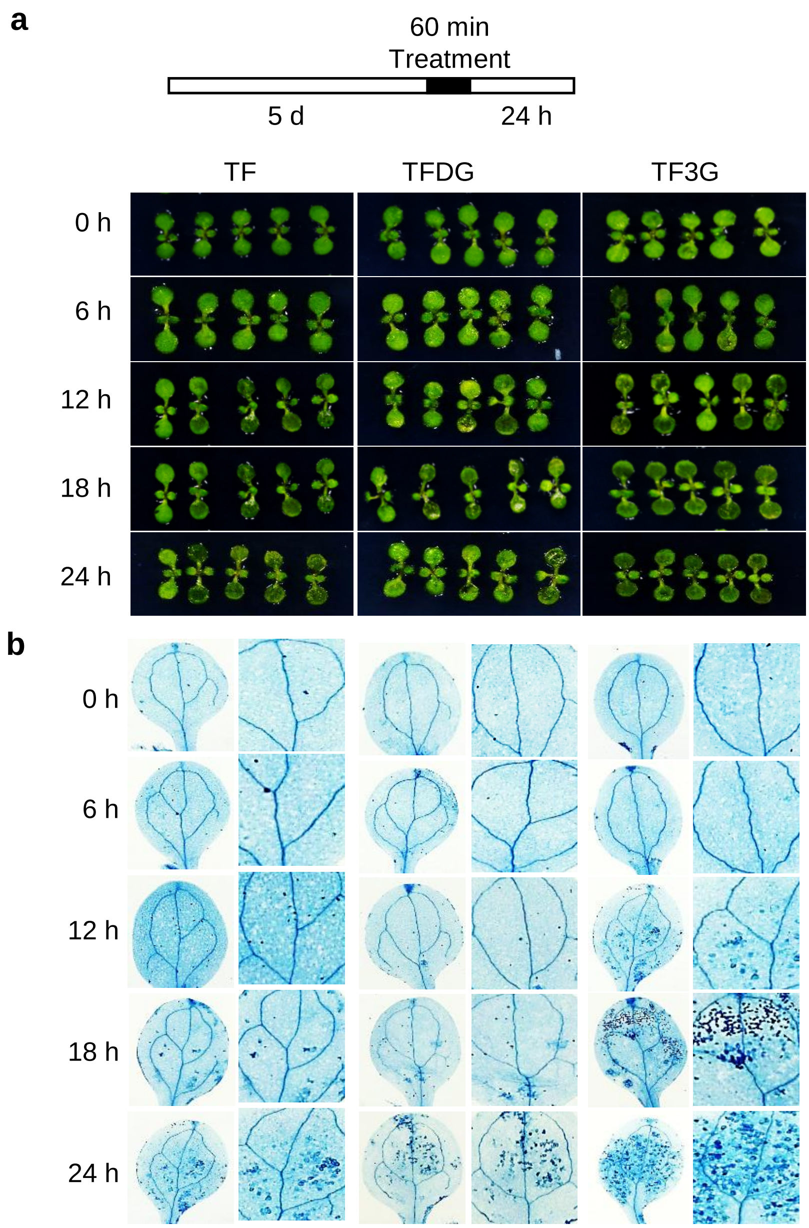


**Supplementary Fig. 9:** TFs-induced phenotypes are time-dependent.

**a** Plant phenotype. **b** Cell death. Five-day-old seedlings of wild-type Arabidopsis were vacuum infiltrated in a solution containing 200 µm TFs for 5 min followed by 55 min incubation in the same solution. Treated seedlings were then transferred to fresh liquid MS and shifted to the growth chamber. Five seedlings from each treatment were harvested after indicated time points and stained with TB staining.

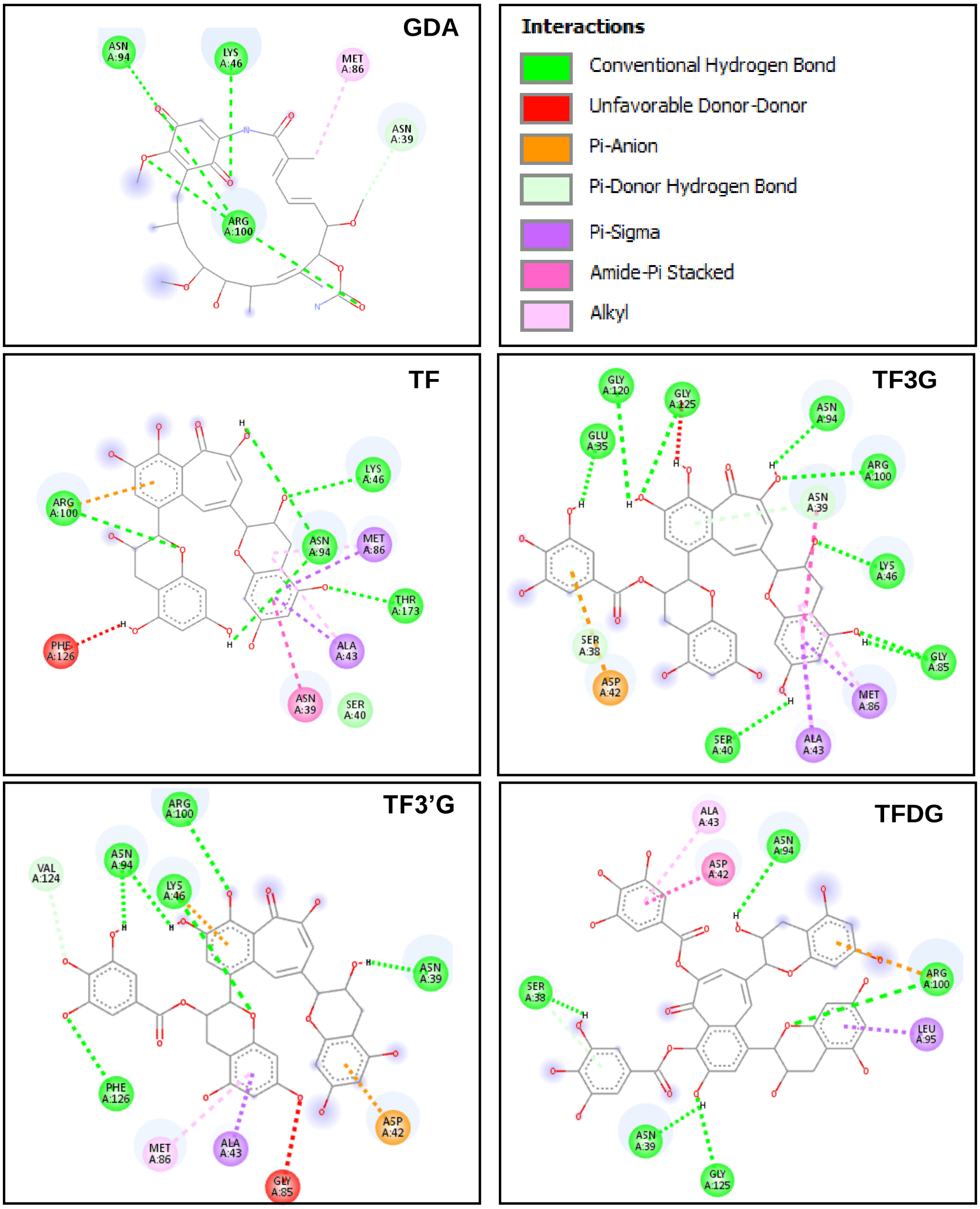


| Chemical | Binding affinity | Interacting amino acids residues |
|---|---|---|
| GDA | -5.9 | Asn39, Lys46, Met86, Asn94, Arg100 |
| TF | -8.7 | Asn39, Ser40, Ala43, Lys46, Met86, Asn94, Arg100, Phe126 |
| TF3G | -9.1 | Glu35, Ser38, Asn39, Ser40, Asp42, Ala43, Lys46, Gly85, Met86, Asn94, Arg100, Gly120, Gly125 |
| TF3'G | -8.8 | Asn39, Asp42, Ala43, Lys46, Gly85, Met86, Asn94, Arg100, Val124, Phe126 |
| TFDG | -8.7 | Asn39, Ser38, Asp42, Ala43, Asn94, Leu95, Arg100, Gly125 |

**Supplementary Fig. 10:** *In-silico* analysis revealed a better affinity and binding of Theaflavins to Arabidopsis HSP90.

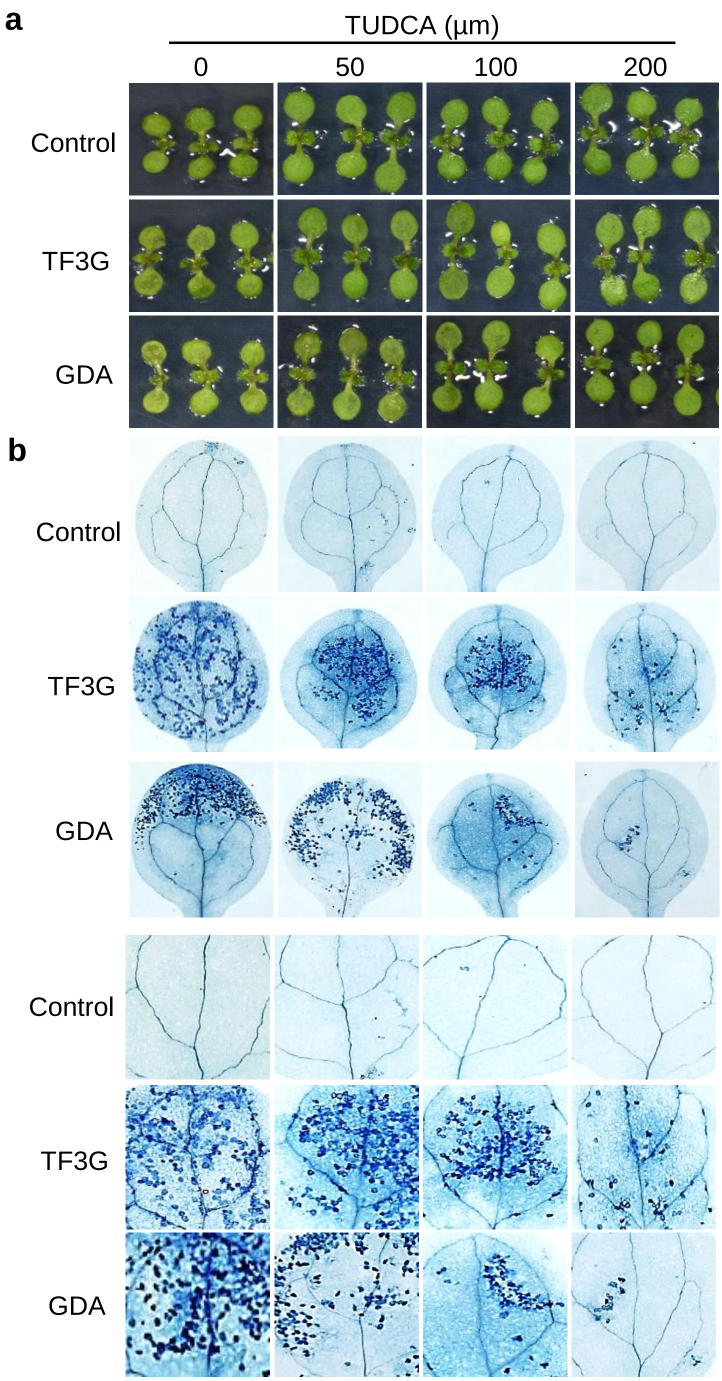


**Supplementary Figure. 11** Synthetic chaperone TUDCA promotes protein folding and attenuates GDA and TF3G-induced ER stress and cell death.

**(a)** Plant phenotype.

**(b)** Cell death. Five-day-old WT Arabidopsis seedlings were vacuum infiltrated in equivolume solution containing TUDCA (50, 100 and 200 µm) and 200 µm TF3G and GDA for 5 min followed by 55 min incubation in the same solution. Treated seedlings were transferred to fresh liquid MS and shifted to growth chamber. Five seedlings from each treatment were harvested after 24 h, imaged, and stained with TB staining.

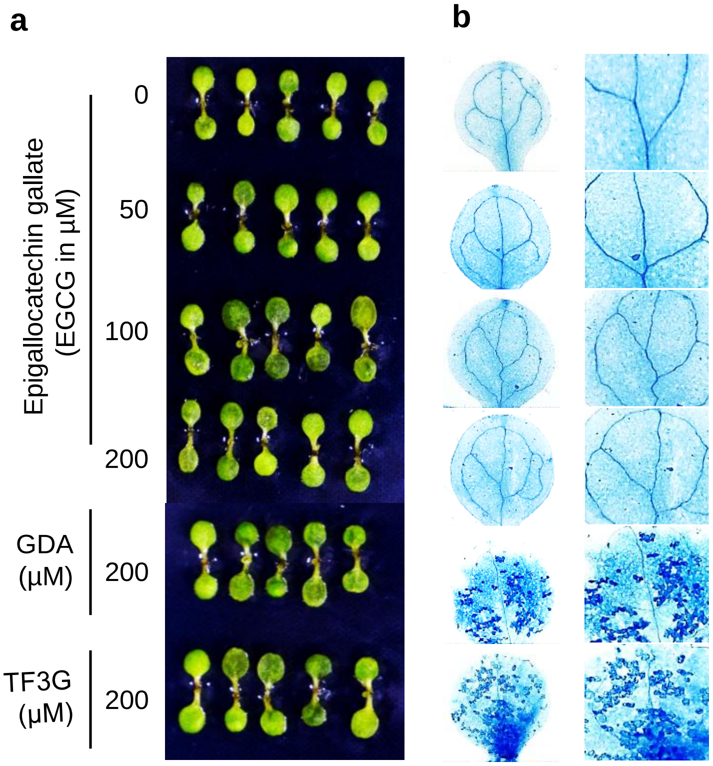


**Supplementary Fig. 12:** Epigallocatechin gallate (EGCG) does not induce cell death.

Five-day-old WT Arabidopsis seedlings were vacuum infiltrated with distilled water (as control) and EGCG at 50,100 and 200 µM for 5 min followed by 55 min incubation. After incubation, chemicals were replaced with liquid MS and the seedlings were transferred to growth chamber. After 24 , seedlings were harvested for phenotypes (**a**), and cell death detection using TB staining **(b)**.

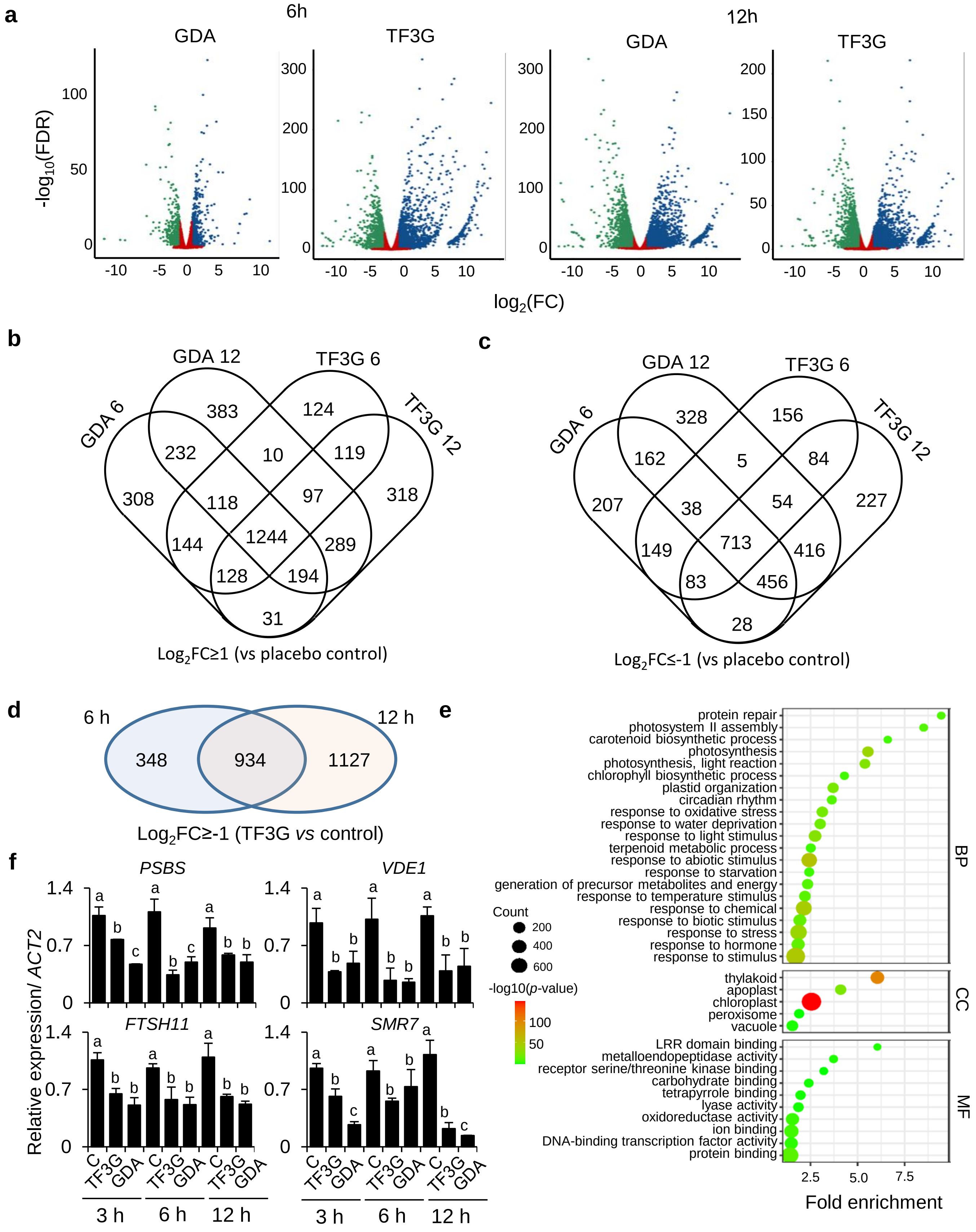
a
6h
12h
GDA
TF3G
-log10(FDR)
log2(FC)
b
GDA 6
GDA 12
TF3G 6
TF3G 12
Log2FC≥1 (vs placebo control)
c
Log2FC≤-1 (vs placebo control)
d
6 h
12 h
348
934
1127
Log2FC≥-1 (TF3G vs control)
e
protein repair
photosystem II assembly
carotenoid biosynthetic process
photosynthesis
photosynthesis, light reaction
chlorophyll biosynthetic process
plastid organization
circadian rhythm
response to oxidative stress
response to water deprivation
response to light stimulus
terpenoid metabolic process
response to abiotic stimulus
response to starvation
generation of precursor metabolites and energy
response to temperature stimulus
response to chemical
response to biotic stimulus
response to stress
response to hormone
response to stimulus
thylakoid
apoplast
chloroplast
peroxisome
vacuole
LRR domain binding
metalloendopeptidase activity
receptor serine/threonine kinase binding
carbohydrate binding
tetrapyrrole binding
lyase activity
oxidoreductase activity
ion binding
DNA-binding transcription factor activity
protein binding
BP
CC
MF
Count
-log10(p-value)
Fold enrichment
f
PSBS
VDE1
FTSH11
SMR7
Relative expression/ ACT2
C
TF3G
GDA
3 h
6 h
12 h

**Supplementary Fig. 13:** Transcriptional signature of TF3G. (Supports Figure 8)

**a-c** Transcriptional changes induced by GDA and TF3G. Volcano plots **(a)** showing the transcriptional changes upon 6- and 12 h of GDA and TF3G treatments. Venn diagrams showing upregulated **(b)** and downregulated **(c)** genes. **d** Venn diagram showing distribution and overlapping of downregulated genes upon TF3G treatments. Log2FC: log2(fold change). **e** Enrichment bubble plot showing gene ontology enrichment of genes downregulated at 6- and 12 h of TF3G treatment. **f** Relative transcript levels of key downregulated genes involved in photosynthesis and cell cycle regulation. Gene expression was determined using qRT-PCR. Total RNA were isolated from 5-day-old seedlings after 3-, 6- and 12 h of treatment. *ACT2* was used as an internal standard. Values represent means ± SD of three independent biological replicates. Lowercase letters indicate statistically significant differences between mean values at each genotype ($P < 0.05$, one-way ANOVA with Tukey's post hoc honestly significant difference (HSD) test.

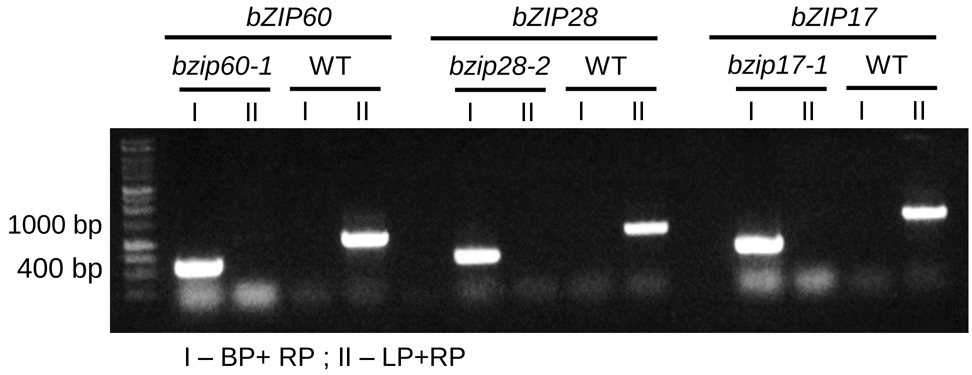


**Supplementary Fig. 14:** PCR-based genotyping of *bzip60-1*, *bzip28-2,* and *bzip17-1* mutants.

DNA was isolated from the pool of 5-day-old seedlings grown on MS media. PCR was carried out using gene-specific (LP and RP) and T-DNA-specific primers (BP) in a combination BP+RP and LP+RP. The homozygous recessive mutant shows amplification of ~400 bp with BP+RP and no amplification with LP+RP. The WT control shows an amplification of ~1000bp with LP+RP and no amplifications with BP+RP.

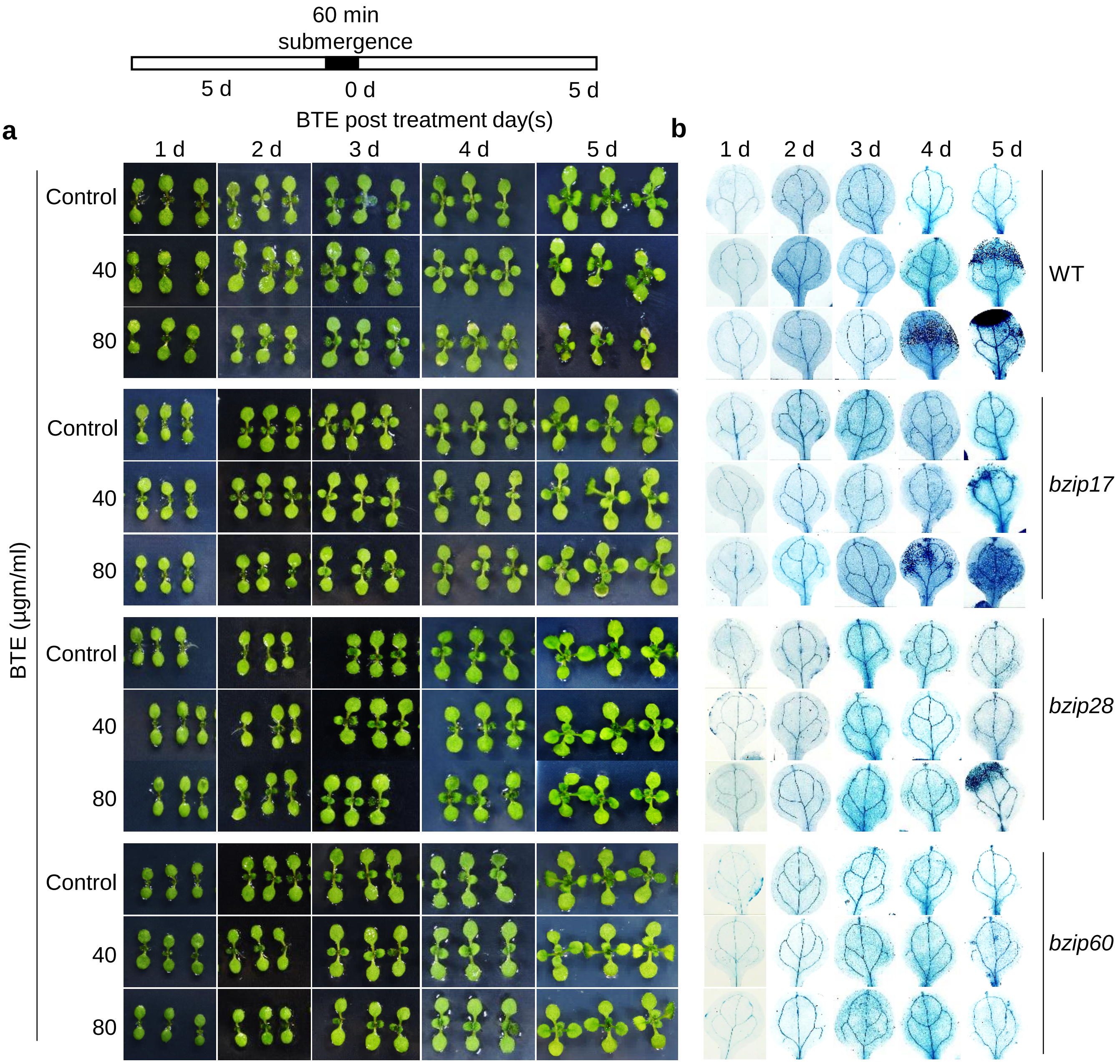


**Supplementary Fig. 15:** Black Tea extract (BTE) induced growth inhibition and cell death is mediated by bZIP60.

**a** Phenotypes of 5-day-old WT seedlings treated with BTE at 40 and 80 µg/ml. Seedlings treated with autoclaved distilled water served as controls. **b** Cell death. Five seedlings used for phenotype were stained with TB.

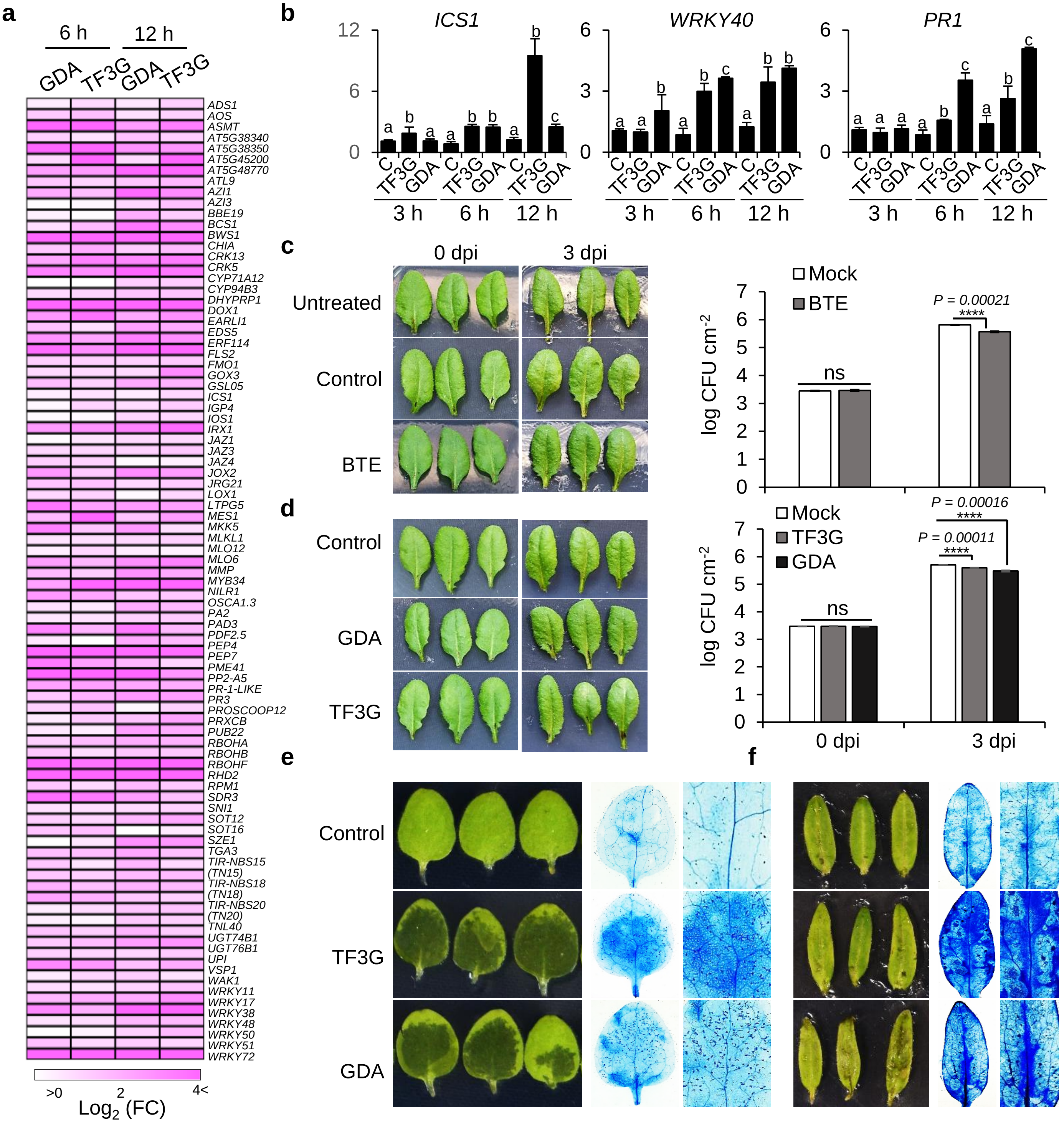
a
6 h
12 h
GDA
TF3G
GDA
TF3G
ADS1
AOS
ASMT
AT5G38340
AT5G38350
AT5G45200
AT5G48770
ATL9
AZI1
AZI3
BBE19
BCS1
BWS1
CHIA
CRK13
CRK5
CYP71A12
CYP94B3
DHYPRP1
DOX1
EARLI1
EDS5
ERF114
FLS2
FMO1
GOX3
GSL05
ICS1
IGP4
IOS1
IRX1
JAZ1
JAZ3
JAZ4
JOX2
JRG21
LOX1
LTPG5
MES1
MKK5
MLKL1
MLO12
MLO6
MMP
MYB34
NILR1
OSCA1.3
PA2
PAD3
PDF2.5
PEP4
PEP7
PME41
PP2-A5
PR-1-LIKE
PR3
PROSCOOP12
PRXCB
PUB22
RBOHA
RBOHB
RBOHF
RHD2
RPM1
SDR3
SNI1
SOT12
SOT16
SZE1
TGA3
TIR-NBS15 (TN15)
TIR-NBS18 (TN18)
TIR-NBS20 (TN20)
TNL40
UGT74B1
UGT76B1
UPI
VSP1
WAK1
WRKY11
WRKY17
WRKY38
WRKY48
WRKY50
WRKY51
WRKY72
>0
2
4<
Log2 (FC)
b
ICS1
WRKY40
PR1
C
TF3G
GDA
3 h
6 h
12 h
c
0 dpi
3 dpi
Untreated
Control
BTE
Mock
BTE
P = 0.00021
****
ns
log CFU cm-2
d
Control
GDA
TF3G
Mock
TF3G
GDA
P = 0.00016
P = 0.00011
****
ns
log CFU cm-2
0 dpi
3 dpi
e
f
Control
TF3G
GDA

**Supplementary Fig. 16:** TF3G is a general elicitor of hypersensitive response-like cell death and activates basal immunity in plants.

**a** Heatmap showing the expression of defense-related genes induced upon 6- and 12 h of GDA and TF3G treatments. Colour scheme shows Z-score with upregulated genes in pink and downregulated in blue. **b** Relative transcript levels of key basal genes involved in basal immunity. Transcript expression of *ICS1*, *WRKY40* and *PR1* was determined using qRT-PCR. *ACT2* was used as an internal standard. Values represent means ± SD of three independent biological replicates. Lowercase letters indicate statistically significant differences between mean values at each genotype ($P < 0.05$, one-way ANOVA with Tukey's post hoc honestly significant difference (HSD) test). **c-d** Effects of Black Tea Extract (BTE) (**c**) , TF3G and GDA (**d**) on the growth of *Pseudomonas syringae* pv. tomato (*Pst*) DC3000. *Pst* DC3000 were infiltrated into 4-week-old Arabidopsis leaves pretreated with $H_2O$ (as mock), and BTE (100 µgm/ml) (**c**) and DMSO (as mock), TF3G and GDA (200µM) (**d**) for 24 h. Bacterial colonies were counted at 0 dpi (to indicate uniform bacterial infection) and 3 dpi (to evaluate treatment effect) at serial dilutions of $10^{-3}$,$10^{-4}$ and $10^{-5}$. Bacteria were extracted from three different leaves of four independent plants and incubated at 28°C for 2 d to evaluate growth. Student's t-test was performed to calculate pairwise statistical significance in comparison to mock-treated plants; **** = $P$<0.0001; ns = nonsignificant. Experiments were repeated thrice with similar results. CFU, colony-forming units. **e-f** TF3G and GDA induced cell death in *Nicotiana benthamiana* **(e)** and Tomato seedlings **(f)**. Five-day-old seedlings grown in continuous light conditions were vacuum-infiltrated with GDA and TF3G (200 µM) along with 0.1% Tween-20 for 5 min followed by 55 min incubation. After incubation, the solution were replaced with liquid MS, and the seedlings were transferred to growth chamber. After 24 h, seedlings were harvested for phenotype and cell death.
